# Rational design and comparative docking and simulation of modified FLT3 inhibitors: A study on enhanced binding stability and inhibition potency

**DOI:** 10.1101/2025.03.12.642924

**Authors:** Uddalak Das, Dheemanth Regati, R. Sowdhamini, Jitendra Kumar

## Abstract

The FLT3 protein is a well-established therapeutic target in the treatment of Acute Myeloid Leukemia (AML), with its inhibition playing a crucial role in disease management. In this study, we identify and propose a novel FLT3 inhibitor that demonstrates superior binding affinity, stability, and pharmacokinetic properties compared to currently available inhibitors. We initially characterized the binding interactions of known FLT3 inhibitors through molecular docking and then strategically modified functional groups to enhance binding affinity, optimize drug-likeness, and minimize toxicity. The resulting analogue exhibits improved metabolic stability, lower toxicity, higher intestinal absorption, and superior permeability. Molecular dynamics simulations further confirm that the novel inhibitor forms stable and persistent interactions with FLT3, as evidenced by reduced conformational fluctuations and compact structural integrity. Free energy calculations reveal stronger ligand stabilization, while dynamic correlation analysis suggests enhanced engagement with critical residues, reinforcing its potential as an effective therapeutic agent. These findings highlight a promising candidate for further experimental validation and potential development in AML treatment.

## 1. INTRODUCTION

Acute myeloid leukemia (AML) stands as a formidable hematologic malignancy making up 80% of all adult leukemia cases [1]. According to recent statistics, there were approximately 20,380 new cases of Acute Myeloid Leukemia (AML) and 11,310 associated deaths in the United States in the year 2023 [2]. These figures underscore the continued significance of addressing and advancing research, diagnosis, and treatment options for AML to improve patient outcomes and reduce mortality rates. Originating from cytogenetic aberrations within bone marrow hematopoietic cells, AML manifests as a clonal transformation of immature myeloblasts, impacting blood cell formation and differentiation [3]. The primary focus of therapeutic investigations has honed in on the intricate molecular landscape of AML, with specific attention directed towards the family of receptor tyrosine kinase FMS-like tyrosine kinase 3 (FLT3).

FLT3, a class III protein tyrosine kinase, orchestrates essential signalling cascades critical for hematopoietic regulation [4]. The receptor, composed of five immunoglobulin-like extracellular domains, a transmembrane domain, and two intracellular tyrosine kinase domains, plays a crucial role in immunological responses and hematopoietic stem cell proliferation [5]. Mutations within the FMS-like tyrosine kinase 3 (FLT3) gene manifest in around 30% of AML instances, with the internal tandem duplication (ITD) constituting the predominant subtype (FLT3-ITD; approximately 25% of AML cases). Additionally, mutations within the tyrosine kinase domain (FLT3-TKD) are observed in a subset of AML cases, accounting for approximately 7-10% of occurrences. These mutations impart constitutive activation to FLT3, resulting in uncontrolled proliferation of leukemic cells [6].

The emergence of FLT3 mutations is identified as a critical driver in the pathogenesis of AML, elucidating the importance of FLT3 as a therapeutic target [7]. The classification of FLT3 inhibitors into first and next generations, based on potency and specificity, signifies a strategic evolution in drug design [8]. The landscape of FLT3 inhibitors encompasses both type I and type II inhibitors, with the latter displaying distinct advantages in targeting the hydrophobic pocket adjacent to the ATP-binding site in the inactive DFG-out conformation [9].

Clinical trials involving the first generation FLT3 inhibitors demonstrated limited efficacy due to issues of kinase selectivity [10]. The second generation, containing potent inhibitors like *Gilteritinib* and *Quizartinib*, offered heightened specificity and selectivity [11]. Despite advancements in the field, the rapid emergence of resistance, coupled with the inadequate binding affinity, instability within the active site, and insufficient potency of several experimentally validated FLT3 ligands identified as potential FLT3 inhibitors, has hindered their clinical success [12].

*In silico* drug discovery has revolutionized therapeutics by overcoming the limitations of traditional experimental methods [13]. Promising lead identification conventionally involves costly and time-consuming high-throughput screening (HTS) [14]. The drug discovery process, taking up to 14 years and costing around $800 million, faces challenges due to failures in clinical trials [15–17]. Rational drug design, or reverse pharmacology, is a cost-effective approach that begins with identifying target proteins and screening small-molecule libraries [18]. Structure-Based Drug Design optimizes lead discovery by utilizing target protein 3D structures and disease insights, employing techniques such as structure-based virtual screening (SBVS), molecular docking, and molecular dynamics (MD) simulations [19]. Due to the complexities of target identification, drug design, and screening in AML drug development, computational methods are essential.

In this study, we focuses on the rational design and optimization of the existing inhibitors. By modulating the top-performing compounds, we aim to enhance their binding affinity to key active-site residues, improve their stability, and ultimately bridge the gap between experimental validation and clinical efficacy, paving the way for more effective FLT3-targeted therapies in AML.

## 2. MATERIALS AND METHODS

This study aims to optimize existing FLT3 inhibitors for AML by enhancing binding affinity, stability, and clinical efficacy, with the overall workflow illustrated in **Figure 1**.

**Figure 1.**
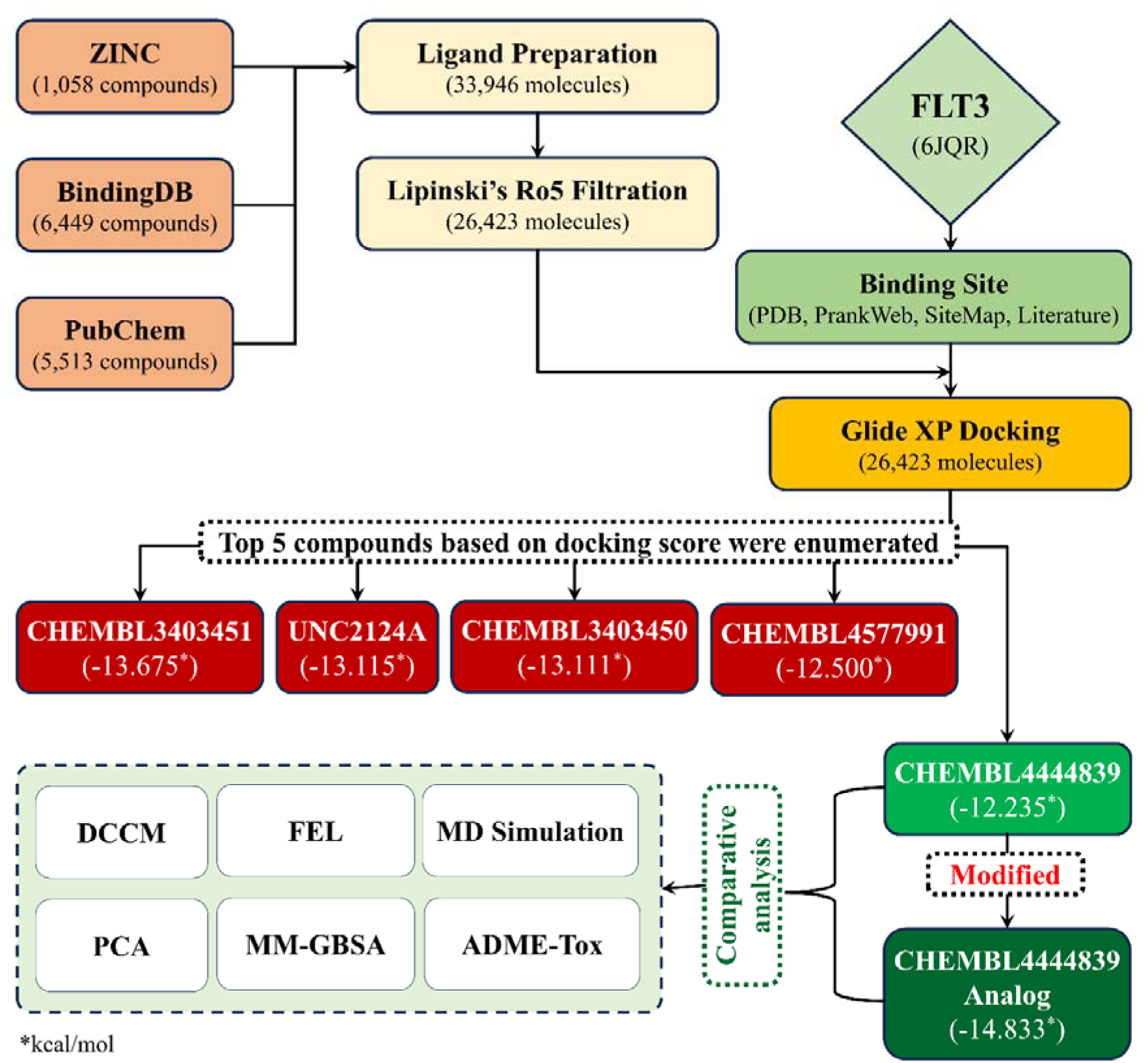
A flowchart illustrating the workflow employed in this study.

### 2.1. Computational systems

A CentOS Linux 7 workstation with 31.1 GiB memory, Intel ® Xeon ® E-2234 CPU with 3.60 GHz x 8 processor, and Quadro p1000/PCle/SSE2 graphics was used. Schrödinger release 2023-1 was used for all computational purposes.

### 2.2. Dataset generation and screening

For the screening, all compounds exhibiting inhibitory activity against FLT3 were selected from the ZINC [20], BindingDB [21], and PubChem databases, comprising 1,050, 6,449, and 5,513 compounds respectively. These compounds were prepared using LigPrep [22], resulting in a total of 33,946 molecules. The ligands were then screened based on Lipinski’s Rule of Five (Ro5) [23] using the QikProp module of Schrodinger. 26,423 compounds showing zero Ro5 violations were kept for further studies.

### 2.3. Molecular Interactions

#### 2.3.1. Protein preparation

The X-ray crystallographic structure of the FLT3 target protein [PDB ID: 6JQR], in complex with *Gilteritinib* [24] was used in the study.

In the Schrödinger Maestro protein preparation wizard, the protein was pre-processed with the PROPKA module for an optimization of H-bonds [25], followed by minimization of structures towards convergence of heavy atoms at RMSD 0.3Å using OPLS4 force field and removal of water molecules more than 5Å away from ligands afterward [26].

#### 2.3.2. Receptor grid generation

The receptor grid was generated keeping the hydrophobic region and also the region where *Gilteritinib* was attached to the complex, at the centroid of the grid. The coordinates of the receptor grid were X=−30.02, Y=−11.86, Z=−26.27, with ligand size up to 18A.

#### 2.3.3. Molecular docking

Docking was limited to ligands with 100 rotatable bonds and fewer than 500 atoms. Van der Waals radii scaling factor was set to 0.80, with a partial charge cutoff of 0.15. Sample nitrogen inversions and sample ring conformations were activated, and the ligand sampling was set to flexible. All predicted functional groups had bias sampling of torsions enabled. The module was configured to promote intramolecular hydrogen bonds and improve conjugated pi groups’ planarity. Extra precision (XP) docking [27] was done using the Glide module of Schrödinger. Top 5 compounds showing the highest docking score were considered for enumerations.

#### 2.3.4. Ligand enumerations

All the functional groups in the ligand that showed proximity to the protein were altered to 2383 other functional groups using the ligand designer [28] and R-group enumeration [29] modules of the Schrodinger Suite.

### 2.4. ADME-Tox analysis

A comparative pharmacokinetic analysis, specifically examining the ADMET profiles of the canonical ligand and its modification, identified from the ligand R-group enumeration, was conducted using several tools: Schrödinger’s QikProp module, SwissADME [30], ProTox-3.0 [31], ADMETlab 3.0 [32], and the pkCSM server [33]. SwissADME, pkCSM, and ProTox-3.0 are web-based tools designed to predict pharmacokinetic properties, assess drug-likeness, and evaluate the medicinal chemistry suitability of small molecules.

### 2.5. Molecular dynamics simulation studies

Desmond package was used to carry out the molecular dynamics simulations for the complex of FLT3 protein with the canonical ligand and its modification. Each system was placed individually in an orthorhombic water box of 10 Å using the TIP3P water model [34]. The ligand-protein complexes were modelled by the OPLS4 force field [35]. Counter ions (Na+) were introduced in the ligand-protein complex structures to neutralize the total charge of the systems undergoing MD simulation. Furthermore, the energies of the systems were minimized to a minimum level using 2000 steps before initiating the MD simulation along an NPT lattice trajectory [36].

After equilibration, a production run was executed at a temperature of 310.15 K and a pressure of 1 atm for 500 ns to evaluate the stability of the system in an aqueous environment. Several structural and stability analyses were conducted on the resulting molecular dynamics trajectories, such as the Root mean square deviation (RMSD), Root mean square Fluctuation (RMSF) and Radius of Gyration (Rg). To evaluate the protein’s exposure to the solvent throughout the simulation, the solvent-accessible surface area (SASA) was calculated. The graphical representations were generated using Matplotlib Library 3.10.0 [40] in using in-house python scripts.

### 2.6. Binding free energy distribution

The VMD analysis scripts were used to visualize trajectory files of the MD simulation. For the analysis of principal components and the evaluation of the free energy landscape (FEL), the covariance matrix was computed and subsequently diagonalized. FEL and MM-GBSA was employed to determine the stable and static interactions.

#### 2.6.1. Free energy landscape (FEL)

The FEL was formulated as: *F*(*X*) = *k_B_TlnP*(*X*), where *F(X)* represents the free energy, *k_B_* is the Boltzmann constant, *T* is the absolute temperature, and *P(X)* represents the probability distribution of the molecular system along the principal component.

The 2D and 3D FEL were constructed to analyze conformational transitions using RMSD, radius of gyration (Rg), and other parameters. For FEL calculations, RMSD and Rg values were projected onto a plane, with population densities estimated via kernel density estimation, and visualized as contour maps in Python [40].

#### 2.6.2. Molecular Mechanics-Generalized Born Surface Area

The prime module was utilized to predict the energy parameters obtained from the MM-GBSA simulation to predict the amounts of the stabilization energy coming from the potential interaction. The VSGB solvation model [41] was used and the force field was set as OPLS4. The MM-GBSA-based binding free energy calculations were done on the 500ns long MDS trajectories. For the selection of protein-ligand complexes, the binding energies calculated by this approach are more efficient than the glide score values. The main energy elements like H-bond interaction energy (DG Bind_Hbond), electrostatic solvation free energy (DG Bind_Solv), Coulomb or electrostatics interaction energy (DG Bind_Coul), lipophilic interaction energy (DG Bind_Lipo), and van der Waals interaction energy (DG Bind_vdW) altogether were considered to the calculation of MM-GBSA based relative binding affinity.

## 3. RESULTS

### 3.1. The FLT3 protein

Human FLT3 has a length of 993 amino acids and weighs 112.903 Da, having three components – the extracellular domain with 516 residues, the transmembrane domain with 19 amino acids, and the cytoplasmic domain of 429 amino acids [42]. To determine the structure of 6JQR which is in the cytoplasmic domain, the PDBsum web server was utilized [43]; this analysis identified fifteen α-helices, six β-hairpins, four β-bulges, as well as 12 helix-helix interactions (see **Figure 2**).

**Figure 2.**
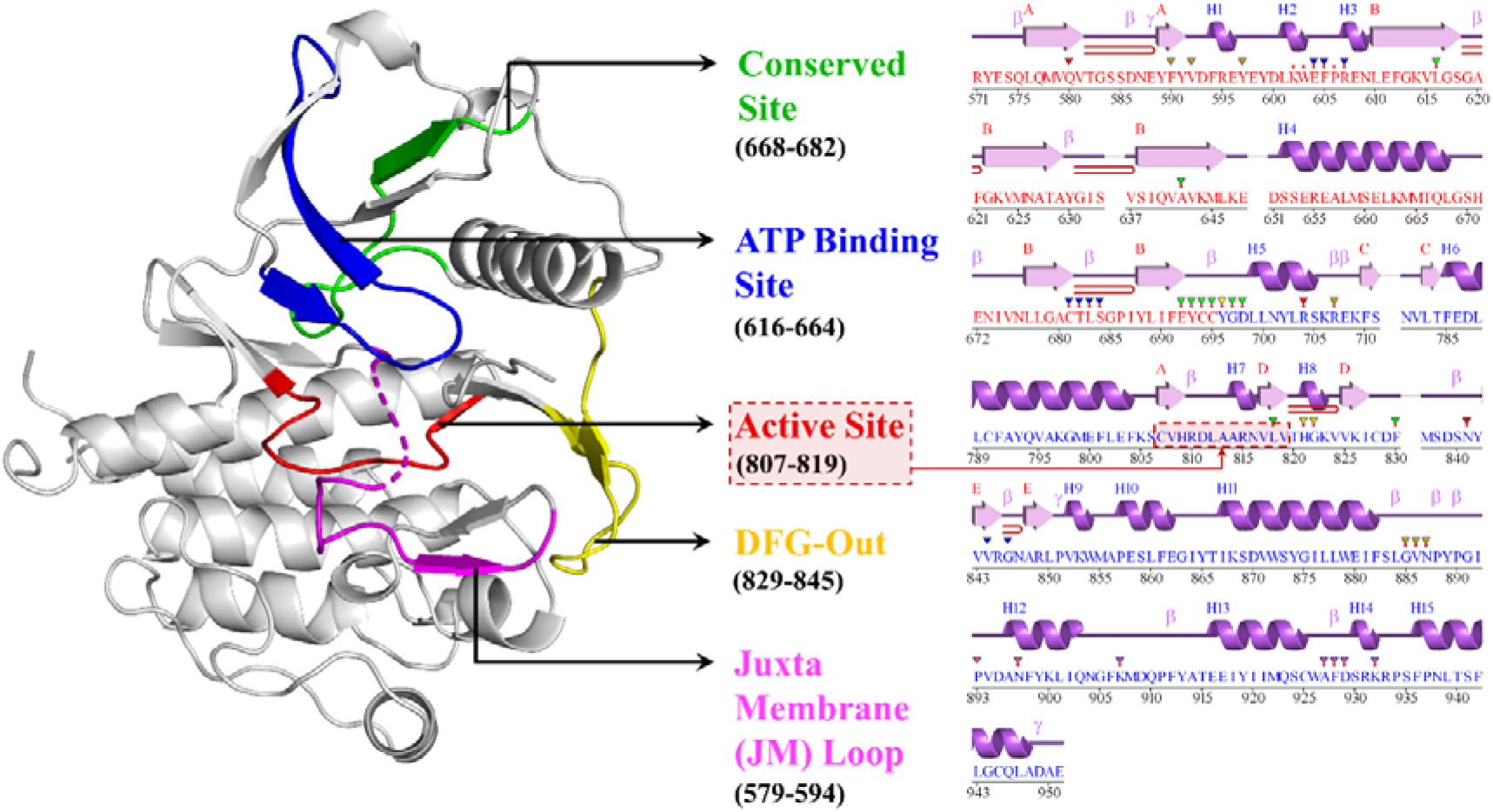
Crystal structure of FLT3 protein kinase domain (residues 610–943): The conserved site (green) highlights a region crucial for protein function and interactions. The ATP binding site (blue) signifies the location where ATP molecules bind to initiate kinase activity. The active site (red) is where phosphorylation events occur, leading to downstream signaling. The DFG-Out motif (purple) plays a role in regulating kinase activity. The juxtamembrane (JM) loop (magenta) is involved in protein localization and interactions.

The Ramachandran plot was also used to validate 6JQR, and it indicated that 93.5% of the residues were in preferred regions, 6.5% were in additional residue regions, none in generous regions, and 0.9% in disallowed regions with a total G-Factor of 0.00 [44]. (see **Supplementary Figure S1**)

### 3.2. Molecular docking studies

The docking assessments for the top 5 compounds, i.e., CHEMBL3403451 [45], UNC2124A (Patent ID: US10004755B2), CHEMBL3403450 [46], CHEMBL4577991 [45] and CHEMBL4444839 [45] yielded scores between −13.675 kcal/mol and −12.500 kcal/mol as a result of the XP docking procedure. These compounds were selected for modification for enhanced binding affinity and stability. The 2D and 3D interaction diagram of these ligands within the FLT3 binding site (**Supplementary Figure S2**).

### 3.3. Ligand enumerations

We performed ligand enumeration of all the top 5 docked compounds from the XP docking. We set a parameter of a Δ docking score of 2 kcal/mol from the original docking score of the canonical compounds. The first four compounds did not produce any analogue with enhanced binding affinity compared to their canonical versions. However, CHEMBL4444839 generated multiple analogues with a Δ docking score of 2 kcal/mol. The **best-scoring analogue** of CHEMBL4444839 exhibited a key modification—the incorporation of a fluorocyclobutane moiety **(Figure 3)**. See **Supplementary Table S4** for 2D and 3D structures of CHEMBL4444839 and its analogue. Interestingly, the binding affinity of the best CHEMBL4444839 analogue exhibited an enhanced docking score of more than 1 kcal/mol compared to the best compound, CHEMBL3403451 (−13.675 kcal/mol). CHEMBL3403451 and its analogue were evaluated for their comparative pharmacokinetic and other properties. See **Supplementary Figure S5** for the ligand torsions plot summarizing the conformational evolution of every rotatable bond in the ligands, providing insights into their flexibility and binding adaptability.

**Figure 3.**
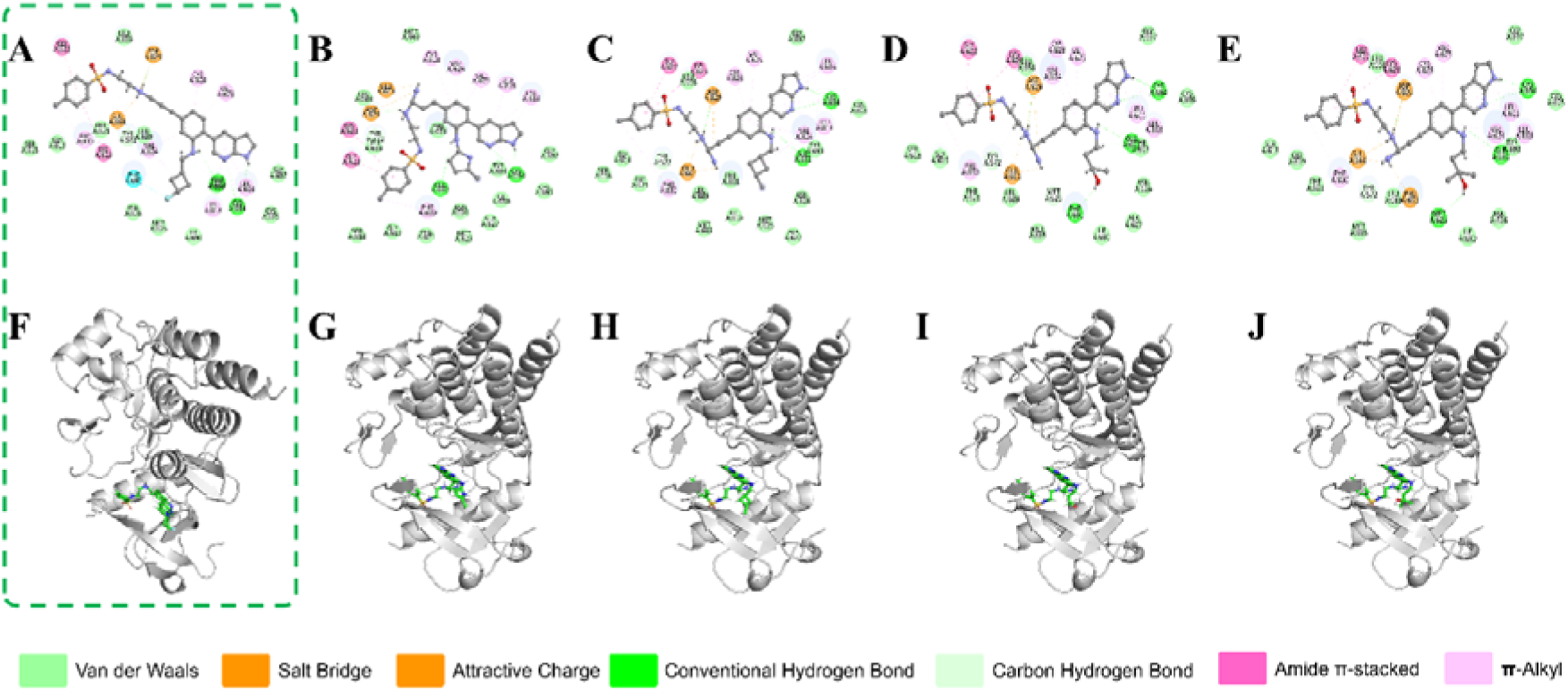
The top 5 enumerations of the ligand CHEMBL4444839. The panels A-E shows the 2D interactions and the panels F-J shows the 3D interactions with FLT3 protein active site, with docking scores of −14. 833, −14.832, −14.743, −14.629, and −14. 629 kcal/mol respectively. The compound in green dotted square shows the best enumeration with the highest docking score.

### 3.4. ADME-Tox predictions

In this study, we utilized SwissADME, ADMETlab 2.0, ProTox-II and pkCMS web servers to predict the ADMET properties of the CHEMBL3403451 and its analogue (**Figure 4**).

**Figure 4.**
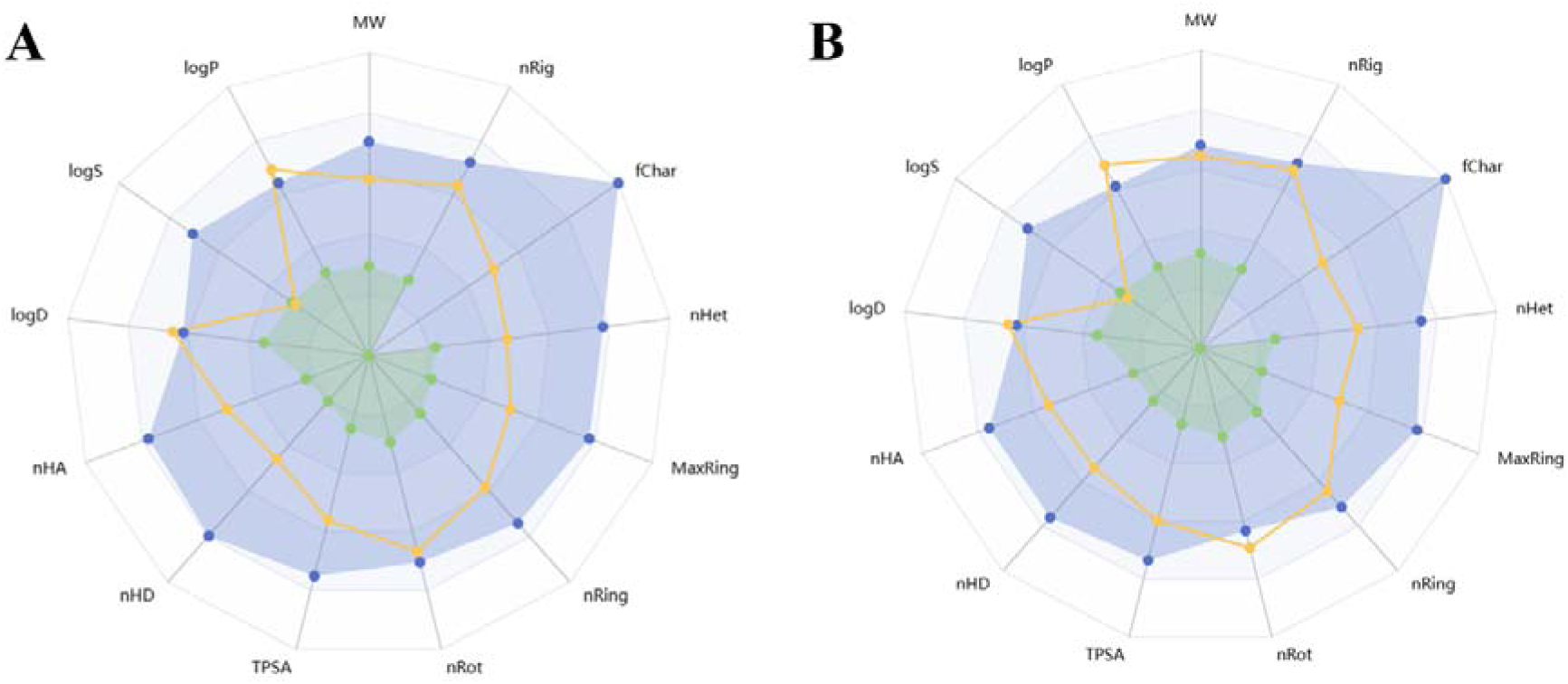
The Bioavailability Radar Plot illustrating the comparative drug-likeness of the two compounds. The blue area indicates the ideal range for each property, while the yellow dots represent the properties of the compounds. A: CHEMBL4444839 and B: CHEMBL4444839-analogue.

With a toxicity class of 6 compared to class 4 for CHEMBL4444839, the analogue exhibits lower acute toxicity, supported by a much higher predicted LD50 of 5500 mg/kg versus 420 mg/kg. Unlike CHEMBL4444839, which is AMES toxic, the analogue is not, reducing its mutagenicity risk. The analogue also shows better systemic tolerance, with a higher maximum tolerated dose (0.496 vs. 0.4). While both compounds exhibit similar absorption profiles, the analogue demonstrates slightly improved Caco-2 permeability (0.55 nm/sec vs. 0.539 nm/sec) and human intestinal absorption (78.041% vs. 77.745%). The analogue excels in distribution characteristics, with a higher volume of distribution (1.324 log L/kg vs. 1.053) and lower plasma protein binding (93% vs. 96.5%), enhancing its bioavailability and pharmacodynamic activity. The metabolic profiles differ, with the analogue being a CYP2D6 substrate, suggesting a broader metabolic pathway, while both compounds interact similarly with key CYP enzymes. The analogue has a marginally higher total clearance rate (1.277 log ml/min/kg vs. 1.223), indicating faster elimination. Importantly, CHEMBL4444839-Analogue has reduced toxicity risks, including the absence of AMES toxicity, lower Pyriformis toxicity (µg/L = 0.285 vs. 0.286), and lower minnow toxicity (log mM = −0.102 vs. 0.656), suggesting better environmental safety.

**Supplementary Tables S1, S2, and S3** offers comprehensive comparative pharmacokinetic and toxicity assessments of CHEMBL4444839 and its analogue.

### 3.5. Molecular Dynamics Simulation

The MD trajectories were compared in terms of their RMSD, RMSF, Radius of Gyration, the protein ligand contacts, and the SASA values (**Figure 5**). CHEMBL4444839 appears to allow greater fluctuations in RMSD of the protein compared to the analogue bound structure, suggesting the increased stability of the protein when complexed with the analogue (Panel A).

**Figure 5.**
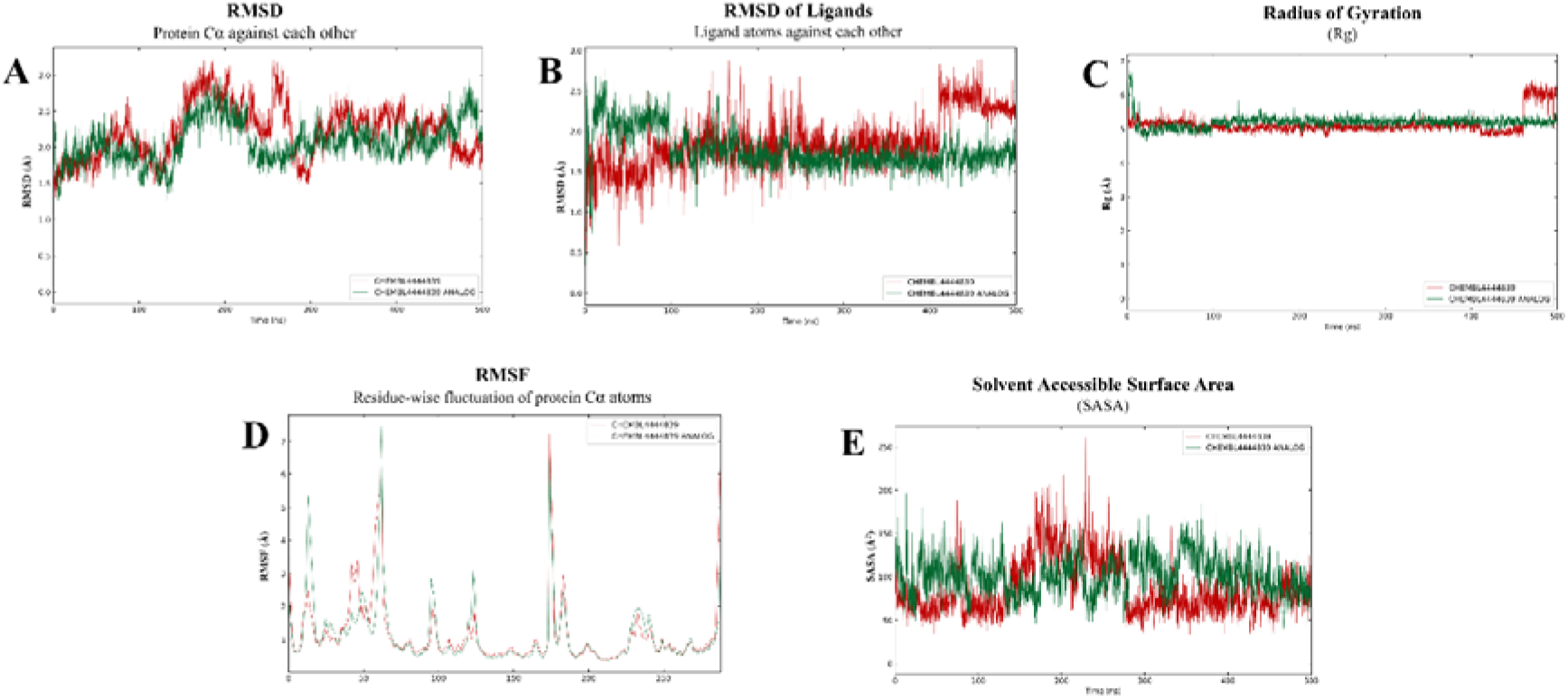
The comparative trajectory assessment of FLT3 backbone and ligand complex during a 500 ns trajectory. **(A)** RMSD of protein, **(B)** RMSD of ligand, (**C)** Rg, **(D)** RMSF, and **(E)** SASA.

The Residue Mean Square Fluctuation (RMSF) plot provides valuable insights into the dynamic flexibility of the FLT3 protein residues when complexed with two ligands: CHEMBL4444839 and its analogue. Certain regions which are responsible for the activity of FLT-3 such as Juxta Membrane (JM) loop (579-594), ATP binding site (616-664), the active site (807-819), and other conserved sites show reduction in the RMSF in the analogue bound structure, which appears to indicate that the analogue stabilizes the structure more efficiently compared to the original ligand (Panel D). **Supplementary Figure S4** shows the ligand RMSF analysis indicating positional fluctuations of ligand atoms, providing insights into their entropic contributions and interactions with FLT3.

The RMSF analysis of FLT3 regions reveals distinct ligand-induced stability patterns. In the JM loop (579-594), CHEMBL4444839-Analogue exhibits reduced fluctuations compared to CHEMBL4444839, indicating enhanced stabilization, crucial for FLT3 activation control [47]. At the ATP binding site (616-664), both ligands show low RMSF, but the analogue displays a slight reduction, suggesting stronger interactions with ATP-critical residues, enhancing inhibition. The conserved site (668-682) shows a fluctuation peak, but lower RMSF for the analogue implies stronger binding and structural integrity. In the active site (807-819), the analogue stabilizes the region more effectively, crucial for enzymatic inhibition. The DFG-out region (829-845) also shows reduced fluctuations with the analogue, suggesting enhanced stabilization of the inhibitory state (see **Figure 6**).

**Figure 6.**
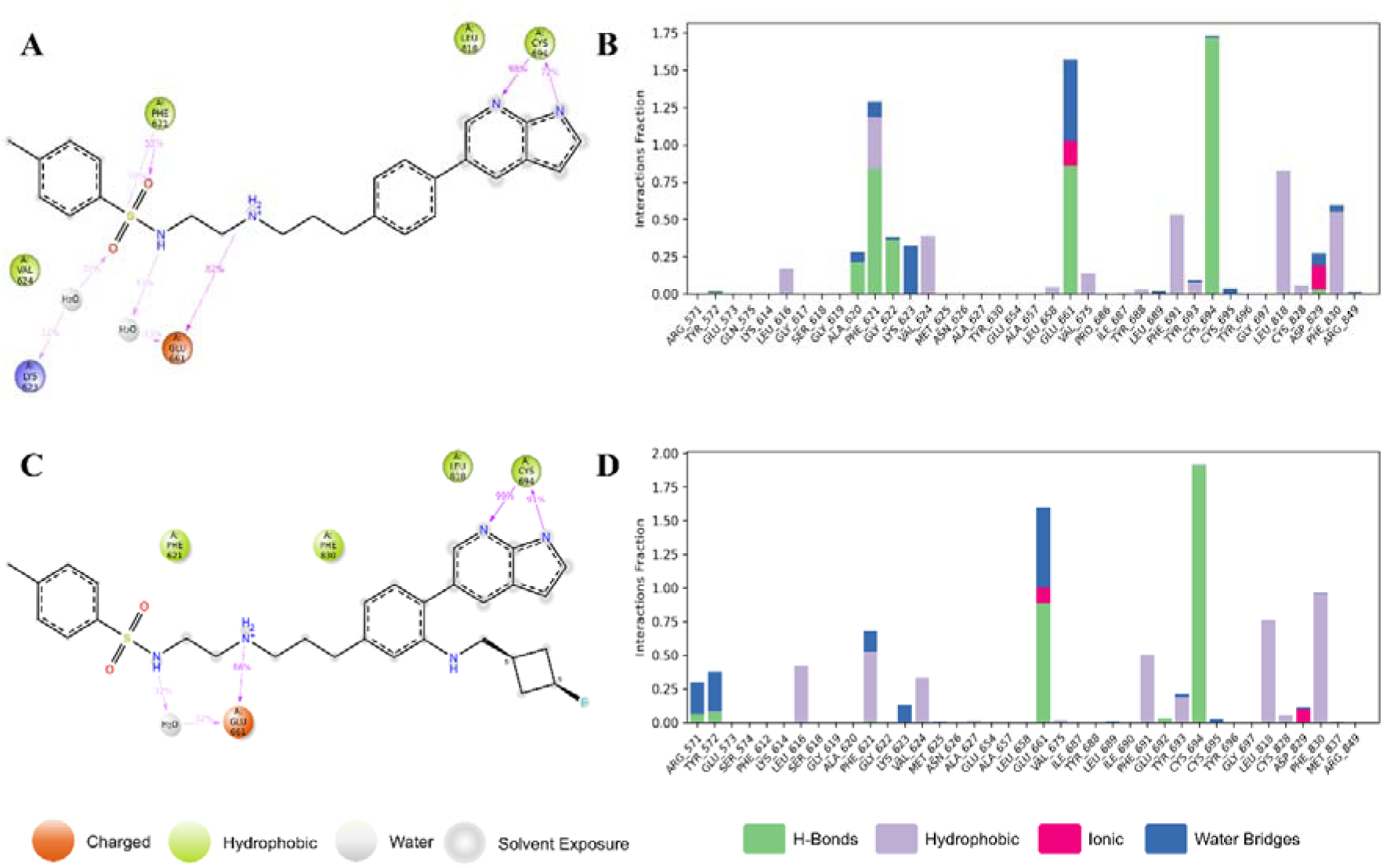
Molecular interactions of CHEMBL4444839 and its analogue in FLT3 binding pocket. Panels A and B depict the interactions for CHEMBL4444839, while Panels C and D illustrate those for the analogue.

High fluctuation peaks (~45, 125, 240 residues) indicate flexible regions, with the analogue moderating these movements, reinforcing overall protein stability. Interaction analysis (Panel A) shows CHEMBL4444839 forms strong H-bonds (GLU 661, 82%; CYS 694, 72%) and hydrophobic interactions (PHE 621, 55%; LEU 818, 98%), with moderate water-bridging contributions. Panel B highlights dominant hydrophobic interactions, particularly at LEU 818 and PHE 621, with minimal ionic interactions. The analogue exhibits increased interaction duration and novel contacts (e.g., PHE 830), suggesting superior FLT3 inhibition. The timeline representation of interactions in **Supplementary Figure S3** further illustrates the stability and duration of these contacts, revealing how the protein maintains ligand interactions across the simulation trajectory.

Towards the end of the simulation, radius of gyration (Rg) marginally increases for the structure bund with CHEMBL4444839, suggesting the stability of the protein could potentially be better with the analogue bound. Initially, CHEMBL4444839 exhibits a higher Rg (~6.5 Å) than the analogue (~6 Å), indicating a less compact starting state. Both systems stabilize by 20–30 ns, with the analogue maintaining a lower Rg (~5 Å) compared to CHEMBL4444839 (~5.2 Å) throughout 50–400 ns, suggesting enhanced structural integrity. The analogue’s Rg trajectory is smoother, implying reduced instability. In the final phase (400–500 ns), CHEMBL4444839’s Rg rises beyond 6 Å, indicating destabilization, while the analogue remains stable (~5 Å), suggesting superior long-term stabilization [48] (Panel C). To further assess the dynamic behavior of the ligand-protein complexes, we analyzed the variations in RMSD, rGyr, MolSA, intraHB, SASA, and PSA over 500 ns of MD simulation time (**Supplementary Figure S6**), which provide insights into the stability and compactness of the system.

SASA analysis shows dynamic ligand-protein interactions. CHEMBL4444839 maintains lower SASA values (50–150 Å²), with minimal spikes (<200 Å²), whereas the analogue exhibits higher SASA (100–200 Å²) with pronounced peaks, suggesting greater structural rearrangements. In 100–300 ns, the analogue shows significant fluctuations, reflecting dynamic binding, while CHEMBL4444839 remains more rigid. In 300–500 ns, the analogue sustains elevated SASA, indicating persistent interaction, whereas CHEMBL4444839 retains lower SASA, implying more restricted binding dynamics [49]. These findings highlight the analogue’s superior protein stabilization and dynamic binding properties, which may enhance efficacy (Panel E).

### 3.6. Binding free energy distributions

#### 3.6.1. Free energy landscape (FEL)

In **Figure 7**, Panels (A) and (C) represent CHEMBL4444839, while (B) and (D) depict its analogue. Each FEL plots free energy against RMSD and Rg. For CHEMBL4444839, the 2D FEL (A) shows a primary minimum at RMSD = 0.5-1.0 Å and Rg = 5.75-6.0 Å (~2.85 kcal/mol), stabilizing FLT3. A secondary basin at Rg ~6.2 Å suggests an alternative, less stable conformation. High-energy regions (RMSD > 2.0 Å) indicate destabilization. The 3D FEL (C) confirms a deep energy well at the primary minimum and sharp peaks for high RMSD, indicating conformational penalties. The energy range spans 2.85–5.28 kcal/mol, suggesting moderate flexibility. For the analogue, the 2D FEL (B) shows a minimum at RMSD = 0.5-1.0 Å and Rg = 6.0-6.25 Å (~2.61 kcal/mol), indicating better stabilization than the parent compound. A broader low-energy basin suggests higher conformational adaptability, with fewer high-energy states. The 3D FEL (D) reveals a deeper, smoother energy surface, supporting enhanced stability across a wider conformational space. The energy range (2.61–5.31 kcal/mol) highlights its superior ability to minimize unfavorable states.

**Figure 7.**
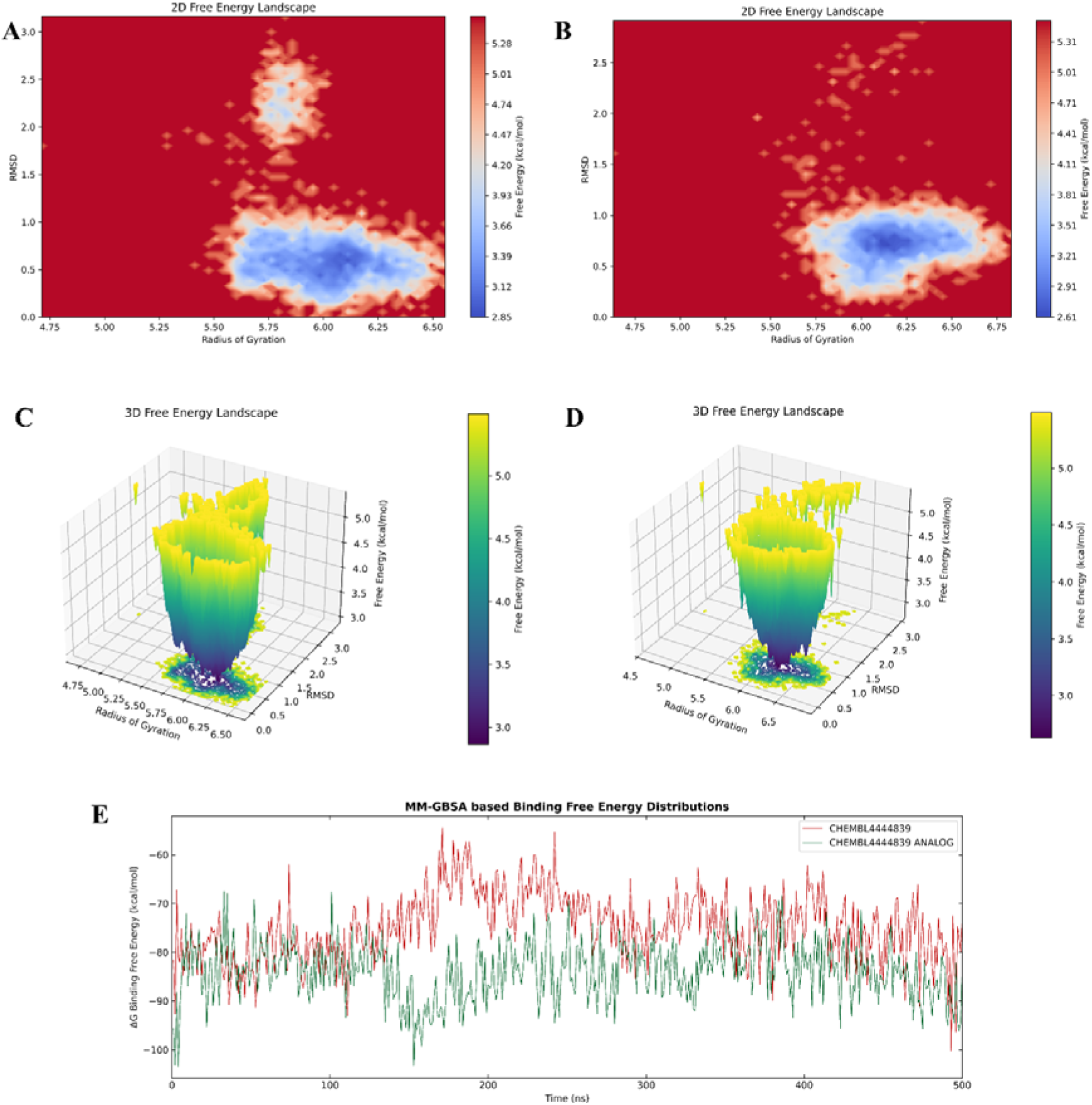
Comparative Free energy landscapes and binding free energy distributions of CHEMBL4444839 and its analogue. Panels A and B display 2D free energy landscapes projected onto RMSD and radius of gyration, revealing distinct conformational preferences. Panels C and D show 3D free energy landscapes, highlighting the energy minima associated with stable conformations. Panel E illustrates the MM-GBSA based binding free energy distributions over 500ns, demonstrating differences in binding affinity between the two molecules.

#### 3.6.2. Molecular Mechanics Generalized Born Surface Area (MM-GBSA)

The binding free energy distributions of FLT3 in complex with CHEMBL4444839 and its analogue were analysed over a molecular dynamics simulation (**Figure 7 (A)**). The ΔG values (kcal/mol) over time (ns) reveal distinct binding behaviours (**Supplementary Table S5 & S6**). CHEMBL4444839 (red line) exhibits greater fluctuations, with peaks exceeding −60 kcal/mol, indicating transient weaker binding. In contrast, the analogue (green line) maintains a more stable profile, rarely exceeding −70 kcal/mol, suggesting stronger, more consistent binding. CHEMBL4444839 shows a broader ΔG range (−100 to −60 kcal/mol), reflecting variability, whereas the analogue remains within −100 to −70 kcal/mol, indicating stability. Temporal analysis reveals erratic fluctuations for CHEMBL4444839, while the analogue stabilizes over time, suggesting durable interactions. The analogue’s lower average ΔG and smoother fluctuations indicate superior binding affinity and stability, likely due to structural modifications enhancing hydrogen bonding, hydrophobic interactions, or π-π stacking. These properties suggest its potential as a more reliable FLT3 inhibitor.

### 3.7. Trajectory analysis

#### 3.7.1. Principal component analysis (PCA)

The PCA plots (**Figure 8**) illustrate the conformational dynamics of FLT3 in complex with CHEMBL4444839 (Panel A) and its analogue (Panel B). Panel A exhibits greater dispersion, indicating higher conformational flexibility and structural variability, suggesting weaker and less stable binding. Conversely, Panel B shows tighter clustering, signifying a more stable complex with restricted conformational sampling. Temporal progression in Panel A is erratic, reflecting transient binding, while Panel B exhibits smoother transitions, indicating sustained stability. The broader distribution in Panel A suggests non-specific interactions, whereas Panel B’s confined range implies strong stabilization of FLT3 in a biologically relevant state.

**Figure 8.**
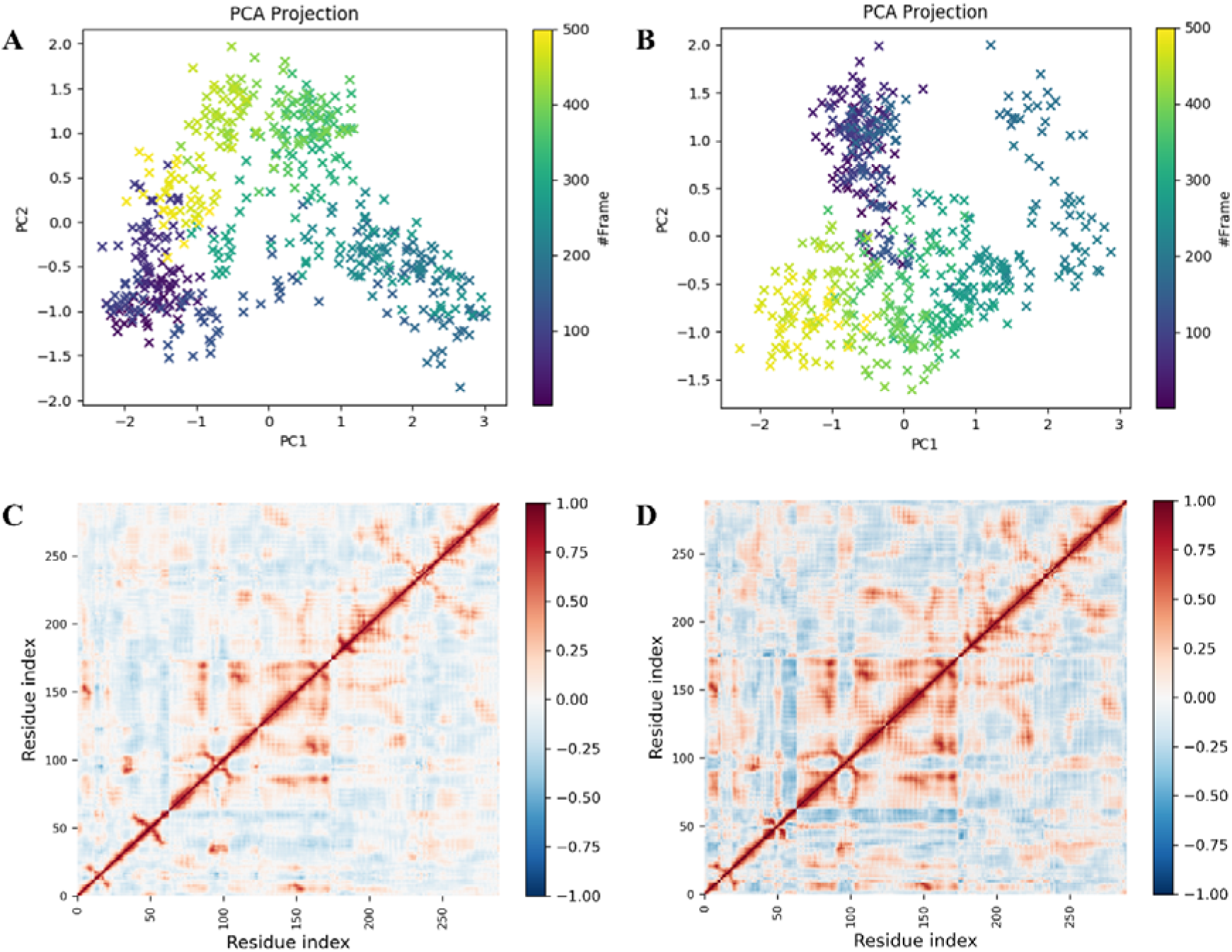
A-B: PCA projections of molecular dynamics trajectories, color-coded by frame number, showing conformational clustering along PC1 and PC2. C-D: Residue-residue contact maps highlighting dynamic correlations, with colour intensity indicating the strength of interactions over the simulation, revealing key structural changes.

#### 3.7.2. Dynamic cross-correlation matrix (DCCM)

In **Figure 8**, both CHEMBL4444839 (Panel C) and its analogue (Panel D) maintain self-correlation integrity, validating DCCM computation. However, Panel D exhibits a sharper diagonal, indicating more localized protein dynamics. Positive correlations in Panel C are weaker and dispersed (50–100, 150–200, ~250), while Panel D shows stronger, structured correlations (100–150, 200–250), enhancing FLT3 stability. Negative correlations are more extensive in Panel D, particularly between 50–100 and 150–200, suggesting enhanced dynamic adaptability. Panel C displays scattered correlations, whereas Panel D induces structured, functionally relevant allosteric effects. The analogue stabilizes FLT3 more effectively, promoting enhanced allosteric signaling and residue communication, particularly in the 50–100 and 150–200 regions, supporting its superior binding and dynamic impact over CHEMBL4444839.

## 4. DISCUSSION

The comparative analysis between CHEMBL4444839 and its analogue strongly suggests that the latter exhibits superior stability and binding characteristics with the FLT3 protein. The smoother and more consistent RMSD trends of the analogue indicate more stable interactions, potentially due to improved binding affinity and reduced conformational fluctuations. The analogue’s lower RMSF values across critical regions further support its superior stabilization and potential for enhanced inhibition.

Structurally, CHEMBL4444839 consists of a p-toluenesulfonyl (-SOL-) group attached to a secondary sulfonamide (-SOLNH-) functionality, which connects to a flexible alkyl linker (-NH-(CHL)L-NHL). This linker bridges the sulfonamide moiety to the 1H-pyrrolo[2,3-b]pyridine core, an aromatic bicyclic system that likely contributes to binding through π-π stacking and hydrogen bonding interactions. In the best analogue, a significant structural modification is the incorporation of a fluorocyclobutane moiety at position 5 of the pyrrolopyrimidine ring. This modification introduces a four-membered cyclobutane ring substituted with a fluorine atom, which increases steric bulk and imposes conformational rigidity on the heterocycle. The presence of fluorine imparts an electron-withdrawing effect, potentially altering dipole moments and influencing interactions with nearby residues. Additionally, the p-toluenesulfonyl (-SOL-) group and the flexible-(CHL)L-NHL alkyl linker remain unchanged, ensuring that key hydrogen bond donor-acceptor interactions in the parent compound are preserved.

The interaction profile of the analogue reveals increased binding stability through additional interactions with key residues, such as PHE 830 [24], GLU 661 [50], and CYS 694 [51], and optimized hydrophobic interactions with LEU 818 [52]. The analogue also maintains consistent water-bridging interactions, ensuring hydration dynamics remain intact while enhancing overall binding efficacy [53]. The lower Rg values for the analogue indicate a more compact and tightly bound complex, further emphasizing its superior stability over the original compound.

Fluorocyclobutane enhances metabolic stability, lipophilicity, and binding interactions through conformational rigidity. The fluorine atom increases resistance to enzymatic oxidation [54], while the four-membered ring reduces metabolic cleavage susceptibility. It modulates lipophilicity by fine-tuning the hydrophilic-hydrophobic balance, optimizing LogP for membrane permeability and bioavailability—though aliphatic fluorination often reduces overall lipophilicity [55]. Fluorine’s high electronegativity enables dipole interactions, strengthening target binding [56], while the rigid cyclobutane ring locks the molecule into a bioactive conformation, minimizing entropic penalties [57]. These combined effects might enhance stability, solubility, and binding affinity, contributing to improved docking performance and potential biological activity.

Free energy calculations reveal a deeper free energy minimum (~2.61 kcal/mol) for the analogue compared to the parent compound (~2.85 kcal/mol), indicating stronger stabilization of the FLT3 protein-ligand complex. The analogue’s energy landscape is narrower, reducing entropic penalties and off-target interactions while improving binding affinity [58].

Cluster and dynamic stability analyses confirm that the analogue stabilizes the FLT3 structure more effectively than CHEMBL4444839, potentially improving therapeutic efficacy. The analogue’s interactions restrict protein motion, stabilizing it in a biologically favorable conformation [59]. DCCM analysis further demonstrates that the analogue better engages critical residues, promoting a more cohesive and functional dynamic state in FLT3.

Overall, the analogue outperforms CHEMBL4444839 in stability, binding strength, and adaptability, making it a more promising candidate for further development as a potent FLT3 inhibitor. Further experimental validation and molecular studies are recommended to confirm these findings and optimize the analogue’s therapeutic potential.

## 5. CONCLUSION

Our study presents a structurally optimized FLT3 inhibitor that exhibits superior binding affinity, stability, and pharmacokinetic properties compared to CHEMBL4444839, positioning it as a promising candidate for AML therapy. The incorporation of a fluorocyclobutane moiety significantly enhances ligand-protein interactions through increased hydrogen bonding, electrostatic stabilization, and hydrophobic engagement with key FLT3 residues. Molecular dynamics simulations and free energy calculations confirm the analogue’s reduced conformational fluctuations, lower entropic penalties, and stronger protein-ligand stabilization, while DCCM analysis reveals improved allosteric communication within FLT3. Furthermore, favorable ADME and toxicity assessments indicate enhanced metabolic stability, membrane permeability, and drug-likeness. These findings provide compelling computational evidence for the analogue’s therapeutic potential, warranting further *in vitro* and *in vivo* validation to assess its efficacy and translational applicability in AML treatment.

## 6.1 Acknowledgement

The authors thank NCBS, Bangalore for providing computational support. A warm heartfelt thanks to the staff and administration at the NCBS, Bangalore for their support.. We would also like to thank Dr. Abhijit Kayal, Senior Scientist II at Schrödinger, for his assistance with Free Energy Landscape (FEL) calculations and for his support with the Python scripts used in the analysis.

## 6. DECLARATIONS

### 6.3. Ethics Statement

This research did not involve any human participants or animal subjects, and therefore, no ethical approval was required.

### 6.6. Author Contributions

***Uddalak Das***: Conceptualization, Methodology, Formal Analysis, Writing—Original Draft, Writing—Review & Editing. ***Dheemanth Regati***: Conceptualization, Methodology, Formal Analysis, Writing—Original Draft, Writing—Review & Editing. ***R. Sowdhamini***: Supervision, Project Administration. ***Jitendra Kumar***: Supervision. All the authors have read and agreed to the published version of the manuscript.

### 6.7. Declaration of generative AI and AI-assisted technologies in the writing process

The writing of this research paper involved the use of generative AI and AI-assisted technologies only to enhance the clarity, coherence, and overall quality of the manuscript. The authors acknowledges the contributions of AI in the writing process while ensuring that the final content reflects the author’s own insights and interpretations of the literature. All interpretations and conclusions drawn in this manuscript are the sole responsibility of the author.

