## Supplementary material for "Rational design and comparative docking and simulation of modified FLT3 inhibitors: A study on enhanced binding stability and inhibition potency": Supplimentary Information

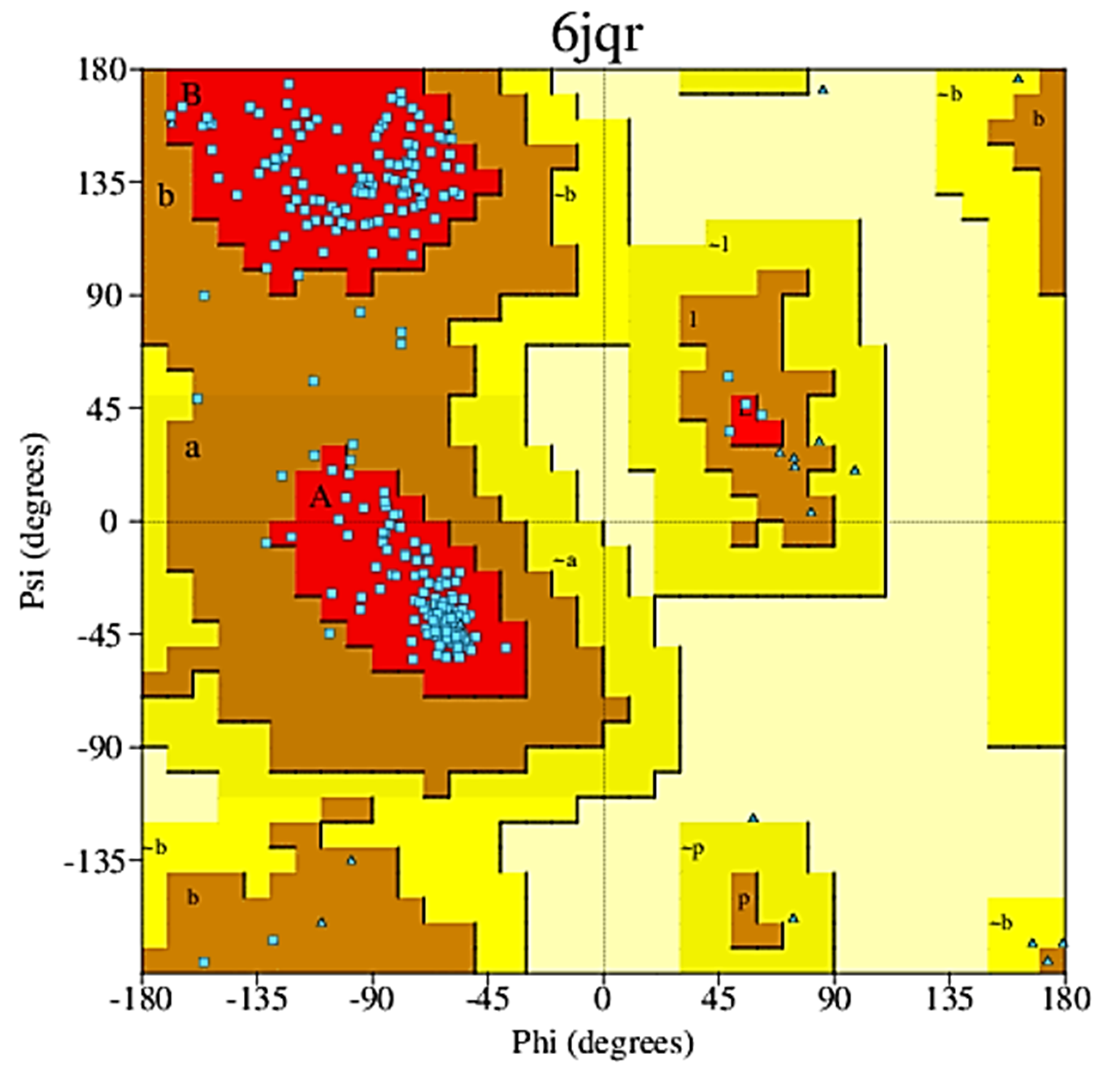

**Figure S1** Ramachandran plot. The color red indicates low-energy regions, yellow allowed regions, pale yellow generously allowed regions, and white disallowed regions. (Generated by PDBsum)

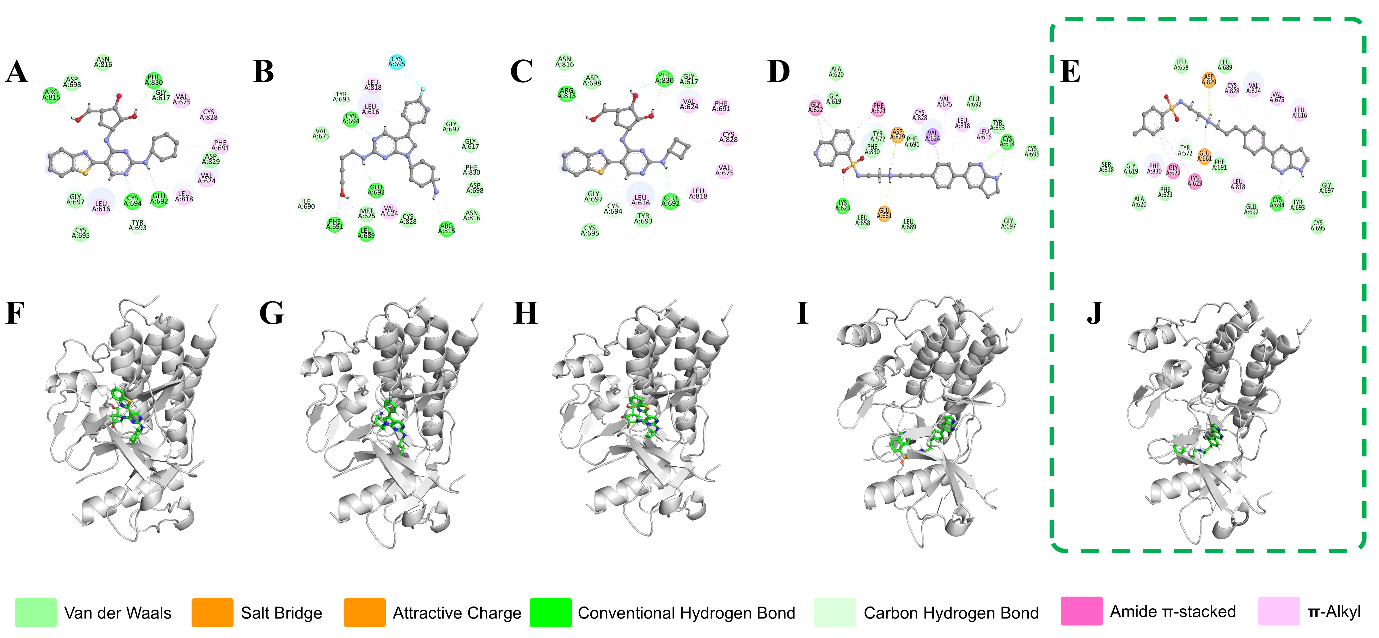

**Figure S2** The top 5 compounds based on docking score that were enumerated. The panels A-E shows the 2D interactions and the panels F-J shows the 3D interactions with FLT3 protein active site, with docking scores of -13.675, -13.115, -13.111, -12.500, and - -12.235 kcal/mol respectively. The compound in green dotted square shows the compounds that showed an analog with a enhances binding affinity.

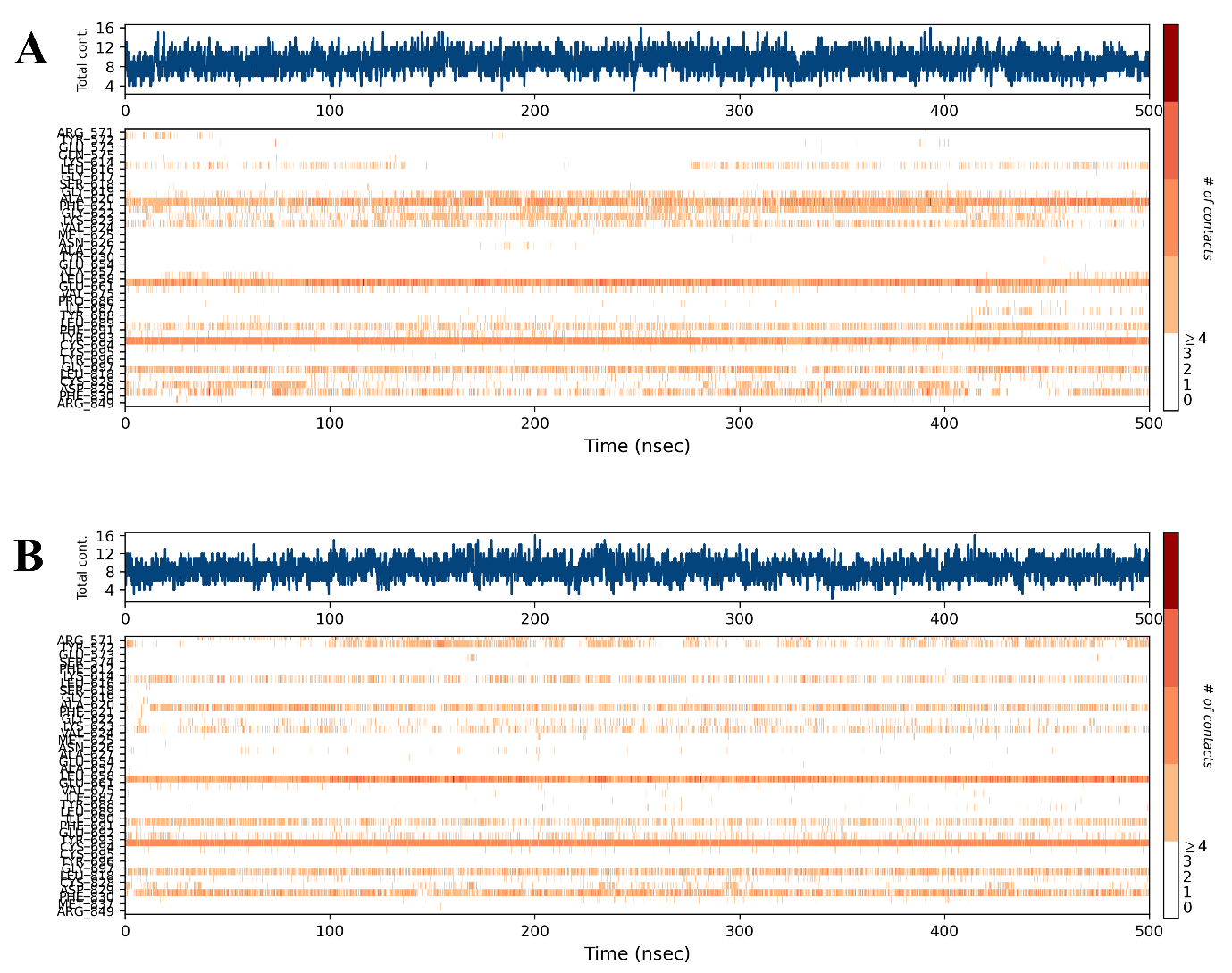

**Figure S3** Timeline representations fraction of interactions between protein-ligand complex (A) FLT3-CHEMBL4444839 (B) FLT3- CHEMBL4444839-analog. The top panel (in blue) shows the total number of specific contacts the protein makes with the ligand over the course of the trajectory. The bottom panel (in orange) shows which residues interact with the ligand in each trajectory frame.

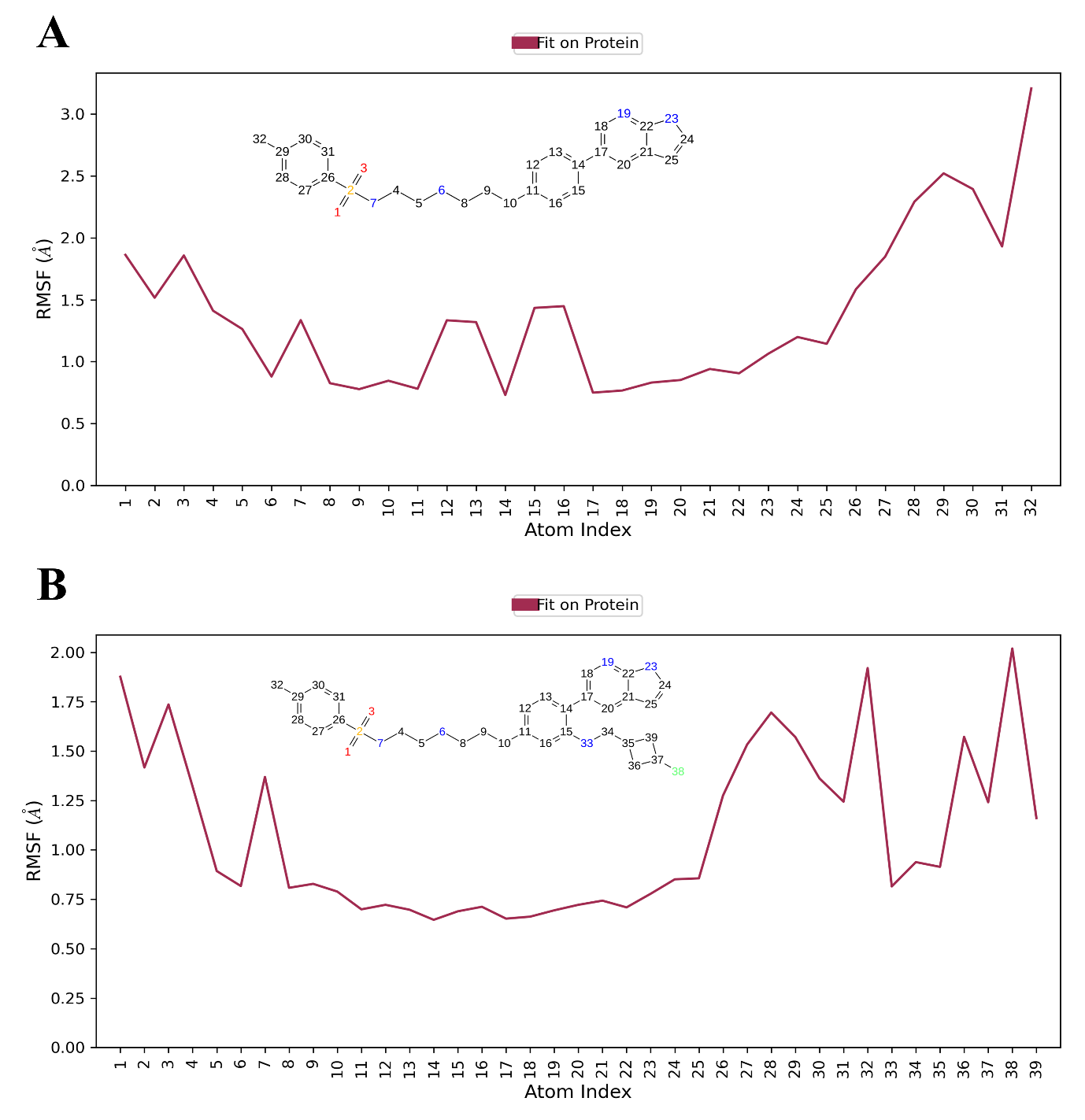

**Figure S4** The Ligand RMSF indicating changes in the ligand atom positions, giving insights into how the ligand fragments interact with the protein and their entropic role in the binding event. A: CHEMBL4444839; B: CHEMBL4444839-analog

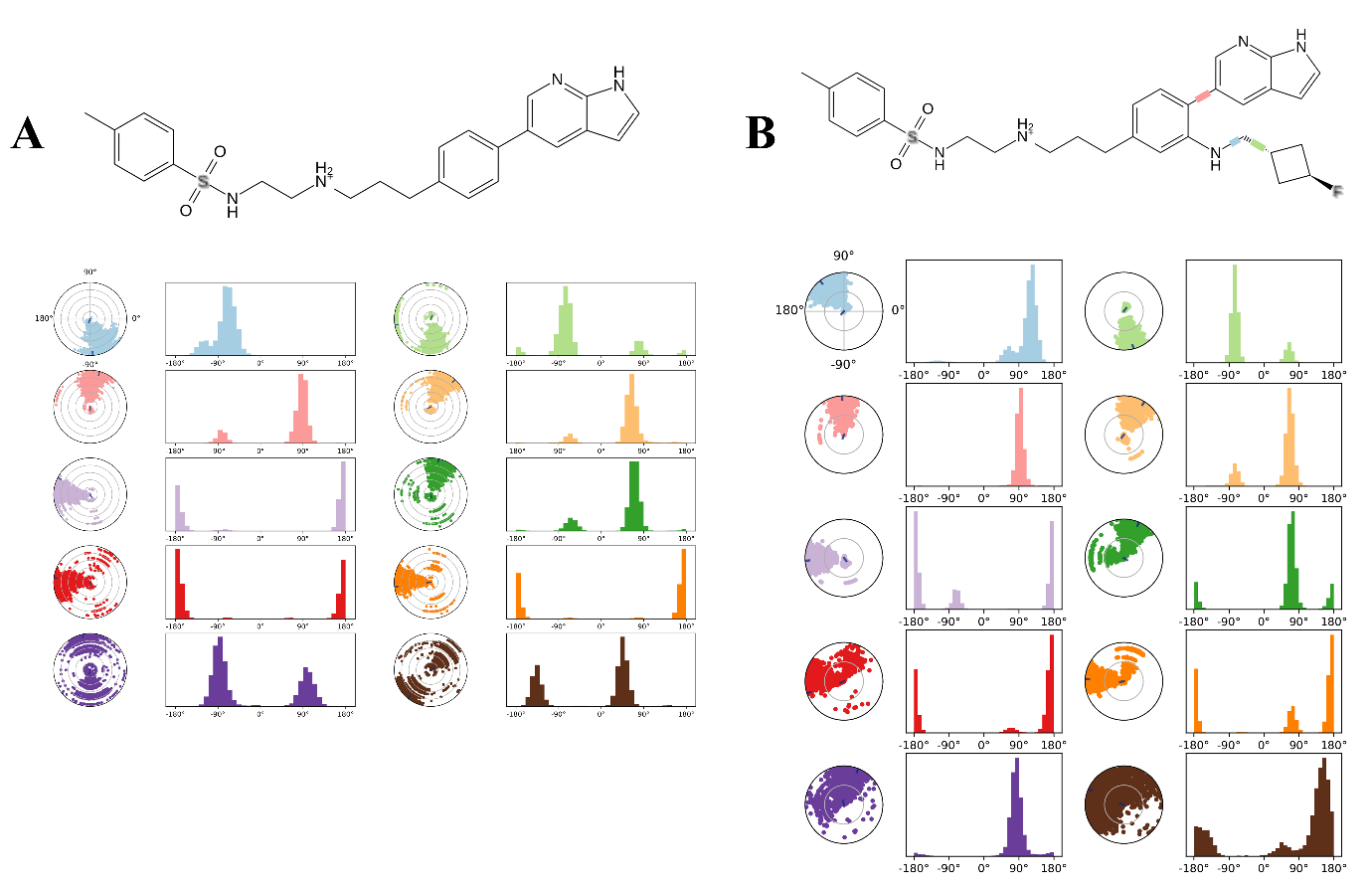

**Figure S5** The ligand torsions plot summarizing the conformational evolution of every rotatable bond (RB) in the ligands. A 2D schematic of CHEMBL4444839with color-coded rotatable bonds; A(ii) Dial plot and bar plots of the rotatable bonds of ligand CHEMBL4444839; B(i) 2D schematic of CHEMBL4444839-Analogue with color-coded rotatable bonds; A(ii) Dial plot and bar plots of the rotatable bonds of ligand CHEMBL4444839-Analogue

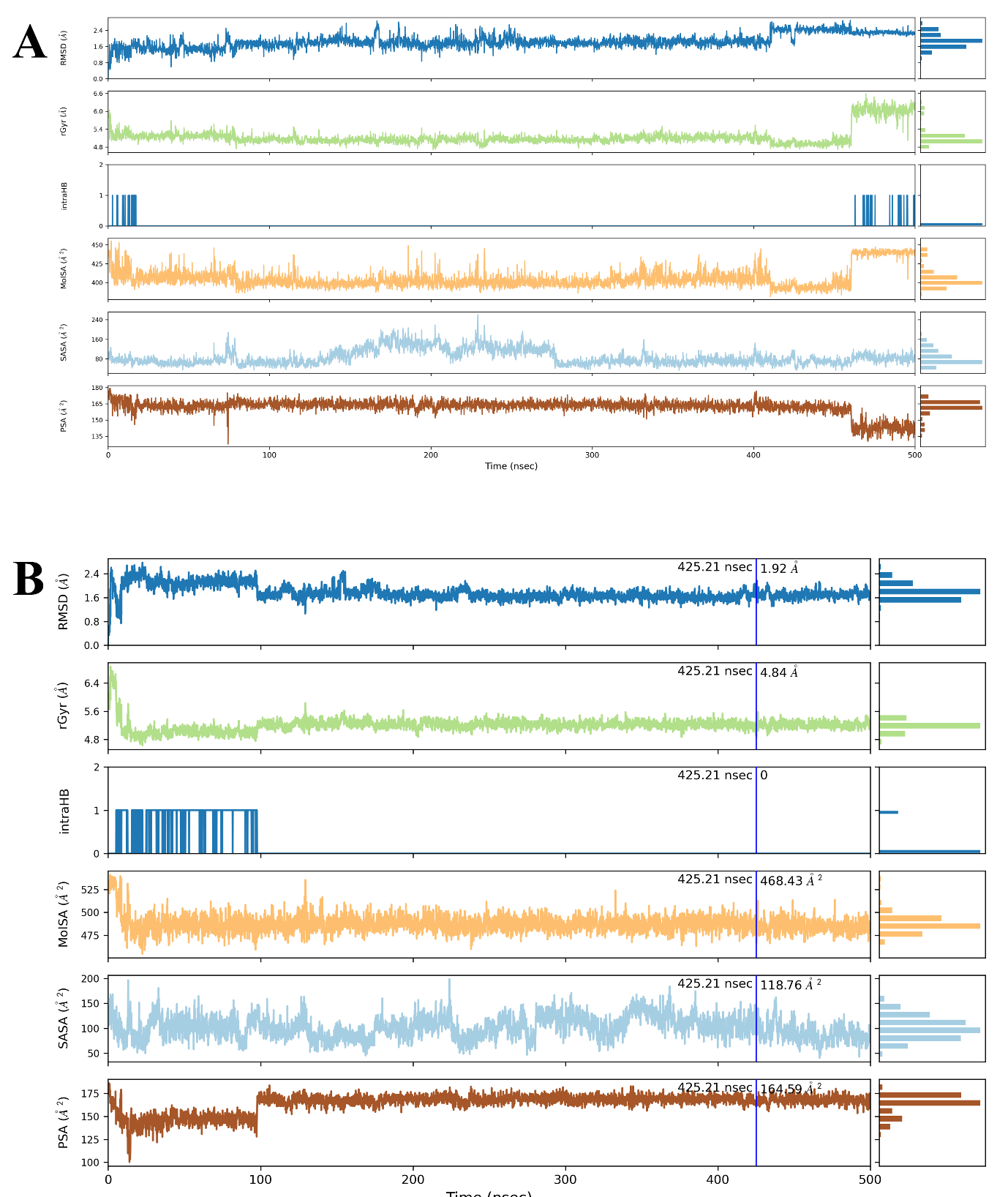

**Figure S6** Variations in the properties RMSD, rGyr, MolSA, intraHB, SASA, and PSA for the two ligands (A: CHEMBL4444839 and B: CHEMBL4444839-analog) during 500ns of MD simulation time.

**Table S1** Physiochemical properties of CHEMBL4444839 and CHEMBL4444839-analog

| Properties | CHEMBL4444839 | CHEMBL4444839-Analog |
| --- | --- | --- |
| Molecular Weight | 449.6 | 550.724 |
| Dipole Moment | 8.994 | 9.01 |
| Density | 0.973 | 0.989 |
| H-bond donors | 3 | 4 |
| H-bond acceptors | 6 | 7 |
| Number of rotatable bonds | 10 | 13 |
| Number of rings | 4 | 5 |
| Number of heteroatoms | 7 | 9 |
| Formal Charge | 0 | 0 |
| Number of rigid bonds | 24 | 28 |
| Sterio Centres | 0 | 0 |
| SASA | 795.463 | 900.247 |
| FOSA | 213.827 | 351.654 |
| FISA | 177.857 | 187.082 |
| PISA | 403.778 | 326.897 |
| WPSA | 0 | 34.614 |
| Total solvent accessible volume (in Å^3^) | 99.84 | 111.87 |
| Polarizability (in Å^3^) | 48.612 | 55.87 |
| Hexadecane/Gas partition coefficient | 16.487 | 18.407 |
| Octanol/Gas partition coefficient | 25.267 | 29.698 |
| Water/Gas partition coefficient | 14.673 | 16.265 |
| Octanol/Water partition coefficient | 3.723 | 4.644 |
| Number of non-conjugated amine groups | 1 | 1 |
| Number of amidine and guanidine groups | 0 | 0 |
| Number of carboxylic acid groups | 0 | 0 |
| Number of non-conjugated amide groups | 0 | 0 |
| Number of reactive functional groups | 0 | 0 |
| PM3 calculated ionization potential (negative of HOMO energy) (in eV) | 8.569 | 8.341 |
| PM3 calculated electron affinity (negative of LUMO energy) (in eV) | 0.742 | 0.736 |

**Table S2** Medicinal Chemistry properties of CHEMBL4444839 and CHEMBL4444839-analog

| Properties | CHEMBL4444839 | CHEMBL4444839-Analog |
| --- | --- | --- |
| Synthetic Accessibility | Easy | Easy |
| MCE-18 Value | 22 | 61.463 |
| Natural Product Likeliness Score | Higher | Lower |
| PAINS | 0 | 0 |
| ALARM NMR Rule | 1 violation | 2 violation |
| BMS Rule | 0 | 0 |
| Chelator Rule | 0 | 0 |
| Lipinski’s Rule of 5 violations | Accepted | Accepted |

**Table S3** ADMET properties of CHEMBL4444839 and CHEMBL4444839-analog

| **Properties** | | **CHEMBL4444839** | **CHEMBL4444839-Analog** |
| --- | --- | --- | --- |
| **A** | Water solubility (log mol/L) | 3.657 | -3.209 |
|  | Caco-2 permeability (in nm/sec) | 0.539 | 0.55 |
|  | Human intestinal absorption (in %) | 77.745 | 78.041 |
|  | Skin Permeability (log K_p_) | -2.735 | -2.735 |
|  | Human oral absorption (in %) | 79.284 | 70.15 |
|  | P-glycoprotein substrate | Yes | Yes |
|  | P-glycoprotein I inhibitor | Yes | Yes |
|  | P-glycoprotein II inhibitor | Yes | Yes |
|  | MDCK Permeability (in nm/sec) | 0 | 0 |
| **D** | Volume of distribution at steady state (in log L/kg) | 1.053 | 1.324 |
|  | Plasma protein binding (in %) | 96.5 | 93 |
|  | Fraction of unbound plasms | 0 | 0.03 |
|  | BBB Permeability (in logBB) | -0.796 | -1.273 |
|  | CNS permeability (in logPS) | -2.725 | -2.94 |
| **M** | CYP2D6 substrate | No | Yes |
|  | CYP3A4 substrate | Yes | Yes |
|  | CYPIA2 inhibitor | No | No |
|  | CYP2C19 inhibitor | Yes | Yes |
|  | CYP2C9 inhibitor | Yes | Yes |
|  | CYP2D6 inhibitor | Yes | Yes |
|  | CYP3A4 inhibitor | Yes | Yes |
| **E** | Total Clearance (in log ml/min/kg) | 1.223 | 1.277 |
|  | Renal OCT2 substrate | No | No |
| **T** | AMES toxicity | Yes | No |
|  | Max. tolerated dose (human) | 0.4 | 0.496 |
|  | hERG I inhibitor | No | No |
|  | hERG II inhibitor | Yes | Yes |
|  | Oral Rat Acute Toxicity (LD50) | 2.877 | 2.94 |
|  | Oral Rat Chronic Toxicity (LOAEL) | 2.08 | 2.08 |
|  | Hepatotoxicity | Yes | Yes |
|  | Skin Sensitisation | No | No |
|  | *T Pyriformis* toxicity (µg/L) | 0.286 | 0.285 |
|  | Minnow toxicity (log mM) | 0.656 | -0.102 |
|  | DILI Risk | High | High |
|  | Carcinogenicity | No | No |
|  | Eye corrosion | No | No |
|  | Eye irritation | No | No |
|  | Respiratory toxicity | Yes | Yes |

**Table S4** 2D and 3D structure of CHEMBL4444839 and CHEMBL4444839-analog

| Compounds | 2D Structure | 3D Structure |
| --- | --- | --- |
| CHEMBL4444839 | 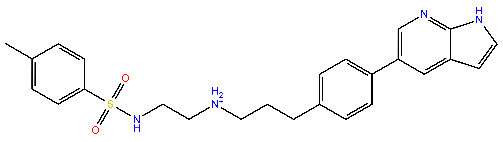 | 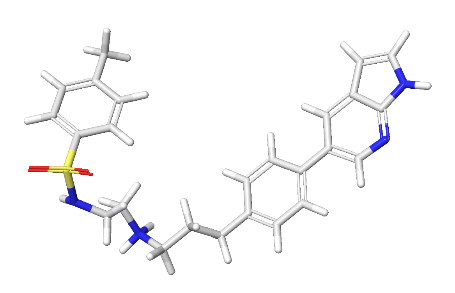 |
| CHEMBL4444839 Analog | 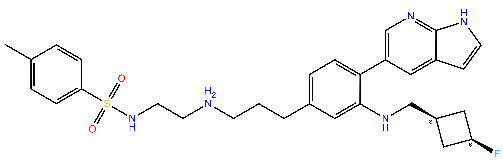 | 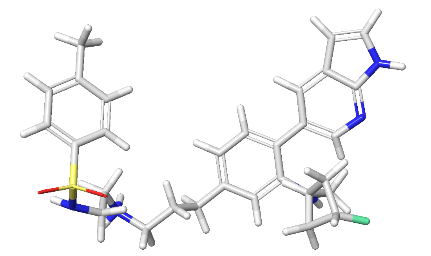 |

**Table S5** Post simulation MM-GBSA based Binding free energy (ΔG Bind in kcal/mol) for the 6JQR-CHEMBL4444839 Complex

| Time (in ns) | ΔG_bind_ Total | ΔG_bind_ Coulomb | ΔG_bind_ Covalent | ΔG_bind_ Hbond | ΔG_bind_ Lipo | ΔG_bind_ Packing | ΔG_bind_ Solv_GB | ΔG_bind_ vdW |
| --- | --- | --- | --- | --- | --- | --- | --- | --- |
| 1 | -81.6 | -46.8 | 1.3 | -2.6 | -24.7 | -1.8 | 51.5 | -58.6 |
| 2 | -92.6 | -52.8 | 2.8 | -3.4 | -31.5 | -1.8 | 59.4 | -65.3 |
| 3 | -67.3 | -42.9 | 1.7 | -2.9 | -21.2 | -0.3 | 52.9 | -54.5 |
| 4 | -81.1 | -59.5 | 0.1 | -3.0 | -23.1 | -0.8 | 63.2 | -58.0 |
| 5 | -82.0 | -53.2 | 2.1 | -3.2 | -22.8 | -0.5 | 58.3 | -62.6 |
| 6 | -75.1 | -60.5 | 1.2 | -2.6 | -23.6 | -0.1 | 74.4 | -63.9 |
| 7 | -84.8 | -56.0 | 2.7 | -3.7 | -25.2 | -1.1 | 58.8 | -60.3 |
| 8 | -82.0 | -55.8 | 2.8 | -3.2 | -25.1 | -1.4 | 60.0 | -59.5 |
| 9 | -79.5 | -48.1 | 2.6 | -2.0 | -24.3 | -0.7 | 54.1 | -61.1 |
| 10 | -77.7 | -48.6 | 2.7 | -2.9 | -24.8 | -0.4 | 59.0 | -62.7 |
| 11 | -83.5 | -55.5 | 3.8 | -2.6 | -25.2 | -2.2 | 62.4 | -64.2 |
| 12 | -74.0 | -32.9 | 2.7 | -2.7 | -23.5 | -1.3 | 44.1 | -60.5 |
| 13 | -77.8 | -31.3 | 2.9 | -3.3 | -23.0 | -0.6 | 38.6 | -61.1 |
| 14 | -79.2 | -37.2 | 3.4 | -2.7 | -23.9 | -0.7 | 45.1 | -63.1 |
| 15 | -78.8 | -30.9 | 3.2 | -2.9 | -24.5 | -1.4 | 39.0 | -61.2 |
| 16 | -84.8 | -53.0 | 1.3 | -3.5 | -25.4 | -1.2 | 55.6 | -58.5 |
| 17 | -78.4 | -41.4 | 2.6 | -3.3 | -22.9 | -0.5 | 47.5 | -60.5 |
| 18 | -79.5 | -49.7 | 1.8 | -3.7 | -21.4 | -0.7 | 53.5 | -59.4 |
| 19 | -84.9 | -65.2 | 1.2 | -3.4 | -23.6 | -0.9 | 65.3 | -58.2 |
| 20 | -77.8 | -46.7 | 3.2 | -3.1 | -25.2 | -1.1 | 56.1 | -61.1 |
| 21 | -79.8 | -45.1 | 2.4 | -2.8 | -23.7 | -0.6 | 53.2 | -63.1 |
| 22 | -78.2 | -46.9 | -0.2 | -2.6 | -22.4 | -0.7 | 53.6 | -59.0 |
| 23 | -76.8 | -48.6 | 0.4 | -2.5 | -23.5 | -0.4 | 55.2 | -57.4 |
| 24 | -78.0 | -32.7 | 0.7 | -2.2 | -23.9 | -0.9 | 43.2 | -62.3 |
| 25 | -74.8 | -82.3 | 1.6 | -2.5 | -22.1 | -0.6 | 88.6 | -57.5 |
| 26 | -76.0 | -64.2 | 1.9 | -3.0 | -21.2 | -0.1 | 68.9 | -58.3 |
| 27 | -75.4 | -43.4 | 1.9 | -2.2 | -23.7 | -0.8 | 52.3 | -59.6 |
| 28 | -80.7 | -40.4 | -0.3 | -3.0 | -23.4 | -0.3 | 47.2 | -60.5 |
| 29 | -75.4 | -46.2 | 0.7 | -2.5 | -23.1 | -0.5 | 56.1 | -59.8 |
| 30 | -71.6 | -45.9 | 2.2 | -2.4 | -21.5 | -1.1 | 54.1 | -56.8 |
| 31 | -79.5 | -37.0 | 1.1 | -2.9 | -24.6 | -0.6 | 45.4 | -61.0 |
| 32 | -83.8 | -46.1 | 1.1 | -2.8 | -26.1 | -0.5 | 51.2 | -60.5 |
| 33 | -85.7 | -44.7 | 0.1 | -3.4 | -24.7 | -1.3 | 51.0 | -62.7 |
| 34 | -82.8 | -50.3 | 2.6 | -2.8 | -25.4 | -0.7 | 55.1 | -61.2 |
| 35 | -78.7 | -50.8 | 2.0 | -2.9 | -24.0 | -0.9 | 60.2 | -62.4 |
| 36 | -90.7 | -60.5 | 1.2 | -2.5 | -26.9 | -0.8 | 63.2 | -64.5 |
| 37 | -87.0 | -61.6 | 1.0 | -2.3 | -27.7 | -1.3 | 70.4 | -65.4 |
| 38 | -83.2 | -45.0 | 2.8 | -2.5 | -26.6 | -1.5 | 57.4 | -67.8 |
| 39 | -79.5 | -51.9 | 1.6 | -3.0 | -24.3 | -0.5 | 57.0 | -58.4 |
| 40 | -83.7 | -61.5 | 3.0 | -2.9 | -25.4 | -2.1 | 68.2 | -63.0 |
| 41 | -82.5 | -52.9 | 1.4 | -2.5 | -26.0 | -0.7 | 62.4 | -64.3 |
| 42 | -88.5 | -67.3 | 2.1 | -3.1 | -26.5 | -2.0 | 68.8 | -60.5 |
| 43 | -84.2 | -42.0 | 1.4 | -2.6 | -26.8 | -1.4 | 50.9 | -63.7 |
| 44 | -86.1 | -47.9 | 1.6 | -3.1 | -26.8 | -1.4 | 55.7 | -64.1 |
| 45 | -82.4 | -42.7 | 0.4 | -2.7 | -27.1 | -0.8 | 51.9 | -61.6 |
| 46 | -89.4 | -58.2 | 2.2 | -3.1 | -30.3 | -1.7 | 62.3 | -60.6 |
| 47 | -81.2 | -54.4 | 1.1 | -2.9 | -25.7 | -1.2 | 59.0 | -57.1 |
| 48 | -74.0 | -44.0 | 1.9 | -2.5 | -23.6 | -2.5 | 49.8 | -53.0 |
| 49 | -83.8 | -52.8 | 2.1 | -3.2 | -25.4 | -1.5 | 58.8 | -61.8 |
| 50 | -83.9 | -48.3 | -0.6 | -2.5 | -25.9 | -1.1 | 57.6 | -63.2 |
| 51 | -83.5 | -46.1 | 2.0 | -3.1 | -26.2 | -1.0 | 55.3 | -64.4 |
| 52 | -85.3 | -48.7 | 1.2 | -2.8 | -24.2 | -0.5 | 51.4 | -61.8 |
| 53 | -91.3 | -48.3 | 0.6 | -2.5 | -29.2 | -1.0 | 54.6 | -65.5 |
| 54 | -81.4 | -45.0 | 2.5 | -2.7 | -23.1 | -0.7 | 48.9 | -61.3 |
| 55 | -82.0 | -52.3 | 1.6 | -2.7 | -23.8 | -0.9 | 56.6 | -60.5 |
| 56 | -77.1 | -55.6 | 1.5 | -3.0 | -21.6 | -0.8 | 62.6 | -60.2 |
| 57 | -78.8 | -45.1 | 2.5 | -2.9 | -23.7 | -0.8 | 55.6 | -64.5 |
| 58 | -80.8 | -48.6 | 1.2 | -2.7 | -24.0 | -0.8 | 56.3 | -62.1 |
| 59 | -80.0 | -38.3 | -0.3 | -2.8 | -23.1 | -0.8 | 45.0 | -59.7 |
| 60 | -84.5 | -43.8 | 1.9 | -3.0 | -26.3 | -1.4 | 51.2 | -63.0 |
| 61 | -73.7 | -47.9 | 1.5 | -2.2 | -23.3 | -0.7 | 59.7 | -60.8 |
| 62 | -76.6 | -30.1 | 1.4 | -2.8 | -22.8 | -1.6 | 37.8 | -58.4 |
| 63 | -72.8 | -29.2 | 1.2 | -2.5 | -22.3 | -0.2 | 37.9 | -57.7 |
| 64 | -81.3 | -48.1 | 0.1 | -2.6 | -24.7 | -0.6 | 56.8 | -62.2 |
| 65 | -84.2 | -53.1 | 0.9 | -2.4 | -25.1 | -1.1 | 61.1 | -64.5 |
| 66 | -76.1 | -47.1 | 1.4 | -2.5 | -23.0 | -0.8 | 56.1 | -60.1 |
| 67 | -73.8 | -47.4 | -0.1 | -2.1 | -23.9 | -0.8 | 56.3 | -55.8 |
| 68 | -69.3 | -19.5 | 0.9 | -2.4 | -21.4 | -0.4 | 31.1 | -57.6 |
| 69 | -79.8 | -67.8 | 0.7 | -2.5 | -22.9 | -0.7 | 71.9 | -58.6 |
| 70 | -80.7 | -49.6 | 1.2 | -2.3 | -22.9 | -0.7 | 55.3 | -61.8 |
| 71 | -78.1 | -50.3 | 0.9 | -2.6 | -23.2 | -0.7 | 57.6 | -59.9 |
| 72 | -82.7 | -60.0 | 1.3 | -2.8 | -24.5 | -0.6 | 65.3 | -61.4 |
| 73 | -83.8 | -63.2 | 1.1 | -2.6 | -25.4 | -0.9 | 68.0 | -60.8 |
| 74 | -62.0 | -50.7 | 1.4 | -2.3 | -21.4 | -1.0 | 58.9 | -46.9 |
| 75 | -77.1 | -38.3 | 1.7 | -2.6 | -25.0 | -0.8 | 43.0 | -55.0 |
| 76 | -79.7 | -53.5 | 1.3 | -3.2 | -25.5 | -2.4 | 57.6 | -54.1 |
| 77 | -78.4 | -45.7 | 3.3 | -2.9 | -24.1 | -1.3 | 53.5 | -61.1 |
| 78 | -79.4 | -36.3 | -0.8 | -3.2 | -22.4 | -2.2 | 42.8 | -57.3 |
| 79 | -74.5 | -46.4 | 2.0 | -2.6 | -23.3 | -3.5 | 52.1 | -52.8 |
| 80 | -80.0 | -63.3 | 1.0 | -2.3 | -24.8 | -1.4 | 69.2 | -58.3 |
| 81 | -81.3 | -55.6 | 1.6 | -3.3 | -24.0 | -1.4 | 58.5 | -57.1 |
| 82 | -85.9 | -53.3 | 3.0 | -3.6 | -25.5 | -0.8 | 59.5 | -65.1 |
| 83 | -87.4 | -40.2 | 1.1 | -3.4 | -27.5 | -1.9 | 48.0 | -63.4 |
| 84 | -85.5 | -58.6 | 2.4 | -2.9 | -29.8 | -1.4 | 64.2 | -59.4 |
| 85 | -75.8 | -50.5 | 1.6 | -3.2 | -23.7 | -0.7 | 59.1 | -58.3 |
| 86 | -82.6 | -53.1 | 0.7 | -3.2 | -23.1 | -0.8 | 54.8 | -58.0 |
| 87 | -78.3 | -39.9 | 1.4 | -3.1 | -23.5 | -1.4 | 46.7 | -58.5 |
| 88 | -77.5 | -45.2 | 1.3 | -3.1 | -22.4 | -2.0 | 50.4 | -56.6 |
| 89 | -77.1 | -38.6 | 2.4 | -2.8 | -22.3 | -0.6 | 46.8 | -62.0 |
| 90 | -82.2 | -48.6 | 3.1 | -3.2 | -24.3 | -1.4 | 54.9 | -62.6 |
| 91 | -79.1 | -45.5 | 1.9 | -2.1 | -25.3 | -0.6 | 55.2 | -62.7 |
| 92 | -75.7 | -51.8 | 2.3 | -2.4 | -25.8 | -0.8 | 64.4 | -61.6 |
| 93 | -84.4 | -61.7 | 3.5 | -2.8 | -30.0 | -2.1 | 68.6 | -60.1 |
| 94 | -76.8 | -52.2 | 1.0 | -2.3 | -25.1 | -1.1 | 62.6 | -59.7 |
| 95 | -79.6 | -41.4 | 0.5 | -2.6 | -25.5 | -2.4 | 48.4 | -56.6 |
| 96 | -84.9 | -50.4 | 1.2 | -2.3 | -28.0 | -3.7 | 60.3 | -61.9 |
| 97 | -84.5 | -50.8 | 0.7 | -2.9 | -29.3 | -1.9 | 62.7 | -63.1 |
| 98 | -75.9 | -44.9 | 2.2 | -2.8 | -24.7 | -0.8 | 52.6 | -57.5 |
| 99 | -81.0 | -59.6 | 1.3 | -2.2 | -25.9 | -0.8 | 69.9 | -63.5 |
| 100 | -76.2 | -45.1 | 2.9 | -2.6 | -24.2 | -1.2 | 54.3 | -60.3 |
| 101 | -79.9 | -49.1 | 2.6 | -2.6 | -24.1 | -1.0 | 58.5 | -64.1 |
| 102 | -80.0 | -29.3 | 1.9 | -2.6 | -26.0 | -1.8 | 41.7 | -63.8 |
| 103 | -77.6 | -71.2 | 2.7 | -2.8 | -25.8 | -1.7 | 82.2 | -61.1 |
| 104 | -87.2 | -51.3 | 1.2 | -3.2 | -26.9 | -1.3 | 60.6 | -66.3 |
| 105 | -78.9 | -43.5 | 2.0 | -2.7 | -25.3 | -1.7 | 51.3 | -59.0 |
| 106 | -78.1 | -48.7 | 2.4 | -2.6 | -25.5 | -1.0 | 58.6 | -61.3 |
| 107 | -78.8 | -46.9 | 1.0 | -2.8 | -24.6 | -1.3 | 54.4 | -58.6 |
| 108 | -86.8 | -57.7 | 1.9 | -2.4 | -28.6 | -1.8 | 65.8 | -63.9 |
| 109 | -75.0 | -59.6 | 3.5 | -2.4 | -24.0 | -1.6 | 67.3 | -58.2 |
| 110 | -86.7 | -71.3 | 0.5 | -3.1 | -26.8 | -1.4 | 77.9 | -62.4 |
| 111 | -93.0 | -60.9 | 1.0 | -3.1 | -28.1 | -1.0 | 66.6 | -67.6 |
| 112 | -82.1 | -55.9 | 2.1 | -2.6 | -27.2 | -1.4 | 61.4 | -58.5 |
| 113 | -73.9 | -40.3 | 1.2 | -2.8 | -23.5 | -1.1 | 48.6 | -56.0 |
| 114 | -76.1 | -64.2 | 1.1 | -3.2 | -23.9 | -2.6 | 72.1 | -55.4 |
| 115 | -70.4 | -39.1 | 2.4 | -2.6 | -24.1 | -1.9 | 50.6 | -55.7 |
| 116 | -70.9 | -48.3 | 1.6 | -2.8 | -23.5 | -0.9 | 61.8 | -58.9 |
| 117 | -78.2 | -47.3 | 2.0 | -2.8 | -24.1 | -0.9 | 54.1 | -59.2 |
| 118 | -73.8 | -48.5 | 0.2 | -3.1 | -24.0 | -3.5 | 59.1 | -54.1 |
| 119 | -74.6 | -36.2 | 2.2 | -2.6 | -24.1 | -0.9 | 46.8 | -59.8 |
| 120 | -77.0 | -57.1 | 3.7 | -2.5 | -25.5 | -0.8 | 62.0 | -56.8 |
| 121 | -79.6 | -47.6 | 1.6 | -2.3 | -26.1 | -1.4 | 55.9 | -59.7 |
| 122 | -78.7 | -48.7 | 1.0 | -2.2 | -28.1 | -1.3 | 59.5 | -58.9 |
| 123 | -78.3 | -49.5 | 0.8 | -3.1 | -24.9 | -1.4 | 58.6 | -58.8 |
| 124 | -68.5 | -46.7 | 1.4 | -2.6 | -22.4 | -1.7 | 59.6 | -56.0 |
| 125 | -78.3 | -34.7 | 2.4 | -2.9 | -25.8 | -2.7 | 46.5 | -61.1 |
| 126 | -80.6 | -51.5 | 2.5 | -2.4 | -27.3 | -4.2 | 62.1 | -59.8 |
| 127 | -75.5 | -57.8 | 0.8 | -2.9 | -25.7 | -0.7 | 67.5 | -56.7 |
| 128 | -83.7 | -51.1 | 1.8 | -2.5 | -28.0 | -1.5 | 59.4 | -61.9 |
| 129 | -76.8 | -57.2 | 0.6 | -2.5 | -24.2 | -1.3 | 66.4 | -58.6 |
| 130 | -80.3 | -38.5 | 2.0 | -2.8 | -26.7 | -2.1 | 50.9 | -63.0 |
| 131 | -73.9 | -68.5 | 1.2 | -2.8 | -22.9 | -2.6 | 76.7 | -54.9 |
| 132 | -75.8 | -40.5 | 1.6 | -2.6 | -25.8 | -1.2 | 51.4 | -58.8 |
| 133 | -69.9 | -55.5 | 1.5 | -2.6 | -24.0 | -1.4 | 66.5 | -54.2 |
| 134 | -82.8 | -40.1 | 2.7 | -2.7 | -27.8 | -2.4 | 48.0 | -60.5 |
| 135 | -79.5 | -47.4 | 1.6 | -3.4 | -26.3 | -3.1 | 59.5 | -60.3 |
| 136 | -80.7 | -46.2 | 0.5 | -3.5 | -26.3 | -2.6 | 54.1 | -56.6 |
| 137 | -71.4 | -40.2 | 1.0 | -3.1 | -24.2 | -1.8 | 49.3 | -52.4 |
| 138 | -76.9 | -59.8 | 1.8 | -3.1 | -25.7 | -4.1 | 70.5 | -56.4 |
| 139 | -76.0 | -60.4 | 2.1 | -3.5 | -27.1 | -2.7 | 65.4 | -49.9 |
| 140 | -73.3 | -54.1 | 0.3 | -3.2 | -22.4 | -2.4 | 61.2 | -52.8 |
| 141 | -75.1 | -51.2 | 3.4 | -3.7 | -25.4 | -2.0 | 53.3 | -49.5 |
| 142 | -76.9 | -64.1 | 2.4 | -3.4 | -23.4 | -2.9 | 68.3 | -53.8 |
| 143 | -69.3 | -51.4 | 0.9 | -2.8 | -23.5 | -1.1 | 59.8 | -51.3 |
| 144 | -78.1 | -64.5 | 1.9 | -3.4 | -25.6 | -5.0 | 69.2 | -50.5 |
| 145 | -67.8 | -39.4 | -0.1 | -3.0 | -21.5 | -0.6 | 49.1 | -52.3 |
| 146 | -75.7 | -66.0 | 1.1 | -3.3 | -23.8 | -5.0 | 76.2 | -54.9 |
| 147 | -79.5 | -75.0 | 0.6 | -3.6 | -23.3 | -3.1 | 79.0 | -54.1 |
| 148 | -73.8 | -61.6 | 0.8 | -3.1 | -22.2 | -3.9 | 68.0 | -51.8 |
| 149 | -70.9 | -51.3 | 0.8 | -3.0 | -25.3 | -1.1 | 60.8 | -51.7 |
| 150 | -69.4 | -58.8 | 0.9 | -3.1 | -23.4 | -2.5 | 67.0 | -49.4 |
| 151 | -78.2 | -60.0 | 0.1 | -3.0 | -24.4 | -3.6 | 68.9 | -56.3 |
| 152 | -69.3 | -65.0 | -0.1 | -2.7 | -22.7 | -2.1 | 76.9 | -53.6 |
| 153 | -71.3 | -57.0 | 0.1 | -3.8 | -21.5 | -1.0 | 61.8 | -49.9 |
| 154 | -67.2 | -41.9 | 1.1 | -3.0 | -23.1 | -1.2 | 50.7 | -49.9 |
| 155 | -70.9 | -49.0 | 0.4 | -3.4 | -22.3 | -0.9 | 57.9 | -53.7 |
| 156 | -79.3 | -45.6 | 0.9 | -3.8 | -25.8 | -3.8 | 52.2 | -53.3 |
| 157 | -69.4 | -51.4 | 2.2 | -3.3 | -21.9 | -2.7 | 61.1 | -53.3 |
| 158 | -71.3 | -45.5 | 2.2 | -3.3 | -23.1 | -3.7 | 56.1 | -54.0 |
| 159 | -73.4 | -60.0 | -0.2 | -3.3 | -23.5 | -3.2 | 68.2 | -51.3 |
| 160 | -65.4 | -51.1 | 0.9 | -4.0 | -20.3 | -1.8 | 58.1 | -47.2 |
| 161 | -62.4 | -46.3 | 1.1 | -3.1 | -20.1 | -1.6 | 53.4 | -45.9 |
| 162 | -70.5 | -60.5 | 1.2 | -3.7 | -20.9 | -0.8 | 64.9 | -50.7 |
| 163 | -73.9 | -41.4 | 0.4 | -3.1 | -24.1 | -4.2 | 49.6 | -51.0 |
| 164 | -79.9 | -65.5 | 0.9 | -3.5 | -23.9 | -3.1 | 69.3 | -54.3 |
| 165 | -67.1 | -56.1 | 1.5 | -3.6 | -21.7 | -1.8 | 67.8 | -53.3 |
| 166 | -78.9 | -31.1 | 0.6 | -3.4 | -24.5 | -1.7 | 36.9 | -55.7 |
| 167 | -74.6 | -56.8 | 0.7 | -3.3 | -23.0 | -0.8 | 60.8 | -52.2 |
| 168 | -72.8 | -58.4 | 0.7 | -2.8 | -22.6 | -1.4 | 65.4 | -53.7 |
| 169 | -59.2 | -61.1 | 2.2 | -2.8 | -18.3 | -2.7 | 68.2 | -44.7 |
| 170 | -66.0 | -51.0 | 1.7 | -3.2 | -23.0 | -4.1 | 61.9 | -48.2 |
| 171 | -54.6 | -58.5 | 4.5 | -2.5 | -17.3 | -3.9 | 69.1 | -45.8 |
| 172 | -65.5 | -71.7 | 2.9 | -2.2 | -22.9 | -1.9 | 81.1 | -50.8 |
| 173 | -61.4 | -65.7 | 1.9 | -3.2 | -19.5 | -2.0 | 74.3 | -47.1 |
| 174 | -66.7 | -57.2 | 1.9 | -3.3 | -22.3 | -2.9 | 66.0 | -49.0 |
| 175 | -65.9 | -49.2 | 0.9 | -3.3 | -20.6 | -1.7 | 56.3 | -48.4 |
| 176 | -69.5 | -63.0 | 3.0 | -2.8 | -21.7 | -1.5 | 68.7 | -52.1 |
| 177 | -67.8 | -59.3 | 1.5 | -3.4 | -19.9 | -3.4 | 64.0 | -47.4 |
| 178 | -57.0 | -39.1 | 1.5 | -2.8 | -17.8 | -1.0 | 47.7 | -45.4 |
| 179 | -62.9 | -46.5 | 3.1 | -3.8 | -17.0 | -1.4 | 46.7 | -43.9 |
| 180 | -65.6 | -54.5 | 2.4 | -3.3 | -21.2 | -1.5 | 61.3 | -48.9 |
| 181 | -58.8 | -39.3 | 0.0 | -2.5 | -20.7 | -0.8 | 52.9 | -48.4 |
| 182 | -63.3 | -54.8 | 1.2 | -3.1 | -20.4 | -2.7 | 63.8 | -47.4 |
| 183 | -70.3 | -44.4 | 1.2 | -3.2 | -22.4 | -4.4 | 51.7 | -48.8 |
| 184 | -61.8 | -61.5 | 0.8 | -2.9 | -19.7 | -3.3 | 72.6 | -47.8 |
| 185 | -59.5 | -40.7 | 1.3 | -2.3 | -19.0 | -1.7 | 49.4 | -46.4 |
| 186 | -57.4 | -46.8 | 2.2 | -2.9 | -19.2 | -1.7 | 54.3 | -43.3 |
| 187 | -62.8 | -55.1 | 1.8 | -2.6 | -20.6 | -2.1 | 67.0 | -51.2 |
| 188 | -57.4 | -40.0 | 1.2 | -3.1 | -19.4 | -0.7 | 49.2 | -44.5 |
| 189 | -60.5 | -40.6 | 0.7 | -2.9 | -20.8 | -0.9 | 50.7 | -46.8 |
| 190 | -66.4 | -44.3 | 0.4 | -3.3 | -20.0 | -0.6 | 51.8 | -50.4 |
| 191 | -65.1 | -59.5 | 0.3 | -3.2 | -20.2 | -0.6 | 67.5 | -49.4 |
| 192 | -69.2 | -59.5 | 3.1 | -3.1 | -20.9 | -3.0 | 65.4 | -51.3 |
| 193 | -67.0 | -38.9 | 1.7 | -3.1 | -22.4 | -2.8 | 48.9 | -50.6 |
| 194 | -70.9 | -59.4 | 2.1 | -3.4 | -21.4 | -4.7 | 65.3 | -49.5 |
| 195 | -63.7 | -40.1 | 2.7 | -2.2 | -19.6 | -0.6 | 48.1 | -52.0 |
| 196 | -68.4 | -38.5 | 2.0 | -2.8 | -20.2 | -2.6 | 44.2 | -50.5 |
| 197 | -66.1 | -53.4 | 2.0 | -3.3 | -20.5 | -0.5 | 57.5 | -47.9 |
| 198 | -69.8 | -46.7 | 0.9 | -3.2 | -21.2 | -1.5 | 53.6 | -51.7 |
| 199 | -65.4 | -46.8 | 1.6 | -2.8 | -20.3 | -2.1 | 53.1 | -48.1 |
| 200 | -74.3 | -65.2 | 1.8 | -3.4 | -22.5 | -2.9 | 69.1 | -51.2 |
| 201 | -74.1 | -55.5 | 0.7 | -3.1 | -19.4 | -2.5 | 58.3 | -52.4 |
| 202 | -77.5 | -46.7 | 1.3 | -3.4 | -23.3 | -1.9 | 50.6 | -54.0 |
| 203 | -65.3 | -53.5 | 1.2 | -3.2 | -19.6 | -1.1 | 59.6 | -48.8 |
| 204 | -75.6 | -60.7 | 1.1 | -3.6 | -21.9 | -2.9 | 65.1 | -52.7 |
| 205 | -63.5 | -45.4 | 3.1 | -3.2 | -20.7 | -2.5 | 52.5 | -47.3 |
| 206 | -63.2 | -56.3 | 1.8 | -3.3 | -20.4 | -0.8 | 61.5 | -45.6 |
| 207 | -70.3 | -45.7 | 1.3 | -3.5 | -20.8 | -0.7 | 49.4 | -50.2 |
| 208 | -67.7 | -51.0 | 0.8 | -3.7 | -20.6 | -0.7 | 57.8 | -50.3 |
| 209 | -69.3 | -59.4 | 3.2 | -3.0 | -23.0 | -1.5 | 67.3 | -53.0 |
| 210 | -63.4 | -47.9 | 2.0 | -3.1 | -19.6 | -0.6 | 56.0 | -50.2 |
| 211 | -66.4 | -38.4 | 2.8 | -2.8 | -21.2 | -4.2 | 49.4 | -51.8 |
| 212 | -71.6 | -60.2 | 1.8 | -3.3 | -21.5 | -1.5 | 64.8 | -51.6 |
| 213 | -74.5 | -51.7 | 2.4 | -3.1 | -23.5 | -4.4 | 58.4 | -52.5 |
| 214 | -75.0 | -71.2 | 2.0 | -3.5 | -22.5 | -3.5 | 75.3 | -51.5 |
| 215 | -74.4 | -63.9 | 2.0 | -3.1 | -23.4 | -2.7 | 70.9 | -54.2 |
| 216 | -69.2 | -52.2 | 2.6 | -3.3 | -21.4 | -3.8 | 58.7 | -49.9 |
| 217 | -77.7 | -56.9 | 2.9 | -3.3 | -24.1 | -2.2 | 61.3 | -55.4 |
| 218 | -72.5 | -53.2 | 2.6 | -3.2 | -22.8 | -0.9 | 57.0 | -52.1 |
| 219 | -67.9 | -58.7 | 3.2 | -3.1 | -21.8 | -2.1 | 66.4 | -51.8 |
| 220 | -62.6 | -56.1 | 4.6 | -3.2 | -19.3 | -2.7 | 63.6 | -49.5 |
| 221 | -60.6 | -55.8 | 1.7 | -2.2 | -21.2 | -1.6 | 62.3 | -43.7 |
| 222 | -72.8 | -61.1 | 2.6 | -3.5 | -20.8 | -2.6 | 63.0 | -50.3 |
| 223 | -67.1 | -54.6 | 2.5 | -3.6 | -20.2 | -1.3 | 57.1 | -47.1 |
| 224 | -66.2 | -49.1 | 3.3 | -3.1 | -21.0 | -1.0 | 54.5 | -49.8 |
| 225 | -69.1 | -53.0 | 1.2 | -3.0 | -21.3 | -1.4 | 60.2 | -51.9 |
| 226 | -64.1 | -54.0 | 2.8 | -3.4 | -20.0 | -1.7 | 61.1 | -48.8 |
| 227 | -68.2 | -62.5 | 0.9 | -3.3 | -20.5 | -1.0 | 70.0 | -51.8 |
| 228 | -64.3 | -54.2 | 4.2 | -2.5 | -21.7 | -1.4 | 63.4 | -52.1 |
| 229 | -60.0 | -42.8 | 1.3 | -2.7 | -21.0 | -0.9 | 53.6 | -47.4 |
| 230 | -60.0 | -46.0 | 2.4 | -2.8 | -17.3 | -1.0 | 53.2 | -48.3 |
| 231 | -68.3 | -48.1 | 2.8 | -3.0 | -22.7 | -2.1 | 54.7 | -49.9 |
| 232 | -66.1 | -52.9 | 3.1 | -3.5 | -21.6 | -2.5 | 59.4 | -48.0 |
| 233 | -68.6 | -60.5 | 3.2 | -2.9 | -21.0 | -2.0 | 65.2 | -50.6 |
| 234 | -62.1 | -40.1 | 3.2 | -2.6 | -19.0 | -3.8 | 48.2 | -47.9 |
| 235 | -63.2 | -44.6 | 3.4 | -3.2 | -22.1 | -1.5 | 54.1 | -49.4 |
| 236 | -69.5 | -64.6 | 1.9 | -3.3 | -21.8 | -4.3 | 72.3 | -49.8 |
| 237 | -73.6 | -52.7 | 2.3 | -3.7 | -23.3 | -3.2 | 58.1 | -51.2 |
| 238 | -64.7 | -53.1 | 1.4 | -3.4 | -20.7 | -0.7 | 59.8 | -48.0 |
| 239 | -63.3 | -64.6 | 2.5 | -3.0 | -20.9 | -2.4 | 73.5 | -48.3 |
| 240 | -71.8 | -62.8 | 0.7 | -3.0 | -23.4 | -3.6 | 74.5 | -54.2 |
| 241 | -70.6 | -52.0 | 2.8 | -3.5 | -21.9 | -1.1 | 56.9 | -51.7 |
| 242 | -55.4 | -33.4 | 1.7 | -3.1 | -17.5 | -0.1 | 42.8 | -45.7 |
| 243 | -66.1 | -52.3 | 3.7 | -2.8 | -20.2 | -1.0 | 57.4 | -51.0 |
| 244 | -65.6 | -51.6 | 2.9 | -2.9 | -21.0 | -0.7 | 57.0 | -49.4 |
| 245 | -68.4 | -53.2 | 0.8 | -3.0 | -20.6 | -0.7 | 59.6 | -51.1 |
| 246 | -71.0 | -50.2 | 2.0 | -3.2 | -23.0 | -3.9 | 62.2 | -54.8 |
| 247 | -71.7 | -43.4 | 0.5 | -3.4 | -23.2 | -3.4 | 49.7 | -48.5 |
| 248 | -68.8 | -51.1 | 1.6 | -3.0 | -23.3 | -3.3 | 64.8 | -54.5 |
| 249 | -67.5 | -57.2 | 0.4 | -2.2 | -21.0 | -3.3 | 63.7 | -47.9 |
| 250 | -70.6 | -48.4 | 0.8 | -2.4 | -23.1 | -3.1 | 59.8 | -54.2 |
| 251 | -68.1 | -48.8 | 1.3 | -3.3 | -19.5 | -1.3 | 54.1 | -50.5 |
| 252 | -70.6 | -57.1 | 1.2 | -3.4 | -22.2 | -0.9 | 64.8 | -53.0 |
| 253 | -74.9 | -52.8 | 0.4 | -2.9 | -23.7 | -3.9 | 59.4 | -51.4 |
| 254 | -67.8 | -52.0 | 1.6 | -2.6 | -22.7 | -1.5 | 58.7 | -49.3 |
| 255 | -77.1 | -65.4 | 2.2 | -3.3 | -22.9 | -3.9 | 68.8 | -52.5 |
| 256 | -71.2 | -54.4 | 1.3 | -3.0 | -23.3 | -2.9 | 63.5 | -52.4 |
| 257 | -67.3 | -58.6 | 0.9 | -3.6 | -20.9 | -2.5 | 64.3 | -47.0 |
| 258 | -72.4 | -68.6 | 0.7 | -3.5 | -18.9 | -3.2 | 70.2 | -49.1 |
| 259 | -72.2 | -63.2 | 0.5 | -3.5 | -20.3 | -2.2 | 65.7 | -49.2 |
| 260 | -68.1 | -48.5 | 1.7 | -3.8 | -20.4 | -0.9 | 53.8 | -50.1 |
| 261 | -72.0 | -58.5 | 1.6 | -3.3 | -22.4 | -2.2 | 62.4 | -49.5 |
| 262 | -70.5 | -42.5 | 0.4 | -3.5 | -22.2 | -1.8 | 48.0 | -48.9 |
| 263 | -67.9 | -57.2 | 2.1 | -3.3 | -21.7 | -1.6 | 62.4 | -48.7 |
| 264 | -68.5 | -54.7 | 1.1 | -3.4 | -21.9 | -1.9 | 61.8 | -49.5 |
| 265 | -73.7 | -51.6 | 1.6 | -3.6 | -23.3 | -3.5 | 60.1 | -53.4 |
| 266 | -80.6 | -53.4 | 0.2 | -3.1 | -24.9 | -4.9 | 59.1 | -53.7 |
| 267 | -79.1 | -41.9 | 1.1 | -3.6 | -25.1 | -4.5 | 49.2 | -54.3 |
| 268 | -77.2 | -43.6 | 1.0 | -3.5 | -23.5 | -4.4 | 50.7 | -53.9 |
| 269 | -77.5 | -59.6 | 1.4 | -3.4 | -23.7 | -4.2 | 66.2 | -54.3 |
| 270 | -69.5 | -54.3 | 1.5 | -3.2 | -21.6 | -2.1 | 61.5 | -51.2 |
| 271 | -73.7 | -48.7 | 1.6 | -3.1 | -23.1 | -1.0 | 54.8 | -54.2 |
| 272 | -74.3 | -52.5 | 0.6 | -3.3 | -24.3 | -1.5 | 59.3 | -52.7 |
| 273 | -73.6 | -60.1 | 0.6 | -2.5 | -23.4 | -2.5 | 64.2 | -49.8 |
| 274 | -70.6 | -34.1 | 0.4 | -2.4 | -23.0 | -3.6 | 41.1 | -49.2 |
| 275 | -72.5 | -53.1 | 0.6 | -2.8 | -23.9 | -2.9 | 64.0 | -54.4 |
| 276 | -70.5 | -58.0 | 2.1 | -3.4 | -22.1 | -5.0 | 63.8 | -48.0 |
| 277 | -68.2 | -53.5 | 2.6 | -2.9 | -22.5 | -3.4 | 62.2 | -50.7 |
| 278 | -78.9 | -48.0 | 2.0 | -2.9 | -25.9 | -1.0 | 59.5 | -62.7 |
| 279 | -71.3 | -63.0 | 2.2 | -3.0 | -23.3 | -1.3 | 71.3 | -54.2 |
| 280 | -76.2 | -50.7 | 1.1 | -3.1 | -24.1 | -1.1 | 59.4 | -57.8 |
| 281 | -79.1 | -56.9 | 3.3 | -3.3 | -26.9 | -1.3 | 65.0 | -58.9 |
| 282 | -72.3 | -44.0 | 4.0 | -3.2 | -24.1 | -1.3 | 54.8 | -58.5 |
| 283 | -78.0 | -42.5 | 1.7 | -2.5 | -24.8 | -0.9 | 51.7 | -60.7 |
| 284 | -77.5 | -46.3 | 2.5 | -2.6 | -25.2 | -1.1 | 57.3 | -62.1 |
| 285 | -83.3 | -53.1 | 1.2 | -2.9 | -27.2 | -0.9 | 65.9 | -66.2 |
| 286 | -71.4 | -65.7 | 1.8 | -2.3 | -23.5 | -1.3 | 76.3 | -56.8 |
| 287 | -79.7 | -61.8 | 0.0 | -2.7 | -26.1 | -1.6 | 71.7 | -59.0 |
| 288 | -84.2 | -60.5 | 1.7 | -3.1 | -26.7 | -1.5 | 68.6 | -62.7 |
| 289 | -76.4 | -59.7 | 0.7 | -3.3 | -23.8 | -1.2 | 67.2 | -56.3 |
| 290 | -65.0 | -65.2 | 0.6 | -2.8 | -22.2 | -0.9 | 79.5 | -53.9 |
| 291 | -77.2 | -50.8 | 0.8 | -2.8 | -24.8 | -1.0 | 63.8 | -62.3 |
| 292 | -71.6 | -38.8 | 1.1 | -2.7 | -23.3 | -1.3 | 50.3 | -56.9 |
| 293 | -85.3 | -59.8 | 0.6 | -2.9 | -29.3 | -1.2 | 71.1 | -63.9 |
| 294 | -73.9 | -61.8 | 1.2 | -2.5 | -25.2 | -1.0 | 74.0 | -58.6 |
| 295 | -74.1 | -37.2 | 2.1 | -3.2 | -23.7 | -1.1 | 48.1 | -59.1 |
| 296 | -75.4 | -49.4 | 1.5 | -2.4 | -25.0 | -0.9 | 61.6 | -60.8 |
| 297 | -76.7 | -47.6 | 0.1 | -2.9 | -24.3 | -1.1 | 56.6 | -57.6 |
| 298 | -80.1 | -62.4 | 0.2 | -2.2 | -27.1 | -1.3 | 76.5 | -63.9 |
| 299 | -80.8 | -48.1 | 1.1 | -2.6 | -25.1 | -1.4 | 57.0 | -61.6 |
| 300 | -74.8 | -46.6 | 1.2 | -3.1 | -22.9 | -0.9 | 56.3 | -58.9 |
| 301 | -84.7 | -56.9 | 2.0 | -3.0 | -27.5 | -1.9 | 66.3 | -63.8 |
| 302 | -69.2 | -45.6 | 1.3 | -1.8 | -25.3 | -1.7 | 63.5 | -59.6 |
| 303 | -77.1 | -52.2 | 2.0 | -2.3 | -27.5 | -2.0 | 66.0 | -61.1 |
| 304 | -76.3 | -58.4 | 2.6 | -2.6 | -24.5 | -1.2 | 68.8 | -60.9 |
| 305 | -77.6 | -63.3 | 2.8 | -2.6 | -25.4 | -1.9 | 71.5 | -58.8 |
| 306 | -78.2 | -40.2 | 2.3 | -2.6 | -25.7 | -1.3 | 49.5 | -60.3 |
| 307 | -76.9 | -63.9 | 2.0 | -2.9 | -24.7 | -0.7 | 74.2 | -60.9 |
| 308 | -75.7 | -50.8 | 2.8 | -2.6 | -25.7 | -0.7 | 60.1 | -58.7 |
| 309 | -76.8 | -65.3 | 1.3 | -2.6 | -24.5 | -0.8 | 73.9 | -58.8 |
| 310 | -77.5 | -35.1 | 2.3 | -3.1 | -25.9 | -1.2 | 45.3 | -59.8 |
| 311 | -75.0 | -45.5 | 0.3 | -2.1 | -26.1 | -1.4 | 58.7 | -58.8 |
| 312 | -73.9 | -43.9 | 0.1 | -2.9 | -23.2 | -0.7 | 55.2 | -58.6 |
| 313 | -77.7 | -40.8 | 0.0 | -3.0 | -23.5 | -1.0 | 50.7 | -60.2 |
| 314 | -68.6 | -43.0 | 1.5 | -3.1 | -22.6 | -1.0 | 53.8 | -54.2 |
| 315 | -73.0 | -57.6 | 0.3 | -2.6 | -23.8 | -1.1 | 71.2 | -59.5 |
| 316 | -74.3 | -59.9 | 1.3 | -3.3 | -23.6 | -1.2 | 68.8 | -56.4 |
| 317 | -71.2 | -57.4 | 4.7 | -3.4 | -23.6 | -1.4 | 67.7 | -57.8 |
| 318 | -76.0 | -63.7 | 1.5 | -3.8 | -22.7 | -1.1 | 70.6 | -56.8 |
| 319 | -64.7 | -50.2 | 0.9 | -2.2 | -22.7 | -1.2 | 62.7 | -52.0 |
| 320 | -68.2 | -56.7 | 1.3 | -2.4 | -20.8 | -1.1 | 65.5 | -53.9 |
| 321 | -78.5 | -46.9 | 2.4 | -3.3 | -24.4 | -1.2 | 55.6 | -60.7 |
| 322 | -75.8 | -55.0 | -0.5 | -3.6 | -21.9 | -1.0 | 62.0 | -55.9 |
| 323 | -74.6 | -44.8 | 2.2 | -2.5 | -25.0 | -2.2 | 55.2 | -57.6 |
| 324 | -74.2 | -51.3 | 2.6 | -2.9 | -23.0 | -1.1 | 56.6 | -55.2 |
| 325 | -67.9 | -44.0 | 0.9 | -2.6 | -22.6 | -1.3 | 56.4 | -54.6 |
| 326 | -67.9 | -46.0 | 2.5 | -2.7 | -23.5 | -1.4 | 59.8 | -56.8 |
| 327 | -75.4 | -30.0 | 0.4 | -2.6 | -24.8 | -1.7 | 39.3 | -56.0 |
| 328 | -73.7 | -35.1 | 1.4 | -2.5 | -23.0 | -1.1 | 46.2 | -59.5 |
| 329 | -70.8 | -33.2 | 2.1 | -2.3 | -25.3 | -1.5 | 50.1 | -60.6 |
| 330 | -68.9 | -52.6 | 3.6 | -2.3 | -22.4 | -1.1 | 64.8 | -58.8 |
| 331 | -68.4 | -50.4 | 0.6 | -1.9 | -23.6 | -1.0 | 66.5 | -58.6 |
| 332 | -68.2 | -60.4 | 1.1 | -3.5 | -21.1 | -1.3 | 67.1 | -50.1 |
| 333 | -62.7 | -45.4 | 3.5 | -3.0 | -19.7 | -1.0 | 53.5 | -50.5 |
| 334 | -76.5 | -63.6 | 1.8 | -3.1 | -23.2 | -0.8 | 68.9 | -56.5 |
| 335 | -66.2 | -71.5 | 1.9 | -3.5 | -20.6 | -1.7 | 81.3 | -52.1 |
| 336 | -70.5 | -41.4 | 1.9 | -3.0 | -22.4 | -1.2 | 54.6 | -58.8 |
| 337 | -67.4 | -47.4 | 3.3 | -2.6 | -21.8 | -2.1 | 60.4 | -57.2 |
| 338 | -69.6 | -66.7 | 2.3 | -3.6 | -20.3 | -0.9 | 75.0 | -55.5 |
| 339 | -78.6 | -53.7 | 1.9 | -2.4 | -25.7 | -1.3 | 64.7 | -62.2 |
| 340 | -72.3 | -34.8 | 2.1 | -2.7 | -23.6 | -1.9 | 45.8 | -57.2 |
| 341 | -76.1 | -39.8 | 2.0 | -3.1 | -26.0 | -1.6 | 50.3 | -58.0 |
| 342 | -68.6 | -61.9 | 0.6 | -2.9 | -21.2 | -1.0 | 72.8 | -55.1 |
| 343 | -75.2 | -53.0 | 0.7 | -3.0 | -23.5 | -2.9 | 66.7 | -60.2 |
| 344 | -70.0 | -53.7 | 1.8 | -3.4 | -23.7 | -1.0 | 64.9 | -54.9 |
| 345 | -76.7 | -42.0 | 1.6 | -3.5 | -24.8 | -0.9 | 53.3 | -60.2 |
| 346 | -77.2 | -62.8 | 1.3 | -2.9 | -24.7 | -0.9 | 75.9 | -63.2 |
| 347 | -74.8 | -54.7 | 1.5 | -2.4 | -27.4 | -1.1 | 69.2 | -59.9 |
| 348 | -69.5 | -63.9 | 1.6 | -3.5 | -22.2 | -1.6 | 71.6 | -51.6 |
| 349 | -81.4 | -53.6 | 2.1 | -3.2 | -26.8 | -1.4 | 64.1 | -62.6 |
| 350 | -68.7 | -56.6 | 2.2 | -2.7 | -24.0 | -2.2 | 70.8 | -56.1 |
| 351 | -64.6 | -59.5 | 1.2 | -2.9 | -20.0 | -1.6 | 71.9 | -53.7 |
| 352 | -78.0 | -57.8 | 1.8 | -3.0 | -24.2 | -1.0 | 66.7 | -60.5 |
| 353 | -76.3 | -52.2 | 2.2 | -2.6 | -25.1 | -2.1 | 65.4 | -61.9 |
| 354 | -81.2 | -54.4 | 1.0 | -3.1 | -26.9 | -1.1 | 65.5 | -62.1 |
| 355 | -76.8 | -46.2 | 0.7 | -3.2 | -25.0 | -0.8 | 59.4 | -61.8 |
| 356 | -77.0 | -43.1 | 0.9 | -2.9 | -24.4 | -0.6 | 54.2 | -61.1 |
| 357 | -85.2 | -65.3 | 0.8 | -3.5 | -24.8 | -1.1 | 71.2 | -62.6 |
| 358 | -75.5 | -54.1 | 1.7 | -2.7 | -23.9 | -1.4 | 62.9 | -57.8 |
| 359 | -82.7 | -42.9 | 1.8 | -3.2 | -25.6 | -1.6 | 50.2 | -61.3 |
| 360 | -77.4 | -51.1 | 3.9 | -3.1 | -24.6 | -1.1 | 57.6 | -59.0 |
| 361 | -74.5 | -45.1 | 0.7 | -3.4 | -23.6 | -1.9 | 52.2 | -53.4 |
| 362 | -79.7 | -39.7 | 1.0 | -3.2 | -25.3 | -1.2 | 48.3 | -59.4 |
| 363 | -80.6 | -53.6 | 3.7 | -2.9 | -24.8 | -1.0 | 60.7 | -62.7 |
| 364 | -85.5 | -44.7 | 0.4 | -2.6 | -27.5 | -1.7 | 55.5 | -64.8 |
| 365 | -85.2 | -66.4 | 0.2 | -3.0 | -26.9 | -1.7 | 71.6 | -59.0 |
| 366 | -66.6 | -55.2 | 2.7 | -2.9 | -21.5 | -0.3 | 67.5 | -57.0 |
| 367 | -71.5 | -54.2 | -0.7 | -3.5 | -22.3 | -0.9 | 64.6 | -54.5 |
| 368 | -73.5 | -48.5 | 1.6 | -2.8 | -23.7 | -1.1 | 59.9 | -58.9 |
| 369 | -77.0 | -50.7 | 1.1 | -3.2 | -23.6 | -1.2 | 58.6 | -58.0 |
| 370 | -70.9 | -44.5 | 1.2 | -3.6 | -20.7 | -0.6 | 51.7 | -54.5 |
| 371 | -76.9 | -48.6 | -0.7 | -3.2 | -23.1 | -1.6 | 56.1 | -55.8 |
| 372 | -86.5 | -34.6 | 1.8 | -3.0 | -28.9 | -2.0 | 42.2 | -62.0 |
| 373 | -73.6 | -52.5 | 2.1 | -3.0 | -22.6 | -1.3 | 59.1 | -55.4 |
| 374 | -73.4 | -40.5 | 1.5 | -3.7 | -24.2 | -1.7 | 53.6 | -58.4 |
| 375 | -66.2 | -53.8 | 1.1 | -3.0 | -21.8 | -1.2 | 61.7 | -49.3 |
| 376 | -75.1 | -48.1 | 1.6 | -3.0 | -23.4 | -1.0 | 58.1 | -59.2 |
| 377 | -74.2 | -42.9 | 1.8 | -2.7 | -24.4 | -0.8 | 54.8 | -60.0 |
| 378 | -68.1 | -75.6 | 1.2 | -3.1 | -22.0 | -1.4 | 87.3 | -54.6 |
| 379 | -72.0 | -33.3 | 2.4 | -2.8 | -23.7 | -1.4 | 44.2 | -57.5 |
| 380 | -89.9 | -49.9 | 2.7 | -3.7 | -27.4 | -1.9 | 54.3 | -64.0 |
| 381 | -69.3 | -60.4 | 2.5 | -3.5 | -22.7 | -1.5 | 70.3 | -54.0 |
| 382 | -69.2 | -45.3 | 1.1 | -2.9 | -21.4 | -0.6 | 58.0 | -58.0 |
| 383 | -72.9 | -44.4 | 1.4 | -3.0 | -23.5 | -1.7 | 56.5 | -58.2 |
| 384 | -82.0 | -39.0 | 1.0 | -2.8 | -25.8 | -1.1 | 50.7 | -64.9 |
| 385 | -71.4 | -35.0 | 0.9 | -2.9 | -23.2 | -0.9 | 48.8 | -59.2 |
| 386 | -72.6 | -42.6 | 1.1 | -3.3 | -23.2 | -0.6 | 53.0 | -57.1 |
| 387 | -65.1 | -37.0 | 3.1 | -3.2 | -22.5 | -1.5 | 49.0 | -52.9 |
| 388 | -79.0 | -58.4 | 0.7 | -3.2 | -25.8 | -1.3 | 67.5 | -58.5 |
| 389 | -78.7 | -53.0 | 2.3 | -2.7 | -24.4 | -1.2 | 64.1 | -63.9 |
| 390 | -73.2 | -46.2 | 0.9 | -3.0 | -22.2 | -0.7 | 58.7 | -60.7 |
| 391 | -73.2 | -54.6 | 3.9 | -3.5 | -24.2 | -1.8 | 64.9 | -57.9 |
| 392 | -70.3 | -45.6 | 1.5 | -2.4 | -22.6 | -1.5 | 59.5 | -59.2 |
| 393 | -80.8 | -55.4 | 2.3 | -4.0 | -24.7 | -2.0 | 62.8 | -59.8 |
| 394 | -76.9 | -40.3 | 1.6 | -3.1 | -24.8 | -1.3 | 53.2 | -62.2 |
| 395 | -67.5 | -35.3 | 0.7 | -2.9 | -23.0 | -1.1 | 54.9 | -60.9 |
| 396 | -74.2 | -48.8 | 1.1 | -3.2 | -23.4 | -1.5 | 59.2 | -57.6 |
| 397 | -71.0 | -48.9 | 2.0 | -3.8 | -23.0 | -1.1 | 59.6 | -55.9 |
| 398 | -70.4 | -43.3 | 3.2 | -2.6 | -22.6 | -1.4 | 55.7 | -59.4 |
| 399 | -73.8 | -56.5 | 2.8 | -3.9 | -23.1 | -1.4 | 67.3 | -59.1 |
| 400 | -77.1 | -64.0 | 0.3 | -3.6 | -21.7 | -1.8 | 69.7 | -56.0 |
| 401 | -75.9 | -61.3 | 1.3 | -3.3 | -21.5 | -1.2 | 71.8 | -61.6 |
| 402 | -62.2 | -33.9 | 3.8 | -2.4 | -22.2 | -1.4 | 49.5 | -55.5 |
| 403 | -67.5 | -59.0 | 1.1 | -4.0 | -19.5 | -1.1 | 68.3 | -53.3 |
| 404 | -65.9 | -50.7 | 1.5 | -2.9 | -21.1 | -1.1 | 61.3 | -52.9 |
| 405 | -65.9 | -55.7 | 2.7 | -3.4 | -18.5 | -1.1 | 61.7 | -51.7 |
| 406 | -69.8 | -46.8 | 1.2 | -2.6 | -22.7 | -1.0 | 58.6 | -56.5 |
| 407 | -73.7 | -54.4 | 1.8 | -2.8 | -24.2 | -1.2 | 65.3 | -58.2 |
| 408 | -70.9 | -41.2 | 1.5 | -3.3 | -24.4 | -2.3 | 53.7 | -54.8 |
| 409 | -65.6 | -51.7 | 2.6 | -2.8 | -22.4 | -0.7 | 64.8 | -55.3 |
| 410 | -74.1 | -39.6 | 1.1 | -3.0 | -23.7 | -1.4 | 49.2 | -56.7 |
| 411 | -67.7 | -47.4 | 2.1 | -2.5 | -22.2 | -2.3 | 58.0 | -53.4 |
| 412 | -63.3 | -54.1 | 2.3 | -2.8 | -22.1 | -1.1 | 65.3 | -50.9 |
| 413 | -74.7 | -60.9 | 0.2 | -2.8 | -26.2 | -2.3 | 71.2 | -53.9 |
| 414 | -68.3 | -48.8 | 2.1 | -2.6 | -25.2 | -1.4 | 59.4 | -51.6 |
| 415 | -80.8 | -39.2 | 1.3 | -3.0 | -26.6 | -1.7 | 48.5 | -60.1 |
| 416 | -75.1 | -46.9 | 2.3 | -2.3 | -25.8 | -2.6 | 58.3 | -58.1 |
| 417 | -84.0 | -49.3 | 0.3 | -2.5 | -30.6 | -3.0 | 60.8 | -59.8 |
| 418 | -80.7 | -36.6 | 1.5 | -2.7 | -29.6 | -1.6 | 48.0 | -59.8 |
| 419 | -76.2 | -55.0 | 0.6 | -2.9 | -24.5 | -2.1 | 62.4 | -54.9 |
| 420 | -78.6 | -42.1 | 1.0 | -2.6 | -26.3 | -1.4 | 48.7 | -55.9 |
| 421 | -72.6 | -37.9 | 2.0 | -2.3 | -24.5 | -1.5 | 47.0 | -55.4 |
| 422 | -82.0 | -56.6 | 0.6 | -3.4 | -26.9 | -1.8 | 64.9 | -58.7 |
| 423 | -73.3 | -51.9 | 1.3 | -2.6 | -24.6 | -2.0 | 63.4 | -57.0 |
| 424 | -74.2 | -52.4 | 1.6 | -2.7 | -24.4 | -2.4 | 61.9 | -55.8 |
| 425 | -70.4 | -48.3 | 0.7 | -3.6 | -22.2 | -1.9 | 56.2 | -51.2 |
| 426 | -68.3 | -58.1 | 0.7 | -3.0 | -22.0 | -0.8 | 66.7 | -51.7 |
| 427 | -85.9 | -40.7 | 1.0 | -2.6 | -30.9 | -2.2 | 50.6 | -61.0 |
| 428 | -77.5 | -56.4 | 1.4 | -3.3 | -25.6 | -1.6 | 64.0 | -55.9 |
| 429 | -75.0 | -50.7 | 1.0 | -2.8 | -25.7 | -2.1 | 61.2 | -55.9 |
| 430 | -80.1 | -50.0 | 1.1 | -2.7 | -27.3 | -2.3 | 58.9 | -57.8 |
| 431 | -69.7 | -52.2 | 2.8 | -2.6 | -24.1 | -1.8 | 64.0 | -55.9 |
| 432 | -80.8 | -49.6 | 1.6 | -2.0 | -29.9 | -2.2 | 62.4 | -61.2 |
| 433 | -84.5 | -62.1 | 0.9 | -3.3 | -25.9 | -1.9 | 68.9 | -61.1 |
| 434 | -71.0 | -49.5 | 1.2 | -2.6 | -24.1 | -1.0 | 61.8 | -56.9 |
| 435 | -79.4 | -48.9 | 0.7 | -3.0 | -26.6 | -2.0 | 56.8 | -56.3 |
| 436 | -82.8 | -47.7 | 1.2 | -2.8 | -26.8 | -2.3 | 53.9 | -58.3 |
| 437 | -76.6 | -47.8 | 1.1 | -2.9 | -24.4 | -2.1 | 56.3 | -56.8 |
| 438 | -74.3 | -42.9 | 1.5 | -2.5 | -24.7 | -1.7 | 53.2 | -57.1 |
| 439 | -76.4 | -39.6 | 1.2 | -3.4 | -26.4 | -2.0 | 50.3 | -56.5 |
| 440 | -86.7 | -52.2 | 0.9 | -3.2 | -30.3 | -1.7 | 60.6 | -60.7 |
| 441 | -76.2 | -37.6 | 2.4 | -3.2 | -26.0 | -2.4 | 47.8 | -57.3 |
| 442 | -79.4 | -42.7 | 0.5 | -3.2 | -26.3 | -1.2 | 53.8 | -60.4 |
| 443 | -78.1 | -58.4 | -0.1 | -2.9 | -26.7 | -1.4 | 71.1 | -59.8 |
| 444 | -72.0 | -55.6 | 1.3 | -2.5 | -25.6 | -1.9 | 69.6 | -57.3 |
| 445 | -74.1 | -43.8 | 1.1 | -2.4 | -25.9 | -1.5 | 57.7 | -59.3 |
| 446 | -70.6 | -33.5 | 0.5 | -2.4 | -24.0 | -1.8 | 47.6 | -56.9 |
| 447 | -74.8 | -51.7 | 3.1 | -2.9 | -25.1 | -2.0 | 60.3 | -56.6 |
| 448 | -85.5 | -48.5 | 0.7 | -2.8 | -27.1 | -2.1 | 56.6 | -62.2 |
| 449 | -79.9 | -58.9 | 1.7 | -3.0 | -26.6 | -1.6 | 67.9 | -59.4 |
| 450 | -86.6 | -62.0 | -0.2 | -3.6 | -26.5 | -2.3 | 69.3 | -61.4 |
| 451 | -75.4 | -40.8 | 0.6 | -2.8 | -25.4 | -1.5 | 51.3 | -56.7 |
| 452 | -82.4 | -53.0 | 0.5 | -2.1 | -26.5 | -2.1 | 61.6 | -60.8 |
| 453 | -74.0 | -51.4 | 1.1 | -3.8 | -25.2 | -1.3 | 59.7 | -53.2 |
| 454 | -76.8 | -46.8 | -0.7 | -2.7 | -26.4 | -1.5 | 59.6 | -58.4 |
| 455 | -72.4 | -52.4 | 1.7 | -2.7 | -24.1 | -1.5 | 61.7 | -55.1 |
| 456 | -77.9 | -41.4 | 1.5 | -2.5 | -29.4 | -2.3 | 57.6 | -61.5 |
| 457 | -80.5 | -60.5 | 0.9 | -3.1 | -28.0 | -1.9 | 68.3 | -56.1 |
| 458 | -69.2 | -33.3 | 1.1 | -2.6 | -24.3 | -1.4 | 46.2 | -54.9 |
| 459 | -68.7 | -45.8 | -0.1 | -2.6 | -24.2 | -1.0 | 63.2 | -58.3 |
| 460 | -76.0 | -45.6 | 0.3 | -3.0 | -25.0 | -1.3 | 55.3 | -56.6 |
| 461 | -80.3 | -60.0 | 0.2 | -3.1 | -26.9 | -2.6 | 68.0 | -55.7 |
| 462 | -75.0 | -58.2 | 2.6 | -2.9 | -22.4 | -1.5 | 65.4 | -58.0 |
| 463 | -84.7 | -51.9 | 0.6 | -2.4 | -27.5 | -2.1 | 60.1 | -61.5 |
| 464 | -90.0 | -48.5 | 1.5 | -2.9 | -31.2 | -2.5 | 56.9 | -63.2 |
| 465 | -74.4 | -61.9 | -1.7 | -3.1 | -23.5 | -3.0 | 72.7 | -53.9 |
| 466 | -71.6 | -47.6 | 2.0 | -2.7 | -22.0 | -2.1 | 59.6 | -58.7 |
| 467 | -79.7 | -52.3 | 1.9 | -3.0 | -28.5 | -1.8 | 62.3 | -58.2 |
| 468 | -71.6 | -72.8 | -1.7 | -2.9 | -20.7 | -1.4 | 83.5 | -55.6 |
| 469 | -67.7 | -56.1 | 1.6 | -3.1 | -19.9 | -1.2 | 65.9 | -54.9 |
| 470 | -76.3 | -54.2 | 3.1 | -3.3 | -23.6 | -1.9 | 62.8 | -59.2 |
| 471 | -71.7 | -54.1 | 3.5 | -2.5 | -22.7 | -2.0 | 63.3 | -57.2 |
| 472 | -66.1 | -46.6 | 1.7 | -2.5 | -21.4 | -1.5 | 58.5 | -54.4 |
| 473 | -91.2 | -70.7 | 1.3 | -2.8 | -29.6 | -2.1 | 75.2 | -62.5 |
| 474 | -77.6 | -45.4 | 3.2 | -2.5 | -24.2 | -1.9 | 52.1 | -58.8 |
| 475 | -71.9 | -52.9 | 3.5 | -2.5 | -22.1 | -2.2 | 61.2 | -56.8 |
| 476 | -80.3 | -66.6 | 2.2 | -2.8 | -23.0 | -1.6 | 71.0 | -59.4 |
| 477 | -82.7 | -58.8 | 2.2 | -2.8 | -25.2 | -2.4 | 63.5 | -59.1 |
| 478 | -81.7 | -62.3 | 1.6 | -2.9 | -25.8 | -2.5 | 67.8 | -57.6 |
| 479 | -74.0 | -48.4 | 1.8 | -2.7 | -23.5 | -1.2 | 60.2 | -60.2 |
| 480 | -75.5 | -51.9 | 2.3 | -2.8 | -24.3 | -1.7 | 59.9 | -56.9 |
| 481 | -83.4 | -55.2 | 1.5 | -2.8 | -26.4 | -2.3 | 60.7 | -58.9 |
| 482 | -87.3 | -49.8 | 0.2 | -3.1 | -30.5 | -1.6 | 57.1 | -59.6 |
| 483 | -78.7 | -45.8 | 1.6 | -2.9 | -24.2 | -1.5 | 51.0 | -56.8 |
| 484 | -82.7 | -47.1 | 2.4 | -3.1 | -26.2 | -2.3 | 52.8 | -59.2 |
| 485 | -72.8 | -52.7 | 1.7 | -2.8 | -23.5 | -2.0 | 63.8 | -57.3 |
| 486 | -77.4 | -49.2 | 1.5 | -3.1 | -23.7 | -1.9 | 57.7 | -58.6 |
| 487 | -76.0 | -44.7 | 3.4 | -2.9 | -24.9 | -2.3 | 52.8 | -57.4 |
| 488 | -80.9 | -59.1 | 1.8 | -2.8 | -24.7 | -2.1 | 64.6 | -58.5 |
| 489 | -79.3 | -48.8 | 2.0 | -2.4 | -25.7 | -2.0 | 54.9 | -57.4 |
| 490 | -77.9 | -60.0 | 1.3 | -2.8 | -24.0 | -1.9 | 66.9 | -57.4 |
| 491 | -90.9 | -51.3 | 1.2 | -3.0 | -30.4 | -1.6 | 56.8 | -62.5 |
| 492 | -81.0 | -48.2 | 2.0 | -2.9 | -24.1 | -2.2 | 55.6 | -61.3 |
| 493 | -100.3 | -54.6 | 0.7 | -3.2 | -33.7 | -1.8 | 58.2 | -65.7 |
| 494 | -72.6 | -48.3 | 1.7 | -2.4 | -22.7 | -1.7 | 59.0 | -58.1 |
| 495 | -82.5 | -49.2 | 1.9 | -3.0 | -26.6 | -2.2 | 57.5 | -61.0 |
| 496 | -96.4 | -59.7 | 1.0 | -3.3 | -33.2 | -3.2 | 68.6 | -66.5 |
| 497 | -74.8 | -61.2 | 1.6 | -3.1 | -24.0 | -2.0 | 71.0 | -57.1 |
| 498 | -76.7 | -57.4 | 2.7 | -3.0 | -22.5 | -2.1 | 63.4 | -57.9 |
| 499 | -79.9 | -62.8 | 1.3 | -3.0 | -26.8 | -0.7 | 72.7 | -60.7 |
| 500 | -73.2 | -43.0 | 2.6 | -2.8 | -22.3 | -1.9 | 51.9 | -57.6 |
| MAX | **-100.3** | **-82.3** | **-1.7** | **-4.0** | **-33.7** | **-5.0** | **31.1** | **-67.8** |
| MIN | **-54.6** | **-19.5** | **4.7** | **-1.8** | **-17.0** | **-0.1** | **88.6** | **-43.3** |
| AVG | **-74.9** | **-51.1** | **1.6** | **-2.9** | **-23.9** | **-1.7** | **59.6** | **-56.5** |
| SDV | **6.9** | **8.9** | **1.0** | **0.4** | **2.5** | **0.9** | **8.5** | **4.9** |

**Table S6** Post simulation MM-GBSA based Binding free energy (ΔG Bind in kcal/mol) for the 6JQR-CHEMBL4444839-analog Complex

| Time (in ns) | ΔG_bind_ Total | ΔG_bind_ Coulomb | ΔG_bind_ Covalent | ΔG_bind_ Hbond | ΔG_bind_ Lipo | ΔG_bind_ Packing | ΔG_bind_ Solv_GB | ΔG_bind_ vdW |
| --- | --- | --- | --- | --- | --- | --- | --- | --- |
| 1 | -95.6 | -50.3 | 4.0 | -3.6 | -28.1 | -2.3 | 51.1 | -66.5 |
| 2 | -103.1 | -43.7 | 4.1 | -1.9 | -33.8 | -4.3 | 51.8 | -75.3 |
| 3 | -88.3 | -70.4 | 5.6 | -2.1 | -32.3 | -5.0 | 83.1 | -67.2 |
| 4 | -103.4 | -56.6 | 3.7 | -2.3 | -36.1 | -4.1 | 68.2 | -76.2 |
| 5 | -91.6 | -46.1 | 3.8 | -2.1 | -33.9 | -4.4 | 60.2 | -69.2 |
| 6 | -85.3 | -42.5 | 2.0 | -1.9 | -27.6 | -1.6 | 52.7 | -66.4 |
| 7 | -79.6 | -47.3 | 0.2 | -2.0 | -25.7 | -4.2 | 62.2 | -62.9 |
| 8 | -83.7 | -34.5 | 0.8 | -2.4 | -24.3 | -1.5 | 46.8 | -68.7 |
| 9 | -72.1 | -57.4 | 4.0 | -2.0 | -23.7 | -1.7 | 70.4 | -61.9 |
| 10 | -87.0 | -30.8 | 6.3 | -2.1 | -29.7 | -2.9 | 41.9 | -69.8 |
| 11 | -81.8 | -65.0 | 5.8 | -2.0 | -26.7 | -3.6 | 76.2 | -66.6 |
| 12 | -75.3 | -37.9 | 2.2 | -2.1 | -25.8 | -0.6 | 52.5 | -63.7 |
| 13 | -84.9 | -76.7 | 1.8 | -2.6 | -28.1 | -5.4 | 82.7 | -56.4 |
| 14 | -78.7 | -45.4 | 3.6 | -2.1 | -29.2 | -5.1 | 60.2 | -60.6 |
| 15 | -79.1 | -47.4 | 4.5 | -1.9 | -25.9 | -2.8 | 61.3 | -66.8 |
| 16 | -83.6 | -54.1 | 4.4 | -2.3 | -27.6 | -3.6 | 64.0 | -64.4 |
| 17 | -75.3 | -43.3 | 1.8 | -1.2 | -26.3 | -2.2 | 59.7 | -63.9 |
| 18 | -80.5 | -30.8 | 2.7 | -1.7 | -27.3 | -4.2 | 45.0 | -64.2 |
| 19 | -88.9 | -43.5 | 4.1 | -2.0 | -30.1 | -4.8 | 53.5 | -66.2 |
| 20 | -86.6 | -45.7 | 1.7 | -2.3 | -28.3 | -4.0 | 57.2 | -65.3 |
| 21 | -81.6 | -38.4 | 2.2 | -2.1 | -28.3 | -3.9 | 53.0 | -64.3 |
| 22 | -95.3 | -31.6 | 0.1 | -2.2 | -31.4 | -4.3 | 42.1 | -68.0 |
| 23 | -88.3 | -39.1 | 2.8 | -2.3 | -28.8 | -2.0 | 49.5 | -68.3 |
| 24 | -81.2 | -46.8 | 4.2 | -2.0 | -26.4 | -2.8 | 55.8 | -63.3 |
| 25 | -81.4 | -45.5 | 5.3 | -2.1 | -27.0 | -2.7 | 56.0 | -65.3 |
| 26 | -91.0 | -40.7 | 4.6 | -1.7 | -33.0 | -2.0 | 51.0 | -69.1 |
| 27 | -87.2 | -44.9 | 7.8 | -1.7 | -32.9 | -2.3 | 55.1 | -68.4 |
| 28 | -76.8 | -25.5 | 5.7 | -1.9 | -29.1 | -3.2 | 41.3 | -64.3 |
| 29 | -85.5 | -60.9 | 2.6 | -2.3 | -29.2 | -3.2 | 68.6 | -61.1 |
| 30 | -90.9 | -36.1 | 3.1 | -2.2 | -31.8 | -2.9 | 44.4 | -65.4 |
| 31 | -76.3 | -42.8 | 3.5 | -2.2 | -25.4 | -1.7 | 50.0 | -57.6 |
| 32 | -82.2 | -35.9 | 5.3 | -2.1 | -29.5 | -4.3 | 42.1 | -57.8 |
| 33 | -67.5 | -34.8 | 4.1 | -2.1 | -24.1 | -3.8 | 44.5 | -51.4 |
| 34 | -82.8 | -55.0 | 5.4 | -1.9 | -29.7 | -3.1 | 65.0 | -63.5 |
| 35 | -68.0 | -49.3 | 3.5 | -2.0 | -24.3 | -2.3 | 62.1 | -55.8 |
| 36 | -86.0 | -28.5 | 3.5 | -1.9 | -30.2 | -2.9 | 42.9 | -69.0 |
| 37 | -85.0 | -45.6 | 5.3 | -2.2 | -29.9 | -3.0 | 57.9 | -67.5 |
| 38 | -78.7 | -43.0 | 11.5 | -2.6 | -28.0 | -2.3 | 48.7 | -63.1 |
| 39 | -75.1 | -40.3 | 5.5 | -2.3 | -26.4 | -1.9 | 53.9 | -63.6 |
| 40 | -83.0 | -46.3 | 2.7 | -2.0 | -28.3 | -2.6 | 53.8 | -60.3 |
| 41 | -87.8 | -58.7 | 5.3 | -2.3 | -30.5 | -5.3 | 68.0 | -64.3 |
| 42 | -82.7 | -58.4 | 2.9 | -2.1 | -27.4 | -1.6 | 69.9 | -66.0 |
| 43 | -90.8 | -33.1 | 1.3 | -2.6 | -28.4 | -4.6 | 44.1 | -67.5 |
| 44 | -84.7 | -40.1 | 3.9 | -1.9 | -29.0 | -2.5 | 50.8 | -65.8 |
| 45 | -82.9 | -38.6 | 4.5 | -2.1 | -29.2 | -1.6 | 49.2 | -65.1 |
| 46 | -84.4 | -41.0 | 7.8 | -2.2 | -31.5 | -2.1 | 50.0 | -65.4 |
| 47 | -81.8 | -47.0 | 6.5 | -1.8 | -30.1 | -2.8 | 57.8 | -64.4 |
| 48 | -83.2 | -43.6 | 3.4 | -1.8 | -28.7 | -3.0 | 58.8 | -68.3 |
| 49 | -87.2 | -29.4 | 4.1 | -1.7 | -31.2 | -3.3 | 42.4 | -68.1 |
| 50 | -84.2 | -37.8 | 4.7 | -2.0 | -29.0 | -3.5 | 48.2 | -64.9 |
| 51 | -82.9 | -44.6 | 5.4 | -2.3 | -29.1 | -2.1 | 57.8 | -68.1 |
| 52 | -69.2 | -48.0 | 2.2 | -1.9 | -26.5 | -1.2 | 66.9 | -60.7 |
| 53 | -77.9 | -31.7 | 3.0 | -1.9 | -28.7 | -2.0 | 48.0 | -64.7 |
| 54 | -88.1 | -43.5 | 4.6 | -2.0 | -31.0 | -2.7 | 52.5 | -66.1 |
| 55 | -85.5 | -44.7 | 4.3 | -1.0 | -30.4 | -4.9 | 54.8 | -63.5 |
| 56 | -76.1 | -32.7 | 6.6 | -2.3 | -26.3 | -2.2 | 45.8 | -65.0 |
| 57 | -78.6 | -31.6 | 5.2 | -1.7 | -28.5 | -2.6 | 41.7 | -61.0 |
| 58 | -80.5 | -54.7 | 2.1 | -2.1 | -27.3 | -1.8 | 64.7 | -61.5 |
| 59 | -80.8 | -45.0 | 1.8 | -2.2 | -28.0 | -3.3 | 57.2 | -61.3 |
| 60 | -75.9 | -35.8 | 5.7 | -2.1 | -26.7 | -3.0 | 48.6 | -62.6 |
| 61 | -82.8 | -35.9 | 5.5 | -2.2 | -27.5 | -2.0 | 45.7 | -66.4 |
| 62 | -88.1 | -34.8 | 4.1 | -1.7 | -31.0 | -4.8 | 45.7 | -65.6 |
| 63 | -85.9 | -32.2 | 3.3 | -2.1 | -32.3 | -3.2 | 45.9 | -65.2 |
| 64 | -81.9 | -53.5 | 4.6 | -2.6 | -27.0 | -4.2 | 61.7 | -60.9 |
| 65 | -80.5 | -40.2 | 4.1 | -2.4 | -27.1 | -2.2 | 51.6 | -64.3 |
| 66 | -85.4 | -36.2 | 4.2 | -2.1 | -30.4 | -3.2 | 47.2 | -64.9 |
| 67 | -84.8 | -42.8 | 5.6 | -2.4 | -30.3 | -2.5 | 52.5 | -65.0 |
| 68 | -81.3 | -30.1 | 3.8 | -2.2 | -29.2 | -2.6 | 42.5 | -63.6 |
| 69 | -81.7 | -41.4 | 2.3 | -2.2 | -26.6 | -2.6 | 47.5 | -58.7 |
| 70 | -76.3 | -48.2 | 5.3 | -1.9 | -26.7 | -4.5 | 62.9 | -63.2 |
| 71 | -76.6 | -43.0 | 4.3 | -2.0 | -28.3 | -1.8 | 55.5 | -61.2 |
| 72 | -84.9 | -39.6 | 4.9 | -2.2 | -32.5 | -3.0 | 51.8 | -64.3 |
| 73 | -77.1 | -52.2 | 7.6 | -2.0 | -29.0 | -4.1 | 61.6 | -59.0 |
| 74 | -81.1 | -49.3 | 2.1 | -1.9 | -28.0 | -3.8 | 62.0 | -62.1 |
| 75 | -81.9 | -40.3 | 2.7 | -2.1 | -30.1 | -2.6 | 54.7 | -64.0 |
| 76 | -76.8 | -44.4 | 4.1 | -2.1 | -24.7 | -2.3 | 53.6 | -61.0 |
| 77 | -74.1 | -38.1 | 2.5 | -2.1 | -24.6 | -2.9 | 50.1 | -59.1 |
| 78 | -80.6 | -36.9 | 3.6 | -1.6 | -30.9 | -3.8 | 56.5 | -67.6 |
| 79 | -79.6 | -26.6 | 4.7 | -2.0 | -30.3 | -2.8 | 42.4 | -65.1 |
| 80 | -89.1 | -41.3 | 3.3 | -2.2 | -29.5 | -1.9 | 52.3 | -69.7 |
| 81 | -87.1 | -48.8 | 2.8 | -2.3 | -31.3 | -2.6 | 62.2 | -67.1 |
| 82 | -80.7 | -23.1 | 2.4 | -2.1 | -27.7 | -2.0 | 33.1 | -61.2 |
| 83 | -84.0 | -36.2 | 3.1 | -1.9 | -31.0 | -2.0 | 48.6 | -64.5 |
| 84 | -84.0 | -57.2 | 3.7 | -1.8 | -30.0 | -2.5 | 70.8 | -67.0 |
| 85 | -81.8 | -48.4 | 1.4 | -2.4 | -27.5 | -2.9 | 61.8 | -63.7 |
| 86 | -88.4 | -50.1 | 5.3 | -2.5 | -29.1 | -1.4 | 60.5 | -71.0 |
| 87 | -90.4 | -51.0 | 2.7 | -2.4 | -30.0 | -2.6 | 63.6 | -70.8 |
| 88 | -83.0 | -30.2 | 3.3 | -2.8 | -28.0 | -1.7 | 39.1 | -62.7 |
| 89 | -89.3 | -39.2 | 3.6 | -2.6 | -25.7 | -1.6 | 44.1 | -67.8 |
| 90 | -79.4 | -38.3 | 3.9 | -2.5 | -24.2 | -2.0 | 49.6 | -65.8 |
| 91 | -87.3 | -60.6 | 6.8 | -2.7 | -27.5 | -2.8 | 66.0 | -66.4 |
| 92 | -78.4 | -36.6 | 4.6 | -2.3 | -27.6 | -3.6 | 48.0 | -60.9 |
| 93 | -78.0 | -36.2 | 4.4 | -1.8 | -30.9 | -5.0 | 58.1 | -66.5 |
| 94 | -86.7 | -45.0 | 4.1 | -1.9 | -31.8 | -3.0 | 57.1 | -66.2 |
| 95 | -78.7 | -50.8 | 3.0 | -2.0 | -27.0 | -3.2 | 61.1 | -59.7 |
| 96 | -87.8 | -49.5 | 2.5 | -1.7 | -28.8 | -1.9 | 56.2 | -64.6 |
| 97 | -80.6 | -45.1 | 5.9 | -1.8 | -28.6 | -3.7 | 55.4 | -62.6 |
| 98 | -82.5 | -69.2 | 2.4 | -1.6 | -26.0 | -2.9 | 75.8 | -61.1 |
| 99 | -84.5 | -39.6 | 4.3 | -1.7 | -29.1 | -2.5 | 51.8 | -67.7 |
| 100 | -87.1 | -43.8 | 2.7 | -1.9 | -30.0 | -2.2 | 55.6 | -67.5 |
| 101 | -67.7 | -42.6 | 4.9 | -1.9 | -27.0 | -1.3 | 61.7 | -61.6 |
| 102 | -83.3 | -41.8 | 2.8 | -2.5 | -29.1 | -2.7 | 57.3 | -67.3 |
| 103 | -86.9 | -43.1 | 3.8 | -2.7 | -27.9 | -5.2 | 52.4 | -64.3 |
| 104 | -79.7 | -47.4 | 3.0 | -1.8 | -26.7 | -2.0 | 57.4 | -62.2 |
| 105 | -74.4 | -35.4 | 3.5 | -2.1 | -26.3 | -2.8 | 52.6 | -63.9 |
| 106 | -85.4 | -57.8 | 3.5 | -2.2 | -27.3 | -2.6 | 66.9 | -66.0 |
| 107 | -81.3 | -40.5 | 4.1 | -2.1 | -27.6 | -4.5 | 54.4 | -65.2 |
| 108 | -79.7 | -44.9 | 5.8 | -2.4 | -25.4 | -3.8 | 57.2 | -66.1 |
| 109 | -83.8 | -55.3 | 3.9 | -2.6 | -27.9 | -2.6 | 65.4 | -64.6 |
| 110 | -93.4 | -48.2 | 2.0 | -2.2 | -30.9 | -5.1 | 60.5 | -69.6 |
| 111 | -81.5 | -52.7 | 3.6 | -2.0 | -25.8 | -4.6 | 66.6 | -66.6 |
| 112 | -84.2 | -33.5 | 2.8 | -2.1 | -28.1 | -5.3 | 45.6 | -63.8 |
| 113 | -86.5 | -34.4 | 5.2 | -2.1 | -28.9 | -3.6 | 46.6 | -69.3 |
| 114 | -74.8 | -51.9 | 5.8 | -2.4 | -24.6 | -5.4 | 65.8 | -62.2 |
| 115 | -87.0 | -43.9 | 3.2 | -2.9 | -31.3 | -2.3 | 55.7 | -65.6 |
| 116 | -82.1 | -42.1 | 3.9 | -3.2 | -31.9 | -4.0 | 58.2 | -63.1 |
| 117 | -84.1 | -54.0 | 6.8 | -2.5 | -29.2 | -2.6 | 63.0 | -65.6 |
| 118 | -84.1 | -46.3 | 4.4 | -2.3 | -30.4 | -5.3 | 63.8 | -67.9 |
| 119 | -79.4 | -50.4 | 3.0 | -2.0 | -26.8 | -4.1 | 66.5 | -65.6 |
| 120 | -81.8 | -35.7 | 4.5 | -2.1 | -29.6 | -4.0 | 54.7 | -69.6 |
| 121 | -84.5 | -50.6 | 2.0 | -2.1 | -29.6 | -4.4 | 68.0 | -67.7 |
| 122 | -78.8 | -37.6 | 2.2 | -2.4 | -27.1 | -2.9 | 50.8 | -61.9 |
| 123 | -74.2 | -39.4 | 2.4 | -2.4 | -25.6 | -2.3 | 54.4 | -61.2 |
| 124 | -84.2 | -48.9 | 2.9 | -2.1 | -30.3 | -5.5 | 61.7 | -61.9 |
| 125 | -83.7 | -44.0 | 2.0 | -2.1 | -27.5 | -2.6 | 55.7 | -65.2 |
| 126 | -85.0 | -43.8 | 4.4 | -2.4 | -28.9 | -3.0 | 53.3 | -64.5 |
| 127 | -76.9 | -24.2 | 2.8 | -2.1 | -29.6 | -2.3 | 44.2 | -65.5 |
| 128 | -81.2 | -30.2 | 2.1 | -2.6 | -27.5 | -1.5 | 45.1 | -66.6 |
| 129 | -81.9 | -37.9 | 2.5 | -2.6 | -26.6 | -3.8 | 52.7 | -66.1 |
| 130 | -81.8 | -41.9 | 0.3 | -2.5 | -25.1 | -1.6 | 55.7 | -66.6 |
| 131 | -79.4 | -42.2 | 3.4 | -2.1 | -25.8 | -1.8 | 55.9 | -66.8 |
| 132 | -81.2 | -51.8 | 4.2 | -1.9 | -28.8 | -3.2 | 67.5 | -67.3 |
| 133 | -75.4 | -36.2 | 3.2 | -2.0 | -27.2 | -5.0 | 52.7 | -60.8 |
| 134 | -84.0 | -38.0 | 3.1 | -2.0 | -29.7 | -3.2 | 57.3 | -71.5 |
| 135 | -94.3 | -70.7 | 2.2 | -2.7 | -30.7 | -5.5 | 76.3 | -63.3 |
| 136 | -79.0 | -56.6 | 4.8 | -1.9 | -31.3 | -2.5 | 77.1 | -68.6 |
| 137 | -91.4 | -38.6 | 1.6 | -2.2 | -31.0 | -3.8 | 51.5 | -68.8 |
| 138 | -83.1 | -48.0 | 1.2 | -2.2 | -28.7 | -3.7 | 64.3 | -66.0 |
| 139 | -80.6 | -48.3 | 1.2 | -1.7 | -28.9 | -2.7 | 67.7 | -67.9 |
| 140 | -89.6 | -49.8 | 2.7 | -2.1 | -28.9 | -4.8 | 60.6 | -67.3 |
| 141 | -88.7 | -40.5 | 1.3 | -2.4 | -27.3 | -1.7 | 51.5 | -69.6 |
| 142 | -88.2 | -52.2 | 1.9 | -2.2 | -27.0 | -2.7 | 64.2 | -70.1 |
| 143 | -77.9 | -39.4 | 1.9 | -2.5 | -25.0 | -1.9 | 55.2 | -66.3 |
| 144 | -91.9 | -42.8 | 2.6 | -2.9 | -29.1 | -2.4 | 52.6 | -69.9 |
| 145 | -96.2 | -41.4 | 2.9 | -2.3 | -29.9 | -2.4 | 53.9 | -77.0 |
| 146 | -90.6 | -40.5 | 2.3 | -2.6 | -27.8 | -1.6 | 47.9 | -68.4 |
| 147 | -92.7 | -41.0 | 2.4 | -2.6 | -27.7 | -3.4 | 52.5 | -72.8 |
| 148 | -95.5 | -54.3 | 2.1 | -2.4 | -30.4 | -1.1 | 61.6 | -71.1 |
| 149 | -86.9 | -54.1 | 2.9 | -2.5 | -25.8 | -3.0 | 63.2 | -67.6 |
| 150 | -96.3 | -38.4 | 4.5 | -2.4 | -29.3 | -2.7 | 47.6 | -75.5 |
| 151 | -84.9 | -53.6 | 4.3 | -3.9 | -26.9 | -1.4 | 64.5 | -67.9 |
| 152 | -84.7 | -41.0 | 3.7 | -2.5 | -26.6 | -3.4 | 48.1 | -63.0 |
| 153 | -103.3 | -38.3 | 0.2 | -3.4 | -29.2 | -4.8 | 38.6 | -66.4 |
| 154 | -94.4 | -49.8 | 1.7 | -3.2 | -26.2 | -2.5 | 54.4 | -68.8 |
| 155 | -96.0 | -38.3 | 5.0 | -3.1 | -28.2 | -5.6 | 46.6 | -72.3 |
| 156 | -91.7 | -43.5 | 4.5 | -2.4 | -29.2 | -3.0 | 55.4 | -73.4 |
| 157 | -96.6 | -46.4 | 2.8 | -2.3 | -29.5 | -5.3 | 56.6 | -72.6 |
| 158 | -99.2 | -36.7 | 1.2 | -1.9 | -31.2 | -4.5 | 48.4 | -74.6 |
| 159 | -93.8 | -47.8 | 4.4 | -3.2 | -29.2 | -2.8 | 58.0 | -73.1 |
| 160 | -90.7 | -39.3 | 1.9 | -2.5 | -27.9 | -2.7 | 49.7 | -69.9 |
| 161 | -93.3 | -35.7 | 3.3 | -2.4 | -31.5 | -2.9 | 50.6 | -74.7 |
| 162 | -89.4 | -38.9 | 2.7 | -2.5 | -28.1 | -2.9 | 52.3 | -72.2 |
| 163 | -94.4 | -64.3 | 4.1 | -2.6 | -28.4 | -3.7 | 70.9 | -70.3 |
| 164 | -95.0 | -25.7 | 3.6 | -2.9 | -27.9 | -2.5 | 33.6 | -73.3 |
| 165 | -94.1 | -50.6 | 2.4 | -2.5 | -29.2 | -2.7 | 59.8 | -71.3 |
| 166 | -88.5 | -41.1 | 6.8 | -1.8 | -29.2 | -3.4 | 49.4 | -69.1 |
| 167 | -85.0 | -45.7 | 6.4 | -3.2 | -27.5 | -3.7 | 54.5 | -65.9 |
| 168 | -93.1 | -36.7 | 2.9 | -3.3 | -26.7 | -2.0 | 46.0 | -73.2 |
| 169 | -93.3 | -51.5 | 5.6 | -3.5 | -30.2 | -3.3 | 66.7 | -77.0 |
| 170 | -91.3 | -40.1 | 4.1 | -3.1 | -31.0 | -2.8 | 53.5 | -71.9 |
| 171 | -91.7 | -53.3 | 6.1 | -3.0 | -28.9 | -3.0 | 63.3 | -72.9 |
| 172 | -95.6 | -46.5 | 3.0 | -2.7 | -30.7 | -2.7 | 54.3 | -70.2 |
| 173 | -91.1 | -44.1 | 5.0 | -2.5 | -30.3 | -5.5 | 56.6 | -70.3 |
| 174 | -83.1 | -48.2 | 6.0 | -2.4 | -28.0 | -1.7 | 60.2 | -69.0 |
| 175 | -78.4 | -49.5 | 3.6 | -2.0 | -26.9 | -2.1 | 61.8 | -63.3 |
| 176 | -86.9 | -60.5 | 4.4 | -2.4 | -29.2 | -3.1 | 71.8 | -68.1 |
| 177 | -81.8 | -45.1 | 3.8 | -1.8 | -28.5 | -3.5 | 59.2 | -65.9 |
| 178 | -87.8 | -45.9 | 4.2 | -2.3 | -31.1 | -4.4 | 57.5 | -65.8 |
| 179 | -90.9 | -57.4 | 3.4 | -2.9 | -29.7 | -2.5 | 62.3 | -64.1 |
| 180 | -76.3 | -53.1 | 1.6 | -2.6 | -24.6 | -4.2 | 69.3 | -62.8 |
| 181 | -86.4 | -49.6 | 5.1 | -2.7 | -30.7 | -5.8 | 61.8 | -64.5 |
| 182 | -94.3 | -31.0 | 1.9 | -2.2 | -32.1 | -5.2 | 41.9 | -67.6 |
| 183 | -82.0 | -30.0 | 3.0 | -2.0 | -27.7 | -4.6 | 46.2 | -66.8 |
| 184 | -95.1 | -36.8 | 1.7 | -2.5 | -30.8 | -5.7 | 50.4 | -71.3 |
| 185 | -90.9 | -39.0 | 4.9 | -2.8 | -29.5 | -2.8 | 46.6 | -68.3 |
| 186 | -89.6 | -40.8 | 1.3 | -2.0 | -29.7 | -4.1 | 51.4 | -65.5 |
| 187 | -88.2 | -45.7 | 5.0 | -3.1 | -32.5 | -3.9 | 57.2 | -65.3 |
| 188 | -88.7 | -42.5 | 2.3 | -2.2 | -30.7 | -4.5 | 55.1 | -66.2 |
| 189 | -85.8 | -51.2 | 1.6 | -2.0 | -31.4 | -2.9 | 66.5 | -66.3 |
| 190 | -90.4 | -36.5 | 3.5 | -2.4 | -30.0 | -3.0 | 48.3 | -70.1 |
| 191 | -89.8 | -36.3 | 3.2 | -2.1 | -30.2 | -2.6 | 46.1 | -67.8 |
| 192 | -93.0 | -58.7 | 3.8 | -2.1 | -30.8 | -3.7 | 67.8 | -69.3 |
| 193 | -78.5 | -50.3 | 2.9 | -2.9 | -25.4 | -3.7 | 62.9 | -62.0 |
| 194 | -77.2 | -40.9 | 3.6 | -2.1 | -25.3 | -2.1 | 55.7 | -66.2 |
| 195 | -82.8 | -52.1 | 3.5 | -2.6 | -26.9 | -4.1 | 63.2 | -63.7 |
| 196 | -79.6 | -57.4 | 3.4 | -2.9 | -27.3 | -2.7 | 71.2 | -63.9 |
| 197 | -90.3 | -34.3 | 3.5 | -2.2 | -30.7 | -2.5 | 48.3 | -72.4 |
| 198 | -81.2 | -45.5 | 2.8 | -2.1 | -26.7 | -3.2 | 57.7 | -64.2 |
| 199 | -92.3 | -59.5 | 2.7 | -2.4 | -31.0 | -2.2 | 68.7 | -68.6 |
| 200 | -86.6 | -53.2 | 1.9 | -1.8 | -30.3 | -2.6 | 67.2 | -67.7 |
| 201 | -87.3 | -44.2 | 2.8 | -2.2 | -31.2 | -4.8 | 56.0 | -63.7 |
| 202 | -87.2 | -42.8 | 4.1 | -2.3 | -28.3 | -1.3 | 53.3 | -69.8 |
| 203 | -80.3 | -53.9 | 6.4 | -2.4 | -28.7 | -1.5 | 68.8 | -69.0 |
| 204 | -87.3 | -56.2 | 3.2 | -1.7 | -31.1 | -5.8 | 63.8 | -59.5 |
| 205 | -89.8 | -49.5 | 3.8 | -2.2 | -32.2 | -2.9 | 61.4 | -68.2 |
| 206 | -87.9 | -38.8 | 2.7 | -2.2 | -30.6 | -5.2 | 56.1 | -69.8 |
| 207 | -76.8 | -54.2 | 1.9 | -1.8 | -29.0 | -4.0 | 72.2 | -61.8 |
| 208 | -84.5 | -45.1 | 0.7 | -2.5 | -28.0 | -5.3 | 56.6 | -60.9 |
| 209 | -89.6 | -53.3 | 3.5 | -2.2 | -28.9 | -5.9 | 63.9 | -66.8 |
| 210 | -80.2 | -61.2 | 4.1 | -1.9 | -28.5 | -4.8 | 77.4 | -65.2 |
| 211 | -85.9 | -31.2 | 2.7 | -2.3 | -31.1 | -4.8 | 46.4 | -65.6 |
| 212 | -75.0 | -47.5 | 3.6 | -2.2 | -26.5 | -3.0 | 62.2 | -61.7 |
| 213 | -77.6 | -36.6 | 2.9 | -2.4 | -27.8 | -3.6 | 52.0 | -62.2 |
| 214 | -76.3 | -34.0 | 8.0 | -2.3 | -28.2 | -4.2 | 49.3 | -64.9 |
| 215 | -81.2 | -46.3 | 2.4 | -2.1 | -28.1 | -2.8 | 60.2 | -64.5 |
| 216 | -86.6 | -45.0 | 2.5 | -2.1 | -31.1 | -4.2 | 57.1 | -63.7 |
| 217 | -79.1 | -59.2 | 2.2 | -1.9 | -27.8 | -3.6 | 73.8 | -62.5 |
| 218 | -83.7 | -44.8 | 2.7 | -2.0 | -28.7 | -2.8 | 58.1 | -66.2 |
| 219 | -82.3 | -53.4 | 3.1 | -1.9 | -31.5 | -4.7 | 70.9 | -64.9 |
| 220 | -84.5 | -53.3 | 7.3 | -2.8 | -32.3 | -4.7 | 67.2 | -65.9 |
| 221 | -86.5 | -57.2 | 1.6 | -2.2 | -32.7 | -4.1 | 71.4 | -63.3 |
| 222 | -83.6 | -47.4 | 1.9 | -1.7 | -31.2 | -3.3 | 61.6 | -63.4 |
| 223 | -79.7 | -55.5 | 9.9 | -2.2 | -27.8 | -5.4 | 69.5 | -68.1 |
| 224 | -75.8 | -56.2 | 7.7 | -2.2 | -28.6 | -2.4 | 72.6 | -66.6 |
| 225 | -81.8 | -24.3 | 2.4 | -1.9 | -30.4 | -2.3 | 42.0 | -67.5 |
| 226 | -76.5 | -57.6 | 3.2 | -2.7 | -27.3 | -4.1 | 74.0 | -62.1 |
| 227 | -79.2 | -67.5 | 5.3 | -2.8 | -26.9 | -4.9 | 81.3 | -63.8 |
| 228 | -82.4 | -29.9 | 1.3 | -1.9 | -28.0 | -3.9 | 44.9 | -64.9 |
| 229 | -72.0 | -41.1 | 2.3 | -2.1 | -24.6 | -1.7 | 55.5 | -60.4 |
| 230 | -81.3 | -42.2 | 6.0 | -1.8 | -30.2 | -3.1 | 59.5 | -69.5 |
| 231 | -87.2 | -29.1 | 5.8 | -2.1 | -34.9 | -4.6 | 46.7 | -69.1 |
| 232 | -93.6 | -43.6 | 3.8 | -2.1 | -33.6 | -3.7 | 58.0 | -72.3 |
| 233 | -89.6 | -46.1 | 10.3 | -2.1 | -33.0 | -3.7 | 61.8 | -76.7 |
| 234 | -80.0 | -27.6 | 2.3 | -2.1 | -27.7 | -3.1 | 43.9 | -65.8 |
| 235 | -89.2 | -40.9 | 5.1 | -2.7 | -32.0 | -2.9 | 56.3 | -72.1 |
| 236 | -83.7 | -55.2 | 2.6 | -2.7 | -31.2 | -2.4 | 73.7 | -68.6 |
| 237 | -75.6 | -46.2 | 7.4 | -2.7 | -28.8 | -2.8 | 66.2 | -68.8 |
| 238 | -72.1 | -23.9 | 3.0 | -1.7 | -24.1 | -4.2 | 43.5 | -64.8 |
| 239 | -74.2 | -60.9 | 1.7 | -1.6 | -30.2 | -2.5 | 84.3 | -65.1 |
| 240 | -88.8 | -56.5 | 7.0 | -3.0 | -31.2 | -3.7 | 68.2 | -69.5 |
| 241 | -80.9 | -44.5 | 5.6 | -2.0 | -30.5 | -3.3 | 60.0 | -66.3 |
| 242 | -94.5 | -56.6 | 3.9 | -2.3 | -31.1 | -3.1 | 67.0 | -72.3 |
| 243 | -91.1 | -46.5 | 1.6 | -2.1 | -29.2 | -3.4 | 57.1 | -68.5 |
| 244 | -85.2 | -44.7 | 0.8 | -2.1 | -26.8 | -3.4 | 58.7 | -67.8 |
| 245 | -93.4 | -42.8 | 2.9 | -2.0 | -29.1 | -3.2 | 51.0 | -70.2 |
| 246 | -85.2 | -41.1 | 0.8 | -2.1 | -27.3 | -3.5 | 52.5 | -64.5 |
| 247 | -79.8 | -50.4 | 2.4 | -2.2 | -28.6 | -2.5 | 66.5 | -65.0 |
| 248 | -83.4 | -47.7 | 2.6 | -2.5 | -29.1 | -2.4 | 59.6 | -64.0 |
| 249 | -82.9 | -52.8 | 3.4 | -2.2 | -30.0 | -5.1 | 66.3 | -62.6 |
| 250 | -75.2 | -33.7 | 5.3 | -2.2 | -26.8 | -3.2 | 49.5 | -64.0 |
| 251 | -69.4 | -45.7 | 6.8 | -2.1 | -26.4 | -2.7 | 66.2 | -65.5 |
| 252 | -86.0 | -53.4 | 1.5 | -2.1 | -28.8 | -5.1 | 67.3 | -65.4 |
| 253 | -85.3 | -47.6 | 0.8 | -2.0 | -29.1 | -4.8 | 64.5 | -67.1 |
| 254 | -87.2 | -39.2 | 2.4 | -2.1 | -30.8 | -2.0 | 53.9 | -69.4 |
| 255 | -88.6 | -45.6 | 1.0 | -2.2 | -31.8 | -5.0 | 61.5 | -66.6 |
| 256 | -78.7 | -44.7 | 1.0 | -2.1 | -28.5 | -2.1 | 61.4 | -63.8 |
| 257 | -78.6 | -58.6 | 2.0 | -1.9 | -27.3 | -2.2 | 71.7 | -62.3 |
| 258 | -84.0 | -51.6 | 2.1 | -2.2 | -31.8 | -4.1 | 70.3 | -66.8 |
| 259 | -76.2 | -50.3 | 4.5 | -2.2 | -25.8 | -2.0 | 61.8 | -62.2 |
| 260 | -80.9 | -44.4 | 1.7 | -2.1 | -30.7 | -3.0 | 60.6 | -63.1 |
| 261 | -82.5 | -41.3 | 2.8 | -1.7 | -29.9 | -2.1 | 58.7 | -69.0 |
| 262 | -81.3 | -41.4 | 5.1 | -2.6 | -30.3 | -3.0 | 54.3 | -63.4 |
| 263 | -91.5 | -45.5 | 3.1 | -2.3 | -32.5 | -5.5 | 58.7 | -67.4 |
| 264 | -88.1 | -49.2 | 5.6 | -2.2 | -32.8 | -3.9 | 63.2 | -68.7 |
| 265 | -86.5 | -60.0 | 3.7 | -1.9 | -33.4 | -3.7 | 75.3 | -66.5 |
| 266 | -74.1 | -50.7 | 2.4 | -1.9 | -25.7 | -2.1 | 66.0 | -62.2 |
| 267 | -88.0 | -47.7 | 1.9 | -2.3 | -32.2 | -4.3 | 64.6 | -68.0 |
| 268 | -85.7 | -52.3 | 4.0 | -2.3 | -31.8 | -1.8 | 64.0 | -65.6 |
| 269 | -92.6 | -59.1 | 1.9 | -2.0 | -33.3 | -5.8 | 76.5 | -70.8 |
| 270 | -86.7 | -59.7 | 5.2 | -2.3 | -30.9 | -3.7 | 71.5 | -66.7 |
| 271 | -73.6 | -39.1 | 1.6 | -0.9 | -26.1 | -2.2 | 57.2 | -64.1 |
| 272 | -86.2 | -55.1 | 1.0 | -2.3 | -28.9 | -2.5 | 64.6 | -63.1 |
| 273 | -79.4 | -30.5 | 2.3 | -2.0 | -28.9 | -3.2 | 50.4 | -67.5 |
| 274 | -84.5 | -52.0 | 3.3 | -2.6 | -28.7 | -2.3 | 67.2 | -69.5 |
| 275 | -93.5 | -55.1 | 5.1 | -2.5 | -32.2 | -4.0 | 67.9 | -72.6 |
| 276 | -90.4 | -50.6 | 3.2 | -2.6 | -31.9 | -2.9 | 66.6 | -72.1 |
| 277 | -90.7 | -49.1 | 4.6 | -2.7 | -31.9 | -2.4 | 61.8 | -71.2 |
| 278 | -85.9 | -49.5 | 6.6 | -2.6 | -30.0 | -3.7 | 61.5 | -68.1 |
| 279 | -82.2 | -42.5 | 3.2 | -2.7 | -29.3 | -6.1 | 60.5 | -65.2 |
| 280 | -94.6 | -32.4 | 0.9 | -2.3 | -32.4 | -2.8 | 46.3 | -71.9 |
| 281 | -81.9 | -43.7 | 2.9 | -2.1 | -27.3 | -4.4 | 56.3 | -63.5 |
| 282 | -88.7 | -57.0 | 1.0 | -2.2 | -27.8 | -4.1 | 65.3 | -64.0 |
| 283 | -75.2 | -40.6 | 3.2 | -2.0 | -27.3 | -2.7 | 57.4 | -63.1 |
| 284 | -77.3 | -63.4 | 1.8 | -2.3 | -26.6 | -2.0 | 75.4 | -60.2 |
| 285 | -80.6 | -37.7 | 2.8 | -1.9 | -30.3 | -3.6 | 56.4 | -66.3 |
| 286 | -77.7 | -57.2 | 1.3 | -2.4 | -26.4 | -3.2 | 71.4 | -61.3 |
| 287 | -78.0 | -49.5 | 3.3 | -2.3 | -28.6 | -2.8 | 65.7 | -63.8 |
| 288 | -81.4 | -32.0 | 1.2 | -1.7 | -28.1 | -2.0 | 46.7 | -65.5 |
| 289 | -80.1 | -40.1 | 1.5 | -2.1 | -29.5 | -4.4 | 57.1 | -62.5 |
| 290 | -84.0 | -43.4 | 2.9 | -1.9 | -28.7 | -2.5 | 55.3 | -65.9 |
| 291 | -81.3 | -53.1 | 3.0 | -2.2 | -29.2 | -2.0 | 65.0 | -62.7 |
| 292 | -77.1 | -55.2 | 5.8 | -2.4 | -28.2 | -2.4 | 69.3 | -64.0 |
| 293 | -80.1 | -47.5 | 2.9 | -2.0 | -28.6 | -1.5 | 61.3 | -64.7 |
| 294 | -88.5 | -56.4 | 4.7 | -1.6 | -28.7 | -4.7 | 63.8 | -65.6 |
| 295 | -78.4 | -52.7 | 2.8 | -1.9 | -24.3 | -1.8 | 59.4 | -59.9 |
| 296 | -82.9 | -50.5 | 5.5 | -1.2 | -30.2 | -3.0 | 65.4 | -69.0 |
| 297 | -79.5 | -36.2 | 3.9 | -1.2 | -27.9 | -4.6 | 48.8 | -62.3 |
| 298 | -81.7 | -28.4 | 4.7 | -1.3 | -28.6 | -3.3 | 40.6 | -65.4 |
| 299 | -81.5 | -40.9 | 3.9 | -2.2 | -26.4 | -3.0 | 53.1 | -66.0 |
| 300 | -87.1 | -54.2 | 8.1 | -2.6 | -32.3 | -4.0 | 64.3 | -66.4 |
| 301 | -88.5 | -59.9 | 1.8 | -2.1 | -28.0 | -3.5 | 66.9 | -63.9 |
| 302 | -85.9 | -41.6 | 4.3 | -1.8 | -28.6 | -3.0 | 50.5 | -65.7 |
| 303 | -77.9 | -41.7 | 3.8 | -1.6 | -27.1 | -3.4 | 54.1 | -62.0 |
| 304 | -80.1 | -44.2 | 1.9 | -2.1 | -26.7 | -1.8 | 57.9 | -65.2 |
| 305 | -72.9 | -47.0 | 4.0 | -1.6 | -26.4 | -2.9 | 64.4 | -63.4 |
| 306 | -88.7 | -50.5 | 3.2 | -2.1 | -33.0 | -3.4 | 65.7 | -68.5 |
| 307 | -81.6 | -42.2 | 3.4 | -2.0 | -30.1 | -2.7 | 56.9 | -64.8 |
| 308 | -80.7 | -35.1 | 3.6 | -2.1 | -28.8 | -2.5 | 50.5 | -66.4 |
| 309 | -78.4 | -45.5 | 4.8 | -1.9 | -28.1 | -1.4 | 59.9 | -66.3 |
| 310 | -89.3 | -44.9 | 4.6 | -2.3 | -31.1 | -5.2 | 59.7 | -70.2 |
| 311 | -88.7 | -46.9 | 3.3 | -1.2 | -30.7 | -3.9 | 57.9 | -67.2 |
| 312 | -81.3 | -43.3 | 2.9 | -2.2 | -28.7 | -5.2 | 60.7 | -65.5 |
| 313 | -83.6 | -45.1 | 2.1 | -2.1 | -30.9 | -2.1 | 62.2 | -67.7 |
| 314 | -79.4 | -51.0 | 4.2 | -2.0 | -28.6 | -2.9 | 64.9 | -64.0 |
| 315 | -88.6 | -53.1 | 2.6 | -2.0 | -31.3 | -2.7 | 65.1 | -67.2 |
| 316 | -80.6 | -37.9 | 4.2 | -2.0 | -31.0 | -0.9 | 61.4 | -74.5 |
| 317 | -77.1 | -42.5 | 2.3 | -1.8 | -29.3 | -2.7 | 61.9 | -64.9 |
| 318 | -85.7 | -46.3 | 1.7 | -1.9 | -30.5 | -3.2 | 60.4 | -65.9 |
| 319 | -78.3 | -41.7 | 3.4 | -1.6 | -28.7 | -2.5 | 59.9 | -67.1 |
| 320 | -87.9 | -54.2 | 4.7 | -1.9 | -30.5 | -3.1 | 65.4 | -68.3 |
| 321 | -85.4 | -41.6 | 5.4 | -2.5 | -27.3 | -4.6 | 51.6 | -66.5 |
| 322 | -89.1 | -31.2 | 6.4 | -2.9 | -31.6 | -5.8 | 43.1 | -67.1 |
| 323 | -86.6 | -43.7 | 3.3 | -2.4 | -32.2 | -2.8 | 58.3 | -67.1 |
| 324 | -83.1 | -34.9 | 3.1 | -2.4 | -29.4 | -2.3 | 48.0 | -65.1 |
| 325 | -86.5 | -58.0 | 2.2 | -2.2 | -28.9 | -2.7 | 69.9 | -66.9 |
| 326 | -90.4 | -64.5 | 1.8 | -2.6 | -29.3 | -2.8 | 73.7 | -66.7 |
| 327 | -85.9 | -63.0 | 3.9 | -2.4 | -29.9 | -4.9 | 73.6 | -63.2 |
| 328 | -89.9 | -48.3 | 4.6 | -2.6 | -32.7 | -3.8 | 60.8 | -67.9 |
| 329 | -84.3 | -38.3 | 3.5 | -2.4 | -29.4 | -3.6 | 54.6 | -68.7 |
| 330 | -91.7 | -54.5 | 3.0 | -3.2 | -31.3 | -3.4 | 67.0 | -69.3 |
| 331 | -92.0 | -35.3 | 3.1 | -2.9 | -32.9 | -3.0 | 47.6 | -68.7 |
| 332 | -81.7 | -43.3 | 3.6 | -2.6 | -29.9 | -5.5 | 61.4 | -65.3 |
| 333 | -88.0 | -50.2 | 3.6 | -2.5 | -28.9 | -3.0 | 61.1 | -68.1 |
| 334 | -79.7 | -69.0 | 5.9 | -2.1 | -25.8 | -4.1 | 80.0 | -64.4 |
| 335 | -84.3 | -49.6 | 1.9 | -2.4 | -30.8 | -3.4 | 64.8 | -64.8 |
| 336 | -79.0 | -53.3 | 4.4 | -2.1 | -28.5 | -4.0 | 69.6 | -65.1 |
| 337 | -82.4 | -66.1 | 1.5 | -2.1 | -28.3 | -4.1 | 80.2 | -63.6 |
| 338 | -77.1 | -44.7 | 4.3 | -2.2 | -26.9 | -4.3 | 59.1 | -62.4 |
| 339 | -84.8 | -39.6 | 2.1 | -1.9 | -28.2 | -3.6 | 48.9 | -62.5 |
| 340 | -81.5 | -49.7 | 4.9 | -2.5 | -27.4 | -4.5 | 60.2 | -62.5 |
| 341 | -82.7 | -48.2 | 2.2 | -2.2 | -27.5 | -4.4 | 59.5 | -61.9 |
| 342 | -84.2 | -31.9 | 1.9 | -2.1 | -28.8 | -4.0 | 43.8 | -63.1 |
| 343 | -75.2 | -40.1 | 4.4 | -1.9 | -27.1 | -2.7 | 54.1 | -61.7 |
| 344 | -77.5 | -44.0 | 1.5 | -1.9 | -28.1 | -2.2 | 56.4 | -59.2 |
| 345 | -82.6 | -40.0 | 2.4 | -1.1 | -28.2 | -3.4 | 50.7 | -63.0 |
| 346 | -80.4 | -63.0 | 2.8 | -2.1 | -26.1 | -2.3 | 72.8 | -62.5 |
| 347 | -78.7 | -50.4 | 4.0 | -1.9 | -29.1 | -3.0 | 64.6 | -62.9 |
| 348 | -82.7 | -31.4 | 2.5 | -2.2 | -29.0 | -3.6 | 43.6 | -62.7 |
| 349 | -85.1 | -62.9 | 3.7 | -2.1 | -27.9 | -2.3 | 71.1 | -64.7 |
| 350 | -79.4 | -51.2 | 1.6 | -2.0 | -27.7 | -2.2 | 65.6 | -63.4 |
| 351 | -72.9 | -43.4 | 3.3 | -2.1 | -21.9 | -3.0 | 53.4 | -59.2 |
| 352 | -74.4 | -38.4 | 0.7 | -1.9 | -24.7 | -2.6 | 53.0 | -60.6 |
| 353 | -76.7 | -56.2 | 0.8 | -1.9 | -26.3 | -1.8 | 69.2 | -60.6 |
| 354 | -77.7 | -62.0 | 2.8 | -2.0 | -28.7 | -2.2 | 78.8 | -64.5 |
| 355 | -82.0 | -40.2 | 4.5 | -2.2 | -30.1 | -3.4 | 51.3 | -61.9 |
| 356 | -85.7 | -46.6 | 2.6 | -2.0 | -28.3 | -4.3 | 56.9 | -64.0 |
| 357 | -70.6 | -38.4 | 2.8 | -1.7 | -27.1 | -2.6 | 56.8 | -60.3 |
| 358 | -86.1 | -58.5 | 2.5 | -2.0 | -29.6 | -2.8 | 71.6 | -67.3 |
| 359 | -80.6 | -57.6 | 7.6 | -2.3 | -28.8 | -1.8 | 69.7 | -67.3 |
| 360 | -85.0 | -47.0 | 2.1 | -2.3 | -29.5 | -2.3 | 57.5 | -63.5 |
| 361 | -83.0 | -64.8 | 1.7 | -1.9 | -28.5 | -1.6 | 78.8 | -66.7 |
| 362 | -84.0 | -48.0 | 3.2 | -1.7 | -29.8 | -2.6 | 63.8 | -68.8 |
| 363 | -81.4 | -49.1 | 2.2 | -2.1 | -30.1 | -3.0 | 64.4 | -63.5 |
| 364 | -80.6 | -38.0 | 2.1 | -2.3 | -27.2 | -3.3 | 48.0 | -60.0 |
| 365 | -78.5 | -40.5 | 4.2 | -2.5 | -25.1 | -2.3 | 48.7 | -60.8 |
| 366 | -79.4 | -34.1 | -1.0 | -1.8 | -27.1 | -2.7 | 47.9 | -60.6 |
| 367 | -88.8 | -39.6 | 3.0 | -2.2 | -32.4 | -2.8 | 51.6 | -66.5 |
| 368 | -84.5 | -29.0 | 1.7 | -1.9 | -30.5 | -2.6 | 40.2 | -62.3 |
| 369 | -76.0 | -67.3 | 7.7 | -2.0 | -28.2 | -1.6 | 79.0 | -63.6 |
| 370 | -71.1 | -48.5 | 3.5 | -2.1 | -27.6 | -4.2 | 69.2 | -61.5 |
| 371 | -71.7 | -36.4 | 2.8 | -2.0 | -26.1 | -3.2 | 57.4 | -64.1 |
| 372 | -80.6 | -41.1 | 5.4 | -2.0 | -27.9 | -4.5 | 55.8 | -66.3 |
| 373 | -78.4 | -40.8 | 6.4 | -2.6 | -30.3 | -3.5 | 57.5 | -65.1 |
| 374 | -75.4 | -32.5 | 2.2 | -2.3 | -25.3 | -1.5 | 46.8 | -62.8 |
| 375 | -76.9 | -32.9 | 2.7 | -1.9 | -28.5 | -2.7 | 47.9 | -61.4 |
| 376 | -75.3 | -42.0 | 4.1 | -1.6 | -28.5 | -1.9 | 60.5 | -65.8 |
| 377 | -89.8 | -48.9 | 3.0 | -2.0 | -32.2 | -1.8 | 62.5 | -70.4 |
| 378 | -79.0 | -39.3 | 3.2 | -2.1 | -28.4 | -1.8 | 56.7 | -67.4 |
| 379 | -88.4 | -53.6 | 2.3 | -2.2 | -31.2 | -1.8 | 66.4 | -68.2 |
| 380 | -83.5 | -53.2 | 3.8 | -2.2 | -29.3 | -2.8 | 69.3 | -69.1 |
| 381 | -85.9 | -49.4 | 1.7 | -2.4 | -30.0 | -2.0 | 62.6 | -66.3 |
| 382 | -78.5 | -41.8 | 3.8 | -1.9 | -28.7 | -2.2 | 63.5 | -71.1 |
| 383 | -72.0 | -23.5 | 4.1 | -1.2 | -27.4 | -3.0 | 38.2 | -59.3 |
| 384 | -69.3 | -35.1 | 1.7 | -1.9 | -27.3 | -1.4 | 57.3 | -62.5 |
| 385 | -79.5 | -63.3 | 2.7 | -2.3 | -28.0 | -3.9 | 79.2 | -63.9 |
| 386 | -68.4 | -56.7 | 3.8 | -2.1 | -26.2 | -3.0 | 77.2 | -61.4 |
| 387 | -77.1 | -36.1 | 5.9 | -2.3 | -28.3 | -5.6 | 53.0 | -63.7 |
| 388 | -81.9 | -42.8 | 3.2 | -2.4 | -30.1 | -2.7 | 55.9 | -62.9 |
| 389 | -84.6 | -57.7 | 1.4 | -2.1 | -29.9 | -2.5 | 71.7 | -65.3 |
| 390 | -83.4 | -49.7 | 2.7 | -2.1 | -31.5 | -1.4 | 66.1 | -67.5 |
| 391 | -89.2 | -53.6 | 2.3 | -2.2 | -32.6 | -2.2 | 67.0 | -67.9 |
| 392 | -86.0 | -57.2 | 2.7 | -2.3 | -31.0 | -2.6 | 67.9 | -63.5 |
| 393 | -93.5 | -49.5 | 1.9 | -2.1 | -34.3 | -2.7 | 61.6 | -68.5 |
| 394 | -78.0 | -42.8 | 0.9 | -2.2 | -27.8 | -1.6 | 59.7 | -64.2 |
| 395 | -85.3 | -36.2 | 2.1 | -2.1 | -30.0 | -2.0 | 52.0 | -69.2 |
| 396 | -77.9 | -49.6 | 2.8 | -2.3 | -28.8 | -2.7 | 68.8 | -66.1 |
| 397 | -87.5 | -47.5 | 4.6 | -2.3 | -31.9 | -2.8 | 61.6 | -69.1 |
| 398 | -84.5 | -55.2 | 3.4 | -2.2 | -30.5 | -2.2 | 66.5 | -64.3 |
| 399 | -76.2 | -47.2 | 2.1 | -1.9 | -27.2 | -2.1 | 63.7 | -63.7 |
| 400 | -83.1 | -58.5 | 1.6 | -2.1 | -29.2 | -2.5 | 72.6 | -64.9 |
| 401 | -80.1 | -54.2 | 3.5 | -1.8 | -29.5 | -3.3 | 71.8 | -66.6 |
| 402 | -83.5 | -48.3 | 4.3 | -2.3 | -30.7 | -2.8 | 62.5 | -66.2 |
| 403 | -81.8 | -36.3 | 4.5 | -2.4 | -28.1 | -1.6 | 47.5 | -65.4 |
| 404 | -80.3 | -35.1 | 4.3 | -2.4 | -28.8 | -3.0 | 53.9 | -69.2 |
| 405 | -91.4 | -42.7 | 4.2 | -2.3 | -32.2 | -2.6 | 55.0 | -70.7 |
| 406 | -81.9 | -23.6 | 2.0 | -2.1 | -29.6 | -2.1 | 35.2 | -61.7 |
| 407 | -75.2 | -57.4 | 1.4 | -2.4 | -24.5 | -1.8 | 72.3 | -62.9 |
| 408 | -82.5 | -40.8 | 3.8 | -2.0 | -29.3 | -3.5 | 53.5 | -64.2 |
| 409 | -76.5 | -45.3 | 3.1 | -2.0 | -26.4 | -2.3 | 60.4 | -64.0 |
| 410 | -88.3 | -30.8 | 2.9 | -1.9 | -31.9 | -2.3 | 44.3 | -68.6 |
| 411 | -77.1 | -55.4 | 3.5 | -2.2 | -25.8 | -1.6 | 64.9 | -60.6 |
| 412 | -90.7 | -54.9 | 3.1 | -2.3 | -31.5 | -4.1 | 65.1 | -66.1 |
| 413 | -80.3 | -51.1 | 4.9 | -2.2 | -28.7 | -2.6 | 67.5 | -68.1 |
| 414 | -71.7 | -37.2 | 3.5 | -1.8 | -25.9 | -2.5 | 53.8 | -61.6 |
| 415 | -78.8 | -43.1 | 2.2 | -1.9 | -27.1 | -2.5 | 54.4 | -60.9 |
| 416 | -85.2 | -34.7 | 2.0 | -2.1 | -28.5 | -5.2 | 51.7 | -68.4 |
| 417 | -77.4 | -52.9 | 4.7 | -2.3 | -28.1 | -2.8 | 69.3 | -65.3 |
| 418 | -80.9 | -40.4 | 6.4 | -2.1 | -30.3 | -5.3 | 56.6 | -65.9 |
| 419 | -85.0 | -47.2 | 3.4 | -2.2 | -28.4 | -5.5 | 58.7 | -63.7 |
| 420 | -82.1 | -39.5 | 2.8 | -2.0 | -30.5 | -1.8 | 54.8 | -65.8 |
| 421 | -82.3 | -53.2 | 1.2 | -2.0 | -29.1 | -2.1 | 71.1 | -68.1 |
| 422 | -87.5 | -50.4 | 2.5 | -1.9 | -31.2 | -2.7 | 63.1 | -66.9 |
| 423 | -95.6 | -44.8 | 2.0 | -2.2 | -32.4 | -3.4 | 57.5 | -72.3 |
| 424 | -79.4 | -47.3 | 4.8 | -1.9 | -28.9 | -4.7 | 65.2 | -66.5 |
| 425 | -77.1 | -37.5 | 5.2 | -2.3 | -28.3 | -4.6 | 52.2 | -61.8 |
| 426 | -85.1 | -39.5 | 3.8 | -2.0 | -30.5 | -2.5 | 55.8 | -70.2 |
| 427 | -82.9 | -45.7 | 2.8 | -2.1 | -28.8 | -4.1 | 62.2 | -67.2 |
| 428 | -74.5 | -32.5 | 2.8 | -2.1 | -30.8 | -2.5 | 55.5 | -64.9 |
| 429 | -86.6 | -70.5 | 5.4 | -2.4 | -31.0 | -4.7 | 79.7 | -63.0 |
| 430 | -93.1 | -27.4 | 4.3 | -2.9 | -33.6 | -3.3 | 39.4 | -69.6 |
| 431 | -86.7 | -16.9 | 3.2 | -2.3 | -30.7 | -3.3 | 31.3 | -68.1 |
| 432 | -88.2 | -48.1 | 3.4 | -2.5 | -33.4 | -2.5 | 64.4 | -69.5 |
| 433 | -78.6 | -54.3 | 1.9 | -2.1 | -28.5 | -2.6 | 72.0 | -65.2 |
| 434 | -84.3 | -48.9 | 2.8 | -2.2 | -31.8 | -3.0 | 63.7 | -64.7 |
| 435 | -90.8 | -49.1 | 2.3 | -2.5 | -30.4 | -2.4 | 60.0 | -68.7 |
| 436 | -71.4 | -47.2 | 1.0 | -1.6 | -25.6 | -0.9 | 66.7 | -63.8 |
| 437 | -85.3 | -60.0 | -0.3 | -2.1 | -28.5 | -2.4 | 67.9 | -59.9 |
| 438 | -82.4 | -49.5 | 2.1 | -1.8 | -28.3 | -3.9 | 62.3 | -63.4 |
| 439 | -85.0 | -51.2 | 2.7 | -1.9 | -29.1 | -3.7 | 64.1 | -65.9 |
| 440 | -81.1 | -49.1 | 2.6 | -2.3 | -30.7 | -4.9 | 68.4 | -65.1 |
| 441 | -84.0 | -61.5 | 5.1 | -2.5 | -30.8 | -5.4 | 74.8 | -63.6 |
| 442 | -87.9 | -47.2 | 2.6 | -2.1 | -29.2 | -3.0 | 59.0 | -68.0 |
| 443 | -80.3 | -39.3 | 3.9 | -2.9 | -30.1 | -4.0 | 57.8 | -65.7 |
| 444 | -81.8 | -56.3 | 3.0 | -2.4 | -29.0 | -3.4 | 69.5 | -63.2 |
| 445 | -88.2 | -52.9 | 2.3 | -2.4 | -30.6 | -2.8 | 65.9 | -67.7 |
| 446 | -84.5 | -39.2 | 3.8 | -2.4 | -28.8 | -2.4 | 53.3 | -68.8 |
| 447 | -90.7 | -34.6 | 1.9 | -2.0 | -32.6 | -2.5 | 49.5 | -70.3 |
| 448 | -84.6 | -29.4 | 2.6 | -2.1 | -30.3 | -2.3 | 43.8 | -66.8 |
| 449 | -85.4 | -40.1 | 4.4 | -2.0 | -30.0 | -2.9 | 55.3 | -70.1 |
| 450 | -85.9 | -43.4 | 3.7 | -2.4 | -30.7 | -2.6 | 54.3 | -64.8 |
| 451 | -87.7 | -35.0 | 3.7 | -2.3 | -30.8 | -3.7 | 50.3 | -69.8 |
| 452 | -86.4 | -53.9 | 2.8 | -1.9 | -31.9 | -2.6 | 68.3 | -67.1 |
| 453 | -83.3 | -53.6 | 2.8 | -2.8 | -29.0 | -2.0 | 68.2 | -66.8 |
| 454 | -77.0 | -40.5 | 8.4 | -2.5 | -30.1 | -1.8 | 60.5 | -71.1 |
| 455 | -74.4 | -38.4 | 9.2 | -2.5 | -29.2 | -2.3 | 58.1 | -69.5 |
| 456 | -89.3 | -40.1 | 3.2 | -2.4 | -31.8 | -3.0 | 54.3 | -69.6 |
| 457 | -81.8 | -43.6 | 4.7 | -3.2 | -27.5 | -1.5 | 53.3 | -64.2 |
| 458 | -83.3 | -31.3 | 2.1 | -2.0 | -31.3 | -2.5 | 51.9 | -70.3 |
| 459 | -84.3 | -34.9 | 5.3 | -2.0 | -32.3 | -3.9 | 44.9 | -61.4 |
| 460 | -80.5 | -45.6 | 3.6 | -2.2 | -28.1 | -2.9 | 60.8 | -65.9 |
| 461 | -85.5 | -26.0 | 3.3 | -2.4 | -28.5 | -2.1 | 37.3 | -67.1 |
| 462 | -84.6 | -53.2 | 3.0 | -2.2 | -28.6 | -2.8 | 65.8 | -66.6 |
| 463 | -82.8 | -39.9 | 2.4 | -1.9 | -27.4 | -2.6 | 52.5 | -65.9 |
| 464 | -86.2 | -51.8 | 3.4 | -2.4 | -30.0 | -2.3 | 62.2 | -65.3 |
| 465 | -82.0 | -49.4 | 4.5 | -2.3 | -28.2 | -2.0 | 59.7 | -64.2 |
| 466 | -85.0 | -43.4 | 3.9 | -2.3 | -30.7 | -2.7 | 54.0 | -63.7 |
| 467 | -90.8 | -60.1 | 2.4 | -2.2 | -31.9 | -2.8 | 74.7 | -70.9 |
| 468 | -86.5 | -37.3 | 2.2 | -2.4 | -29.4 | -1.8 | 52.0 | -69.7 |
| 469 | -94.0 | -43.5 | 2.1 | -2.4 | -32.5 | -2.8 | 55.7 | -70.6 |
| 470 | -85.5 | -42.3 | 1.6 | -2.1 | -30.8 | -1.6 | 61.2 | -71.5 |
| 471 | -80.0 | -44.3 | 5.4 | -2.2 | -27.3 | -1.0 | 55.8 | -66.3 |
| 472 | -90.3 | -50.9 | 1.3 | -2.2 | -30.6 | -3.1 | 64.5 | -69.4 |
| 473 | -89.8 | -52.7 | 1.9 | -2.1 | -29.6 | -3.2 | 65.3 | -69.3 |
| 474 | -92.0 | -53.7 | 3.8 | -2.6 | -34.1 | -2.4 | 64.8 | -67.9 |
| 475 | -95.7 | -59.7 | 4.5 | -2.8 | -32.6 | -1.8 | 72.4 | -75.6 |
| 476 | -81.7 | -55.2 | 4.9 | -3.3 | -29.1 | -2.5 | 71.1 | -67.6 |
| 477 | -87.5 | -40.7 | 2.3 | -2.1 | -29.6 | -2.4 | 52.4 | -67.3 |
| 478 | -87.4 | -57.6 | 3.1 | -2.0 | -30.9 | -3.0 | 68.3 | -65.3 |
| 479 | -85.4 | -55.1 | 2.3 | -1.9 | -29.4 | -2.8 | 69.1 | -67.7 |
| 480 | -90.1 | -48.4 | 6.1 | -2.8 | -31.7 | -2.7 | 57.9 | -68.5 |
| 481 | -93.0 | -36.6 | 4.0 | -1.5 | -31.9 | -6.8 | 47.3 | -67.6 |
| 482 | -87.7 | -64.1 | 4.0 | -2.3 | -31.6 | -4.1 | 78.4 | -68.0 |
| 483 | -87.0 | -40.4 | 3.6 | -2.2 | -31.6 | -2.7 | 53.4 | -67.3 |
| 484 | -85.5 | -61.0 | 2.6 | -2.6 | -31.4 | -2.8 | 74.8 | -65.2 |
| 485 | -89.8 | -51.7 | 2.1 | -2.2 | -31.8 | -3.1 | 65.4 | -68.6 |
| 486 | -87.4 | -60.9 | 7.6 | -2.2 | -31.2 | -2.9 | 72.1 | -69.9 |
| 487 | -80.7 | -52.5 | 2.7 | -2.0 | -28.0 | -3.2 | 64.3 | -61.9 |
| 488 | -82.5 | -54.4 | 3.7 | -2.5 | -26.8 | -2.9 | 62.0 | -61.6 |
| 489 | -83.0 | -33.1 | 0.2 | -2.4 | -26.3 | -2.5 | 46.3 | -65.2 |
| 490 | -93.5 | -47.4 | 0.5 | -2.3 | -29.6 | -2.5 | 58.3 | -70.5 |
| 491 | -95.0 | -51.8 | 2.1 | -2.5 | -31.0 | -3.9 | 65.3 | -73.2 |
| 492 | -86.5 | -50.5 | 1.8 | -2.4 | -28.6 | -3.1 | 61.6 | -65.3 |
| 493 | -90.2 | -74.5 | 3.3 | -3.0 | -31.2 | -2.2 | 84.2 | -66.8 |
| 494 | -92.2 | -35.0 | 4.0 | -2.5 | -31.6 | -2.2 | 49.2 | -74.2 |
| 495 | -94.5 | -49.7 | 3.6 | -2.2 | -31.4 | -3.2 | 59.0 | -70.6 |
| 496 | -82.2 | -53.6 | 4.7 | -2.2 | -30.1 | -2.9 | 67.6 | -65.6 |
| 497 | -94.2 | -56.4 | 3.0 | -3.0 | -31.8 | -2.9 | 68.8 | -72.0 |
| 498 | -96.1 | -50.8 | 2.8 | -3.1 | -33.1 | -3.0 | 63.3 | -72.2 |
| 499 | -94.5 | -52.2 | 0.5 | -1.9 | -31.2 | -3.2 | 65.3 | -71.8 |
| 500 | -83.5 | -57.1 | 2.0 | -2.3 | -30.5 | -3.8 | 73.1 | -65.0 |
| MAX | **-103.4** | **-76.7** | **-1.0** | **-3.9** | **-36.1** | **-6.8** | **31.3** | **-77.0** |
| MIN | **-67.5** | **-16.9** | **11.5** | **-0.9** | **-21.9** | **-0.6** | **84.3** | **-51.4** |
| AVG | **-83.9** | **-45.7** | **3.4** | **-2.2** | **-29.1** | **-3.1** | **58.9** | **-66.0** |
| SDV | **6.0** | **9.4** | **1.7** | **0.4** | **2.1** | **1.1** | **9.4** | **3.5** |
